# ΔNp63-restricted viral mimicry response impedes cancer cell viability and remodels tumor microenvironment in esophageal squamous cell carcinoma

**DOI:** 10.1101/2024.03.17.585449

**Authors:** Valen Zhuoyou Yu, Shan Shan So, Bryan Chee-chad Lung, George Zhaozheng Hou, Carissa Wing-yan Wong, Larry Ka-yue Chow, Michael King-yung Chung, Ian Yu-hong Wong, Claudia Lai-yin Wong, Desmond Kwan-kit Chan, Fion Siu-yin Chan, Betty Tsz-ting Law, Kaiyan Xu, Zack Zhen Tan, Ka-on Lam, Anthony Wing-ip Lo, Alfred King-yin Lam, Dora Lai-wan Kwong, Josephine Mun-yee Ko, Wei Dai, Simon Law, Maria Li Lung

**Affiliations:** Department of Clinical Oncology, Centre of Cancer Medicine, School of Clinical Medicine, Li Ka Shing Faculty of Medicine, The University of Hong Kong, Pokfulam, Hong Kong (SAR); Department of Surgery, School of Clinical Medicine, Li Ka Shing Faculty of Medicine, The University of Hong Kong, Pokfulam, Hong Kong (SAR); Division of Anatomical Pathology, Queen Mary Hospital, Pokfulam, Hong Kong (SAR); Divsion of Cancer Molecular Pathology, School of Medicine and Dentistry and Menzies Health Institute Queensland, Griffith University, Gold Coast, Australia

## Abstract

Tumor protein p63 isoform ΔNp63 plays roles in the squamous epithelium and squamous cell carcinomas (SCCs), including esophageal SCC (ESCC). By integrating data from cell lines and our latest patient-derived organoid cultures, derived xenograft models, and clinical sample transcriptomic analyses, we identified a novel and robust oncogenic role of ΔNp63 in ESCC. We showed that ΔNp63 maintains the repression of cancer cell endogenous retrotransposon expression and cellular double-stranded RNA sensing. These subsequently lead to a restricted cancer cell viral mimicry response and suppressed induction of tumor-suppressive type I interferon (IFN-I) signaling through the regulations of Signal transducer and activator of transcription 1, Interferon regulatory factor 1, and cGAS-STING pathway. The cancer cell ΔNp63-IFN-I signaling axis affects both the cancer cell and tumor-infiltrating immune cell (TIIC) compartments. In cancer cells, depletion of ΔNp63 resulted in reduced cell viability. ΔNp63 expression is negatively associated with the anticancer responses to viral mimicry booster treatments targeting cancer cells. In the tumor microenvironment, cancer cell *TP63* expression negatively correlates with multiple TIIC signatures in ESCC clinical samples. ΔNp63 depletion leads to increased cancer cell antigen presentation molecule expression and enhanced recruitment and reprogramming of tumor-infiltrating myeloid cells. Similar IFN-I signaling and TIIC signature association with ΔNp63 were also observed in lung SCC. These results support the potential application of ΔNp63 as a therapeutic target and a biomarker to guide candidate anticancer treatments exploring viral mimicry responses.

## introduction

Squamous cell carcinomas (SCCs) are among the most prevalent cancers. Esophageal squamous cell carcinoma (ESCC) is the predominant subtype of esophageal cancer in Asia and is one of the deadliest SCCs, with a dismal five-year survival rate of 10-20%. The molecular pathogenesis of ESCC is not fully understood. Thus, treatment options for ESCC are highly limited. An improved thorough understanding of the disease is essential for better disease management.

*Tumor protein p63* (*TP63*), encoding the transcription factor (TF) p63, plays a fundamental role in stratified epithelial biology. Two main p63 isoforms exist, including the full-length TAp63 and the truncated ΔNp63; the latter lacks the N-terminal transcription activation domain but retains TF activity. In the esophagus, ΔNp63 is the predominant isoform and is required for normal epithelial development with a basal layer-restricted expression pattern (1). ESCC retains the predominant expression of ΔNp63 with a relatively homogenous pattern observed in most cancer cells (2). ΔNp63 also plays critical oncogenic roles in several other SCCs (3).

Endogenous retrotransposons are actively involved in cancer biology. Developed cancer cells maintain a repressed retrotransposon expression. Derepressed elevated retrotransposon expression, either due to genetic predisposition or anticancer treatment, enhances cancer cell viral mimicry response, a tumor-suppressive cellular state of antiviral response triggered by endogenous stimuli. Expression of retrotransposon-encoded RNAs is prone to cytosolic double-stranded RNA (dsRNA) formation, which is recognized by sensor proteins and is capable of inducing anticancer responses, including type I interferon (IFN-I) signaling (4–7). The awakening of cancer cell IFN-I signaling triggers anticancer immune responses (8). The cancer cell retrotransposon expression has recently gained significant attention in the context of epigenetic therapy (8). Retrotransposon expression regulation serves as a cancer-specific therapeutic vulnerability to exploit for synergistic epigenetic therapies and immunotherapies (4,8).

In the present study, we scrutinized the influences of ΔNp63 expression in ESCC. By detailed functional and molecular characterizations, coupled with multi-method analyses of transcriptomic data, we identified a novel function of cancer cell ΔNp63 in restricting endogenous retrotransposon expression and suppressing tumor-suppressive IFN-I signaling. Cancer cell ΔNp63 expression exerts oncogenic effects on cancer cells and tumor-infiltrating immune cells (TIICs).

## MATERIALS AND METHODS

### Reagents and antibodies

Chemical reagents used in this study were purchased from MedChemExpress (Monmouth Junction, NJ) unless otherwise stated. Cell culture reagents were purchased from Thermo Fisher Scientific (Waltham, MA) unless otherwise stated. Details of primers and antibodies are in Supplementary Tables 1 and 2.

### Cell lines

Immortalized human esophageal epithelial cell lines, including NE1 (Research resource identifier:CVCL_E306), NE2 (9), and NE083 (10), and ESCC cell lines, including EC109 (CVCL_6898), KYSE30 (CVCL_1351)/KYSE30TSI, KYSE70 (CVCL_1356)/KYSE70TS, KYSE150 (CVCL_1348), KYSE180 (CVCL_1349)/KYSE180TS, KYSE450 (CVCL_1353) were cultured as described (11). KYSE30TSI, KYSE70TS, and KYSE180TS were established from parental nude mouse subcutaneous xenograft tumor segregants (11,12) and used for *in vivo* studies. L-WRN (CVCL_DA06) was acquired from AddexBio (San Diego, CA). The L-WRN conditioned medium (CM) containing Wnt-3a, R-spondin, and Noggin was produced as described (13) and used for PDO cultures. HEK293T (CVCL_0063) was used for lentiviral particle production. Cell line authentication by STR DNA profiling was performed for all cell lines. Mycoplasma test by PCR amplification of mycoplasma DNA was routinely performed for all *in vitro* cultures.

### Organoid culture establishment and maintenance

Patient tissue samples were obtained during endoscopic examinations at diagnosis (tumor tissue only), upfront surgery on patients without prior treatment (both tumor tissue and adjacent non-neoplastic tissue), and surgery on patients following neoadjuvant chemoradiotherapy or chemotherapy (both tumor tissue and adjacent normal tissue) in Queen Mary Hospital Hong Kong from 2017-2021, with the volumes of the samples around a few cubic millimeters. The Institutional Review Board of The University of Hong Kong/Hospital Authority Hong Kong West Cluster oversaw the study.

Tissue samples were cut into smaller pieces and subjected to dissociation (50ng/mL EGF, 5mg/mL Collagenase type IV, 5μg/mL DNase I, 0.25% trypsin, 10μM Y-27632, 1× Primocin, 15mM HEPES, in DMEM/F12-Glutamax base medium) with mild agitation for 15-30 mins to dissociate tissue chunks up to 200μm in diameter. Red blood cell (RBC) lysis was performed for samples with significant RBC contents. The cells were then embedded in Matrigel Growth Factor Reduced Basement Membrane Matrix (Corning, Corning, NY) and seeded in ultra-low attachment microplates (Corning), supplied with PDO medium (40% v/v L-WRN CM, additional 2% fetal bovine serum, 1× N-2 supplement, 1× B-27 supplement, 1mM N-Acetylcysteine, 10mM Nicotinamide, 50ng/mL EGF, 10μM SB202190, 0.5μM A83-01, 10μM Y-27632, 10nM Gastrin I, 1× Primocin, 15mM HEPES, in DMEM/F12-Glutamax) for ESCC/EAC tissues and hNEEO medium (10μM Y-27632, 1× Primocin, in Keratinocyte SFM, with additional 0.6mM CaCl_2_) for adjacent normal tissues, respectively. Additionally, hNEEOs were also established from biopsied ESCC tissues by incubating the dissociated tissues with hNEEO medium to selectively promote the growth of non-neoplastic esophageal epithelial cells within the acquired tumor biopsy. ESCC/EAC PDOs and hNEEOs showed distinctive morphological characteristics. The two media were highly selective for the specific organoid type, as PDOs and hNEEOs only grow in the respective medium. Therefore, contamination of different cell types was minimized.

For passaging, Matrigel containing PDO/hNEEO colonies were dissolved and dissociated by incubation with 1× TrypLE Express Enzyme and re-seeded. Passaging was performed every 10-14 days. For cryopreservation, colonies were similarly dissociated, followed by suspension in Recovery Cell Culture Freezing Medium and storage in liquid nitrogen. Multiple freezing/thawing cycles and long-term continuous passaging of established PDOs have been well achieved (continuous passaging over one year until passage 33 with no signs of proliferation defeat or morphological changes); however, proliferative hNEEOs can only be maintained within the first two months of establishment, which is consistent with a previous study (14).

### Plasmids and lentivirus infection

Preparations of plasmids of CRISPR constructs and lentiviral infection on cell lines were performed as previously described (11,12). In brief, oligonucleotides encoding p63- (GCTGAGCCGTGAATTCAACG; TGTGTGTTCTGACGAAACGC), STAT1- (TGCTGGCACCAGAACGAATG), MAVS- (CAGGGAACCGGGACACCCTC), IRF1-targeted sgRNA (CTCCCTGCCAGATATCGAGG), and non-targeting control sgRNAs (GTTCCGCGTTACATAACTTA; CTCTGGCTAACGGTACGCGTA) were cloned into lentiCRISPRv2 vector from Feng Zhang (Addgene plasmid # 52961; http://n2t.net/addgene:52961; RRID: Addgene_52961). The protein depletion efficiency of sgRNA of each target was verified individually by Western Blotting (WB) analysis and pooled together in experiments. For genetic manipulation on organoid cultures by lentiviral infection, organoid cultures were dissociated, and the single-cell suspensions in the respective medium were incubated with viral particles and polybrene overnight on top of a layer of Matrigel; medium containing residual viral particles, polybrene, and unattached cells were removed followed by covering of another layer of Matrigel to fully embed all cells within the matrix before addition of selection antibiotics. The pLenti.PGK.blast-Renilla_Luciferase vector from Reuben Shaw (Addgene plasmid # 74444; http://n2t.net/addgene:74444; RRID: Addgene_74444) was used to label cell lines and organoid cultures for *in vitro* cell viability measurements. For the inducible CRISPR/Cas9 system, the Lenti-iCas9-neo vector from Qin Yan (Addgene plasmid # 85400; http://n2t.net/addgene:85400; RRID: Addgene_85400) was used to express a doxycycline-inducible Cas9 in cell lines, while the sgRNA was delivered separately using the LentiGuide-puro vector from Feng Zhang (Addgene plasmid #52963).

### Animal experiments

Subcutaneous injection of cancer cells was performed as previously described (11). BALB/cAnN-nu (Nude) mice, C.B-17/Icr-scid (SCID) mice, and NOD.CB17-Prkdc^scid^/J (NOD/SCID) mice were used as indicated. Animals were housed in individually ventilated cages under a 12:12 dark/light cycle within environmentally controlled rooms. All experimental procedures were approved by the Committee on the Use of Live Animals in Teaching and Research and performed in AAALAC International accredited Centre for Comparative Medicine Research of the University of Hong Kong Li Ka Shing Faculty of Medicine under licenses from the Hong Kong SAR Government’s Department of Health.

For *in vivo* doxycycline induction experiments in CDXs, engineered cells were subcutaneously inoculated. The doxycycline-induced CRISPR protein depletion started when the tumors reached approximately 100mm^3^ in size. The mice of all groups were supplemented with 0.2mg/mL doxycycline and 2.5% sucrose in drinking water.

### Cell line *in vitro* viability test

Luciferase-labeled cells were quantified by bioluminescence-based live-cell imaging on a CLARIOstar Plus microplate reader (BMG Labtech, Ortenberg, Germany) using Enduren (Promega Corporation, Madison, WI) as the substrate for Renilla luciferase.

### Immunohistochemical staining of *in vitro* PDO cultures

Intact PDO cultures in Matrigel were fixed by 4% paraformaldehyde in PBS for 2 hours at room temperature. Fixed Matrigel containing PDO cultures were gently dissociated by pipetting and embedded in HistoGel Specimen Processing Gel (Thermo Fisher Scientific), followed by standard immunohistochemical procedures as previously described (11).

### Transcriptomic profiling and pathway analysis

RNA sequencing was performed at the University of Hong Kong Li Ka Shing Faculty of Medicine Centre for PanorOmic Sciences and analyzed as previously described (12). We sequenced the ribosomal RNA-depleted total RNA from two ESCC cell lines, KYSE180TS and KYSE450, with ΔNp63 depletion and control in duplicates. We sequenced the poly-A enriched RNA of the ESCC/EAC/hNEEO organoid cultures panel. The transcriptomic data are available (BioProject ID PRJNA995358 for ΔNp63 depletion in cell lines and PRJNA1009949 for organoid profiling). GSEA and GeneMANIA network analyses were performed as described (15,16). Endogenous retrotransposon expression profiling by TEcounts(17) was performed as described (5).

For TF target enrichment analysis on macrophages from the single-cell transcriptomic data, global gene expression profiles of the macrophages in each sample were gathered, normalized, and subjected to correlation analysis with the cancer cell *TP63* expression in the same sample. Genes significantly positively or negatively correlated to cancer cell *TP63* expression were subjected to TF target enrichment analysis using the Human MsigDB TFT transcription factor target geneset database. Only genes of top 2000 expression levels in macrophages, while showing expression in more than half of the samples, were considered due to the limitation on primary immune cells profiling of the 10X Chromium Single Cell RNA sequencing platform (18).

### RNA and protein expression assays

Quantitative PCR and WB analysis were performed as previously described(11). Data of quantitative PCR were normalized to the expressions of *RPS13* or *HSPA4* as indicated. Data of WB analysis were normalized to the expression of p84, Histone H3, or Histone H2A as indicated. Quantitative fluorescent WB data was acquired and analyzed using a Typhoon 5 Biomolecular Imager (Cytiva, Marlborough, MA).

For dsRNA analysis of endogenous retrotransposon expression, single-stranded RNAs (ssRNA) were depleted by digestion with a ssRNA-specific RNase as described (7). The quantitative dsRNA expression was then normalized to the ssRNA expression of *HSPA4*.

### LINE1 methylation assay

LINE1 methylation was used as the indicator to assess the whole-genome methylation of retrotransposons. Genomic bisulfite conversion was performed as previously described (19). QPCR was used to evaluate the LINE1 methylation and unmethylation level, and the relative methylation index was calculated by the formula 2^Ct(unmetLINE1)-Ct(metLINE1)^ as described (20). The hypomethylated EBV-infected 550 cell line and the hypermethylated C666 cell line were used as the negative and positive controls, respectively (19).

### ESCC PDO drug treatment

ESCC PDOs were seeded in ultra-low attachment 96-well plates (embedded in 50uL Matrigel per well) for three days before treatments started. Treatments started in PDO medium without A83-01, SB202190, or Y-27632 (-ASY) to minimize non-specific drug interactions; expanded PDO colonies (seeded in Matrigel for ≥2 days) show comparable growth in both PDO and -ASY media. Treatments lasted for six days with replenishment of drug-containing media. Endpoint colony formation was quantified by bioluminescence imaging.

### Flow cytometry analysis

For *in vitro* cancer cell HLA expression analysis, cells were dissociated, filtered, and incubated with the conjugated antibody for one hour before being subjected to flow cytometry analysis on an ACEA NovoCyte Quanteon analyzer (Agilent, Santa Clara, CA). For xenograft TME cellular profiling and *in vivo* cancer cell HLA expression analysis, xenografts were dissociated into RBC-lysed single-cell aliquots for cryopreservation following the above procedures for organoid culture establishment without the EGF treatment. Aliquots were thawed and incubated with the panel of antibodies following the blockade of non-specific binding of the immunoglobulin to the Fc receptors by TruStain FcX (BioLegend, San Diego, CA). LIVE/DEAD Fixable Near-IR Dead Cell Stain dye (Thermo Fisher Scientific) was used to denote live and dead cells. EPCAM and H-2K^d^ expressions were used to differentiate human cancer cells and mouse stromal/immune cells, respectively.

### Deconvolution

We estimated the proportion of cell types of interest according to the bulk RNA-sequencing data by performing a deconvolution analysis using the CIBERSORT package (21). As the gene expression patterns of cells vary with the microenvironment (e.g., healthy epithelial tissue or areas of different cancer types), we constructed a signature matrix of expected gene expression levels for the major cell types in ESCC samples as described (22). The matrix was derived based on the single-cell sequencing data (23) by separating the cells into 14 cell types and selecting cell type-specific genes. The selected genes are significantly enriched in one cell type with *p*<0.01 in the Wilcoxon test with Benjamini-Hochberg correction.

### Cancer Cell Line Encyclopedia (CCLE) lung SCC cell line analysis

The CCLE cell line transcriptomic data was acquired from the Dependency Map portal (https://depmap.org/portal/). The lung SCC cell lines were categorized according to the *TP63* Transcripts Per Million (TPM) score and denoted as *TP63^-^* cell lines (N=15; TPM ranging from 0.029 to 0.333) and *TP63^+^* cell lines (N=11; TPM ranging from 1.057 to 9.147). The latter, including CALU1, SW900, SKMES1, EPLC272H, KSN62, HARA, LUDLU1, LC1F, LC1SQSF, HCC95, and HCC2814, showed a comparable *TP63* expression profile (medium TPM = 6.403) to CCLE ESCC cell lines (TPM ranging from 0.880 to 8.690; medium TPM = 6.025) and was included for further correlation analysis.

### Statistical analysis

Independent samples *t*-test was applied unless indicated otherwise. A *p*-value less than 0.05 was considered statistically significant. All tests of significance were 2-sided. The error bars shown in the figures represent the 95% confidence interval. For multiple-test comparisons, the p-value was adjusted by the Benjamini-Hochberg correction. An adjusted *p*-value of less than 0.05 is considered significant. An adjusted *p*-value of less than 0.1 is considered marginally significant.

### Data and materials availability

Transcriptomic datasets are available in public repository as described. All cultures are available from the authors upon request.

## RESULTS

### ΔNp63 depletion triggers tumor-suppressive interferon signaling in ESCC cell lines

Most human ESCC tissue samples and *in vitro* cultures predominately express higher levels of *TP63* variants encoding ΔNp63, with a lack of expression of variants encoding TAp63 (Fig. 1A; Supplementary Table 3). ESCC cell lines show differential expression of ΔNp63 protein (Fig. 1B). To elucidate the functional influence of ΔNp63, we deployed an efficient and specific clustered regularly interspaced short palindromic repeats (CRISPR)-mediated protein depletion procedure targeting the *TP63* locus to deplete ΔNp63 protein expression in ESCC cell lines (Supplementary Fig. 1A). Tumorigenesis assays on athymic nude mice by inoculations of ΔNp63-depleted cells confirmed that ΔNp63 expression is essential for ESCC tumorigenesis (Fig. 1C). The effect of ΔNp63 depletion in established xenografts was further demonstrated using an inducible CRISPR-mediated protein depletion system (Fig. 1D; Supplementary Fig. 1B and 1C) since p63 inhibitors are unavailable. Consistently, ΔNp63 depletion in *in vitro* cell culture models decreased cell viability (Fig. 1E).

**Figure 1.**
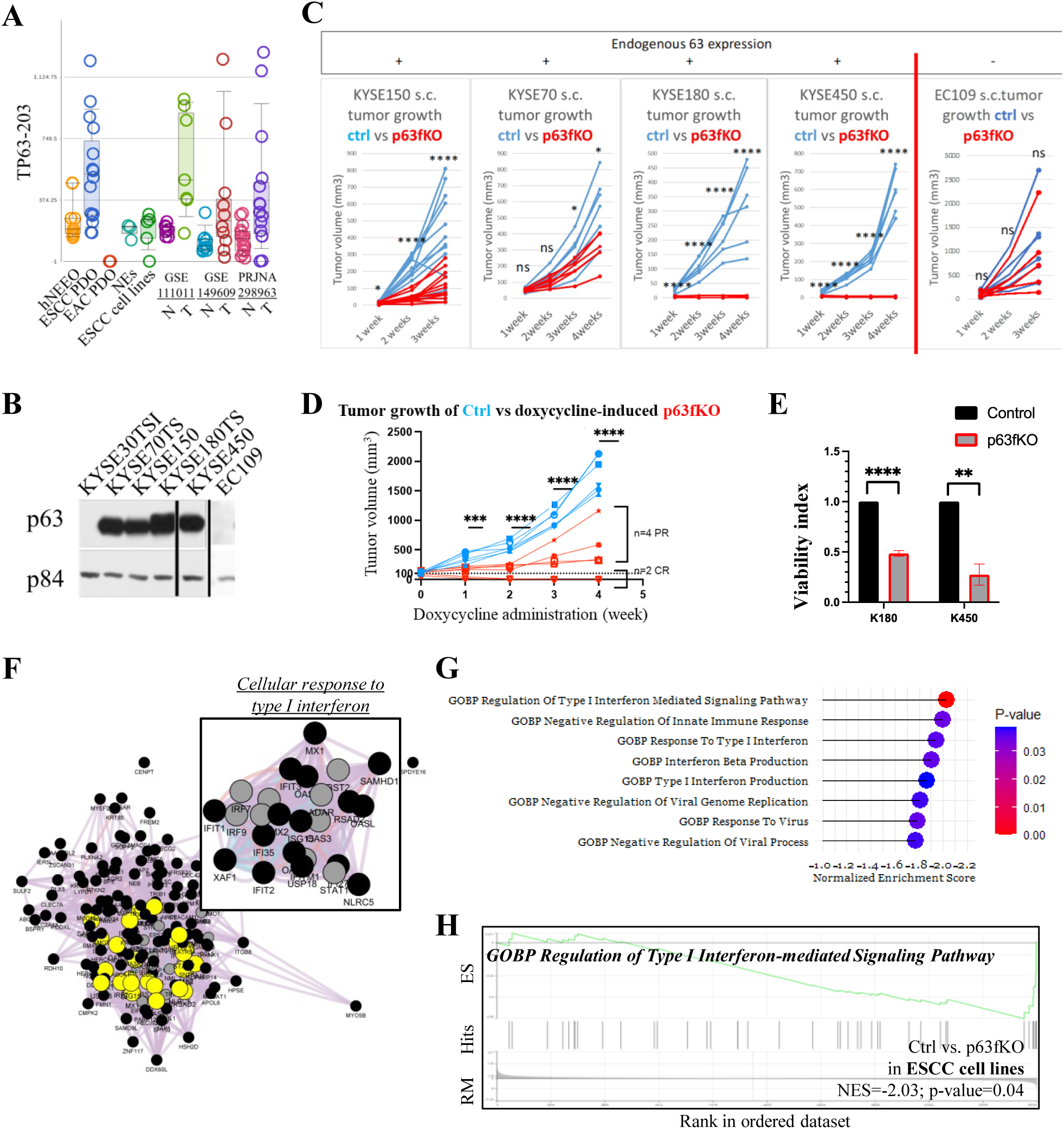
**A**) Summary of the RNA expression of *TP63-203* (encoding the dominant ΔNp63 isoform) in a panel of PDOs and patient normal/ESCC tissue samples. See Supplementary Table 3 for the detailed statistical analysis. NE: normal immortalized esophageal epithelial cell lines; N: patient normal esophageal tissues; T: patient ESCC tissues. **B**) WB analysis shows the expression of ΔNp63 in a panel of ESCC cell lines. **C**) Subcutaneous tumorigenicity assay reveals the critical oncogenic role of ΔNp63. The tumorigenesis of a ΔNp63-negative cell line EC109 was unaffected upon administration of the CRISPR procedure, affirming the specificity of the procedure. Ctrl: non-targeted oligo controls; p63fKO: <u>p63</u> protein functional <u>k</u>nock<u>o</u>ut (fKO). **D**) Induced ΔNp63 depletion on established xenografts leads to significantly suppressed growth and even complete tumor regression. PR: partial response; CR: complete regression. **E**) ΔNp63 depletion results in reduced viability *in vitro* in KYSE180TS (K180) and KYSE450 (K450) cells. **F**) The functional gene network by GeneMANIA analysis of upregulated genes upon ΔNp63 depletion (Supplementary Table 4). Highlighted and top right: the top significant annotation. Black circle: input genes; Grey circle: calculated related genes. **G**) Top IFN-I signaling-related genesets significantly correlate with ΔNp63 depletion in both K180 and K450 by GSEA. A negative normalized enrichment score (NES) indicates enrichment in ΔNp63-depleted cells compared to control cells. See Supplementary Table 6 for the complete list of genesets. **H**) A representative geneset enriched in ΔNp63-depleted cell lines compared to control cells, from G. RM: Rank metric; ES: Enrichment score. ****, p-value < 0.0001; ***, p-value < 0.001; **, p-value < 0.01; *, p-value < 0.05; ns, p-value > 0.05.

The molecular impacts upon ΔNp63 depletion in ESCC cell lines were further examined by transcriptomic profiling to reveal the differentially-expressed genes significantly altered in both cell lines tested (Supplementary Table 4). Interestingly, gene network analysis by GeneMANIA identified IFN-I signaling among the top networks enriched in ΔNp63-depleted cells (Fig. 1F; Supplementary Table 5), which was further verified by gene set enrichment analysis (GSEA) showing IFN-I signaling-related pathways, as the top gene ontology pathway enriched upon ΔNp63 depletion (Fig. 1G and 1H; Supplementary Table 6).

### *TP63* expression level shows negative correlations with IFN-I signaling enrichment in ESCC patient-derived organoids (PDOs)

Cancer patient-derived organoid (PDO) cultures preserve the malignant epithelial compartment and serve as better models than traditional cell lines, with more significant heterogeneity and better representation of cancer phenotypic and molecular spectra(24). Short-term culture of ESCC biopsy samples has been explored, which demonstrates promising clinical application potential (25). To facilitate ESCC research, we established a biobankable panel of human non-neoplastic esophageal epithelial organoids (hNEEO) and ESCC PDO cultures from freshly collected patient tissue samples (Fig. 2A, 2B, and 2C; Supplementary Table 7). Consistent with previous studies (26), ESCC PDOs express a comparably higher level of *TP63* compared to hNEEO, while esophageal adenocarcinoma PDOs do not express *TP63* (Fig. 1A).

**Figure 2.**
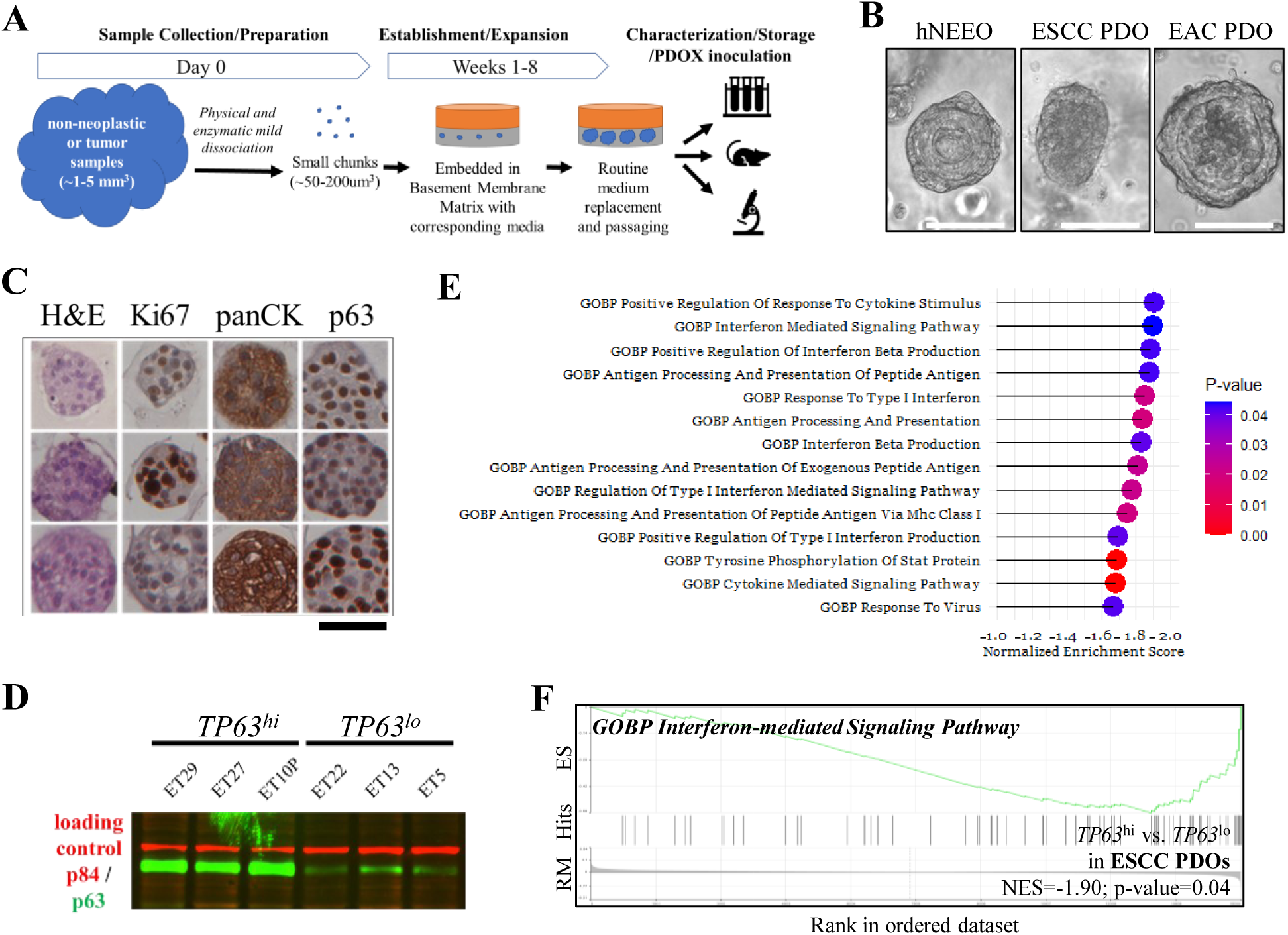
**A**) An illustration of the organoid establishment workflow. Samples were acquired through surgery or endoscopic examination. **B**) Representative microscopic images of *in vitro* hNEEO, ESCC PDO, and EAC PDO cultures showing distinctive morphological features of the colonies. Scale bar:100um. **C**) Representative histological/IHC analyses showing proliferative squamous cell carcinoma-specific features of the ESCC PDO cultures. Scale bar 50 μm. H&E: Hematoxylin and Eosin staining; panCK: pan-cytokeratin staining. **D**) WB analysis verifies the varied relative expression of ΔNp63 in ESCC PDOs with the lowest (*TP63^lo^*) and highest expressions (*TP63^hi^*), ranked based on the *TP63* RNA expressions across the panel of PDOs. **E**) Top IFN-I signaling-related genesets negatively correlated with ΔNp63 expression in PDOs by GSEA. A negative NES indicates enrichment in *TP63^lo^* PDOs compared to *TP63^hi^* PDOs. See Supplementary Table 8 for the complete list. **F**) A representative geneset enriched in *TP63^lo^* PDOs compared to *TP63^hi^* PDOs, from E.

To complement the findings from the genetically manipulated cell lines, GSEA of the transcriptomic data of the unmanipulated ESCC PDO panel consistently showed that *TP63* RNA expression level was negatively associated with IFN-I signaling-related pathway enrichment (Fig. 2D, 2E, and 2F; Supplementary Table 8).

### *Signal transducer and activator of transcription 1 (STAT1)* mediates the IFN-I signaling regulation by ΔNp63

Cancer cell IFN-I signaling plays tumor-suppressive roles (27–29). We verified the upregulation of a panel of IFN-I-regulated interferon-stimulated genes (ISGs) (4,5) upon ΔNp63 depletion in ESCC cell lines and xenografts by quantitative PCR (QPCR) (Fig. 3A and 3B). *STAT1* is among the primordial signal mediators for IFN-I signaling (30) and plays a tumor-suppressive role in ESCC (31). Consistently, GSEA focusing on TF targets revealed that genesets of STAT1 target genes were enriched in cell lines upon ΔNp63 depletion and in ESCC PDOs with the lowest *TP63* expression (Fig. 3C, 3D, and 3E; Supplementary Tables 9 and 10), as compared to control cells and ESCC PDOs with the highest *TP63* expression, respectively. Furthermore, we found that ΔNp63 depletion upregulates the activated phosphorylated-STAT1 in ESCC cell lines (Fig. 3F). STAT1 depletion dramatically rescues the reduced viability upon ΔNp63 depletion (Fig. 3G; Supplementary Fig. 2A), suggesting the critical involvement of STAT1/IFN-I signaling in mediating reduced cell viability upon ΔNp63 depletion. Together, these data support an inhibitory role of ΔNp63 in IFN-I signaling in ESCC cells. IFN-I signaling has previously not been connected to p63/ΔNp63 in any cancer type.

**Figure 3.**
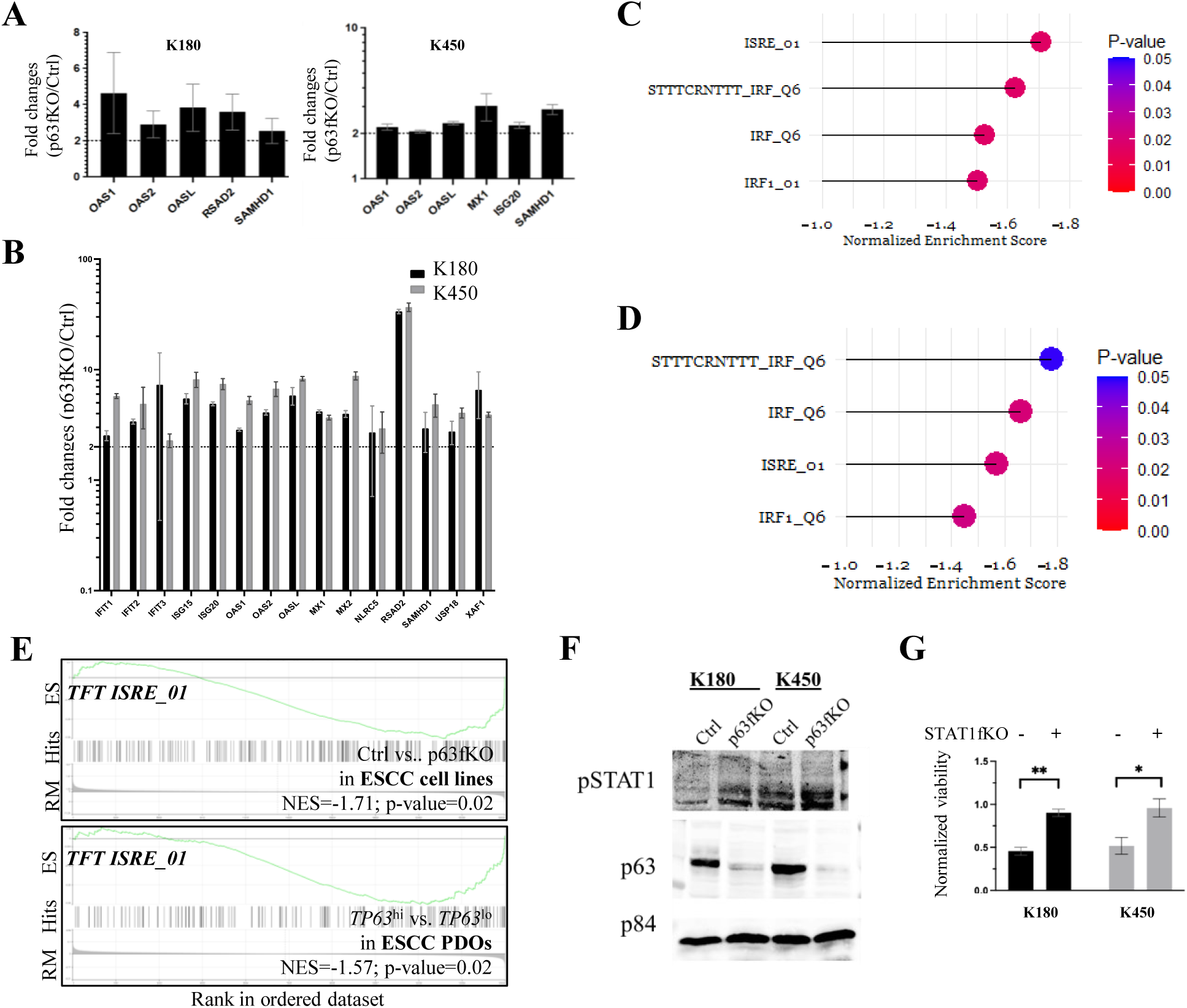
**A** and **B**) ΔNp63 depletion upregulates ISGs in *in vitro* cell line cultures (A) and xenografts (B) by QPCR analysis. Dotted line: 2-fold changes. **C** and **D**) Top TF-binding signature genesets enriched in ΔNp63-depleted cells in both K180 and K450 cell lines (C) and in *TP63^lo^* PDOs (D), as compared to control cells and *TP63^hi^* PDOs, respectively. **E**) GSEA results of the gene set ISRE_01 (STAT1 targets) in ΔNp63-depleted cell lines (Upper) and ESCC PDO cultures with varied p63 expression (Bottom), from C and D. **F**) ΔNp63 depletion increases STAT1 tyrosine 702 phosphorylation (pSTAT1) level by WB analysis in ESCC cell lines. **G**) CRISPR-mediated STAT1 depletion dramatically rescues the reduced cell viability upon ΔNp63 depletion. Shown are the normalized viability of ΔNp63-depleted cells to the corresponding control cells.

### IFN-I signaling regulation by ΔNp63 is cancer-specific

To examine whether ΔNp63 regulates IFN-I signaling in pre-malignant cells, we performed transcriptomic profiling on hNEEO cultures followed by GSEA. In contrast to the ESCC cell line and PDO findings, the expression of *TP63* is not significantly associated with IFN-I signaling-related pathways in hNEEO cultures (Supplementary Table 11). ΔNp63 depletion in NE1, a commonly used immortalized esophageal epithelial cell line, showed no alterations of ISG expression (Supplementary Fig. 2B). ΔNp63 may play distinct roles in ESCC compared to pre-malignant esophageal epithelial cells, likely due to the differential regulation of retrotransposon expression among normal and malignant cells.

### ΔNp63 depletion derepresses endogenous retrotransposon expression

IFN-Is are canonical inducers of the IFN-I signaling through an autocrine/paracrine manner. We reasoned that interferon expression mediates the IFN-I signaling induction upon ΔNp63 depletion. However, genes encoding several IFN-Is had negligible basal expression in ESCC PDOs and cell lines and were not upregulated upon ΔNp63 depletion (Supplementary Table 12). This is most likely due to the frequent loss of the chromosome 9p21 region containing genes encoding several IFN-Is in ESCC (32,33), which further indicates a potential tumor-suppressive role of cancer cell-derived IFN-Is (34,35).

Alternatively, the formation of cytosolic dsRNA derived from the expression of endogenous retrotransposon-encoded RNAs is recognized by cytoplasmic and endosomal sensor proteins and induces IFN-I signaling. A retrotransposon expression quantification algorithm, TEcounts, was applied to quantify global retrotransposon expression (5,17) using the ΔNp63-depleted cell line transcriptomic data and to identify a list of differentially-expressed retrotransposons (Supplementary Table 13). Interestingly, ΔNp63 depletion increased retrotransposon expression (Fig. 4A), which was verified through dsRNA-enriched QPCR examination (Fig. 4B).

**Figure 4.**
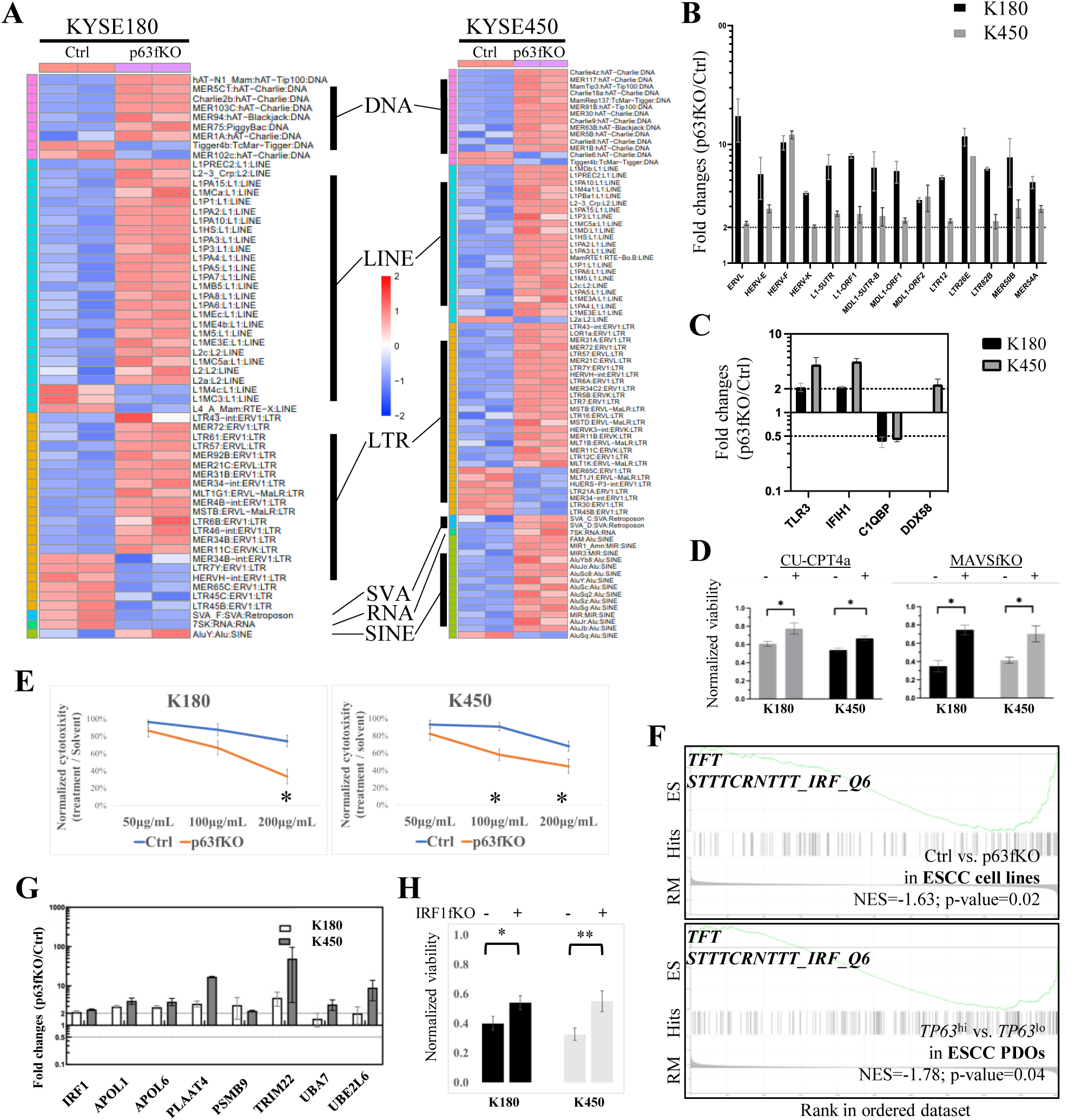
**A**) Endogenous retrotransposon expression profiling in ESCC cell lines upon ΔNp63 depletion shows generally increased expression of retrotransposons. A total of 773 and 760 expressed retrotransposons were identified in the two cell lines, respectively. Shown are all retrotransposons with significantly altered expression (adjusted p-value < 0.05) upon ΔNp63 depletion (60 retrotransposons in K180 and 83 in K450). DNA: DNA transposon; LINE: Long interspersed nuclear element; LTR: Long terminal repeat; SVA: SINE-VNTR-Alus; RNA: small RNA retrotransposon; SINE: Short interspersed nuclear element. **B**) Increased retrotransposon expression upon Np63 depletion was verified by QPCR following dsRNA enrichment. RNA expression was normalized to the single-stranded RNA expression of *HSPA4*. **C**) RNA expression of dsRNA sensors is upregulated upon p63 depletion in ESCC cell lines. The expression of *Complement C1q binding protein* (*C1QBP*), a negative regulator of the sensors, was employed as a control. Data normalized to *HSPA4* expression. **D**) Inhibition of the dsRNA sensing of TLR3 by TLR3/dsRNA interaction inhibitor CU-CPT5a and depletion of MAVS by CRISPR-mediated protein depletion attenuates the proliferation suppressive effect upon ΔNp63 depletion. Shown are the normalized viability of ΔNp63-depleted cells to the corresponding control cells. **E**) ΔNp63-depleted cells demonstrate higher sensitivity to poly(I:C) treatment, leading to further reduced cell viability in ESCC cell lines. **F**) GSEA result of the gene set STTTCRNTTT_IRF_Q6 (IRF1 targets) related to IRF1 binding in ΔNp63-depleted cell lines (Upper) and PDO cultures with varied p63 expression (Bottom), from Fig. 3C and 3D. **G**) ΔNp63 depletion upregulates IRF1 target genes in cell lines. Data normalized to *HSPA4* expression. The dotted line indicates two-fold changes. **H**) CRISPR-mediated IRF1 depletion partially rescues the reduced cell viability upon ΔNp63 depletion. Shown are the normalized viability of ΔNp63-depleted cells to the corresponding control cells.

Expression of RNA-induced silencing complex components regulating retrotransposon expression in cancer cells (5,36) was not altered upon ΔNp63 depletion in ESCC cell lines (Supplementary Table 14). The global genomic DNA methylation and specific histone modification marks, the two key mechanisms maintaining the repression of retrotransposons (37), showed no alterations upon ΔNp63 depletion (Supplementary Fig. 3A and 3B).

### ΔNp63 regulates canonical dsRNA sensors and suppresses dsRNA sensing

We further profiled the cellular dsRNA sensory machinery. The pattern recognition receptors, including the *Toll-like receptor 3* (*TLR3*), *DexD/H-Box Helicase 58* (*DDX58*) encoding RIG-I, and *Interferon induced with helicase C domain 1* (*IFIH1*) encoding MDA-5, play essential roles in cellular dsRNA sensing and IFN-I signaling induction (38). Interestingly, QPCR analysis demonstrated the upregulated expression of all three sensors upon ΔNp63 depletion (Fig. 4C). Chemical inhibition of the TLR3-dsRNA interaction or genetic depletion of the mitochondrial antiviral protein (MAVS), the common adaptor protein for RIG-I and MDA-5, partially restores the reduced cell viability upon ΔNp63 depletion (Fig. 4D; Supplementary Fig. 3C). Furthermore, exogenous synthetic dsRNA analog polyinosinic:polycytidylic acid (polyI:C), known to trigger cellular dsRNA sensing and antiviral/anticancer signaling induction (39), further reduced cell viability upon ΔNp63 depletion (Fig. 4E), which verifies the enhanced dsRNA sensory machinery in ΔNp63-depleted cells.

### ΔNp63 suppresses Interferon regulatory factor 1 (IRF1) signaling

Key pathways and mediators downstream of dsRNA sensing in cancer cells were further scrutinized. The *TANK-binding kinase 1* (*TBK1*)/*interferon regulatory factor 3* (*IRF3*) axis mediates the cellular dsRNA sensory machinery activation and antiviral/anticancer signaling induction (8). However, no rescue effect on cell viability was observed when ΔNp63-depleted cells were treated with a TBK1 inhibitor (Supplementary Fig. 3D). Alternatively, *IRF1* has been shown to mediate IFN-I signaling independent of other interferon regulatory factors, including *IRF3* (40), in a STAT1-dependent (41) or independent manner (42). Interestingly, genesets containing IRF1 target genes showed significant enrichment in ΔNp63-depleted cells and ESCC PDOs with the lowest *TP63* expression (Fig. 3C, 3D, and 4F; Supplementary Tables 9 and 10), as compared to control cells and ESCC PDOs with the highest *TP63* expression, respectively. Upregulation of expression of *IRF1* and known IRF1 targets (*APOL1*, *APOL6*, *PLAAT4*, *PSMB9*, *TRIM22*, *UBA7*, and *UBE2L6*) (42) was observed in differentially-expressed genes upon ΔNp63 depletion (Supplementary Table 4), which was verified by QPCR (Fig. 4G). IRF1 depletion partially rescues the reduced cell viability upon ΔNp63 depletion (Fig. 4H; Supplementary Fig. 3E). These data indicate that IRF1 meditates the anticancer effects downstream of ΔNp63 depletion.

### cGAS-STING is involved in IFN-I signaling induction upon ΔNp63 depletion

In addition to the canonical antiviral pathways involved in dsRNA sensing, the classic cytosolic DNA-sensing cGAS–STING axis also exerts intracellular antiviral responses in cancer cells (43). STING also plays roles in RNA virus sensing and potentiates the induction of IFN-I signaling and other antiviral signaling pathways (43). A STING agonist (44) suppressed the viability of a panel of ESCC PDO cultures (Fig. 5A), verifying the tumor-suppressive role of the activated STING in ESCC.

**Figure 5.**
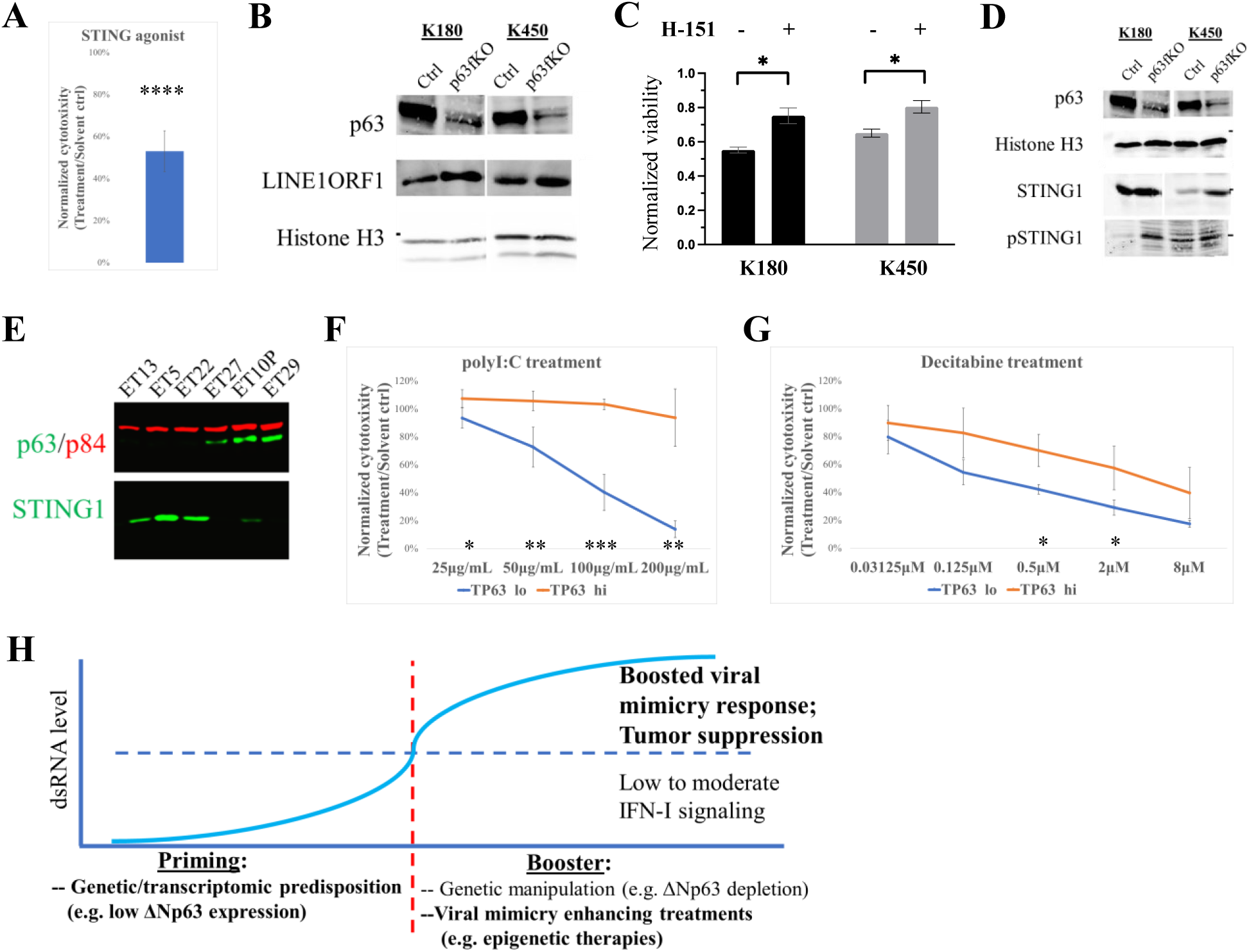
**A**) A cGAS-STING agonist G10 exerts a tumor-suppressive role in a panel of ESCC PDO cultures (N=5). **B**) The protein expression of LINE1 ORF1 is upregulated upon ΔNp63 depletion in ESCC cell lines. **C**) Inhibition of STING1 function by small molecular inhibitor H-151 attenuates the proliferation suppressive effect upon ΔNp63 depletion. Shown are the normalized viability of ΔNp63-depleted cells to the corresponding control cells. **D**) Protein expressions of total STING1 and phosphorylated STING1 (pSTING1) are upregulated upon ΔNp63 depletion in ESCC cell lines by WB analysis. **E**) STING1 and ΔNp63 protein expression show negative correlations in ESCC PDOs by WB analysis. Nuclear protein p84 was used as the loading control. **F** and **G**) ESCC *TP63^lo^* PDOs significantly respond to poly(I:C) treatments (F) and Decitabine treatments (G) compared to *TP63^hi^* PDOs. **H**) Proposed principle for viral mimicry-inducing treatments. Inspired and modified from (8).

Elevated retrotransposon expression, especially LINE-1 retrotransposon, leads to the accumulation of cDNA intermediates and potentially triggers the cGAS-STING (45,46). LINE-1 ORF1 expression increased upon ΔNp63 depletion (Fig. 5B). Administration of a small chemical STING inhibitor leads to partial rescue of the reduced cell viability in ΔNp63-depleted cells (Fig. 5C). ΔNp63 depletion increased STING protein expression in cell lines (Fig. 5D). A similar negative trend of a ΔNp63/STING-expression was observed in the ESCC PDO panel (Fig. 5E). These data suggest that elevated STING expression and enhanced cGAS-STING axis play significant roles in mediating the anticancer effects downstream of ΔNp63 depletion.

All findings suggest a novel function of ΔNp63 in repressing retrotransposon expression and suppressing dsRNA sensing in cancer cells; ΔNp63 depletion triggers viral mimicry response and suppression.

### ΔNp63 expression level indicates therapeutic opportunities for employing cancer cell-targeted viral mimicry boosters

We further explored potential ΔNp63 expression-guided therapeutic opportunities. ΔNp63 depletion in cancer cells results in enhanced dsRNA sensing and sensitizes cells to synthetic dsRNA analog polyI:C treatment in ESCC cell lines (Fig. 4E). PolyI:C and its stabilized derivative polyICLC have been explored clinically to inhibit tumor growth via regulating proliferation/apoptosis (39). Interestingly, ESCC PDOs with the lowest ΔNp63 expression levels demonstrate hypersensitivity to polyI:C treatment, compared to PDOs with the highest ΔNp63 expression (Fig. 5F). This verifies the findings from the cell lines and further suggests that such increased dsRNA sensing in ESCC cells with lower *TP63* expression can be explored pharmaceutically.

Cancer cells maintain retrotransposon expression at a sublethal level to minimize tumor-suppressive antiviral/IFN-I signaling induction (8). Interestingly, treatment of Decitabine, a clinically approved anticancer agent, performed on the ESCC PDO panel showed that PDOs with the lowest ΔNp63 expression respond better to the treatment (Fig. 5G), indicating that ΔNp63 expression may serve as a biomarker for Decitabine responsiveness or other viral mimicry boosting treatments. Based on our findings, we hypothesized that cancer cells with lower ΔNp63 expression may possess higher basal IFN-I signaling activity that is near or beyond tolerable thresholds; the cells may, therefore, become exquisitely vulnerable to further treatments such as Decitabine treatment that magnifies retrotransposon expression and viral mimicry responses to suppress tumor growth (6,8) (Fig. 5H).

### *TP63* expression negatively correlates with TIIC signatures in ESCC patient samples

We have examined the functional influence of ΔNp63 expression in cancer cells. Additionally, cancer cell viral mimicry response and IFN-I signaling boost antitumor immunity in several aspects, including elevating antigen presentation, reducing immunosuppression, and enhancing the recruitment and activation of cytotoxic TIICs (4–8,34,36). To explore the influence of ΔNp63 on TIICs in ESCC, we analyzed public bulk transcriptomic datasets of clinical samples. First, a non-deconvolution The Cancer Genome Atlas (TCGA) data-derived metagene/GSEA-based immune cell abundance signature profile (47) was used for *TP63* correlation analysis (48). Remarkably, general negative correlations between *TP63* expression and TIIC signatures were observed in ESCC (Fig. 6A). Specifically, molecular signatures of tumor-infiltrating monocytes, tumor-associated macrophages (TAMs), and effector memory CD8^+^ T cells were among the most negatively correlated immune cell signatures with *TP63* expression.

**Figure 6.**
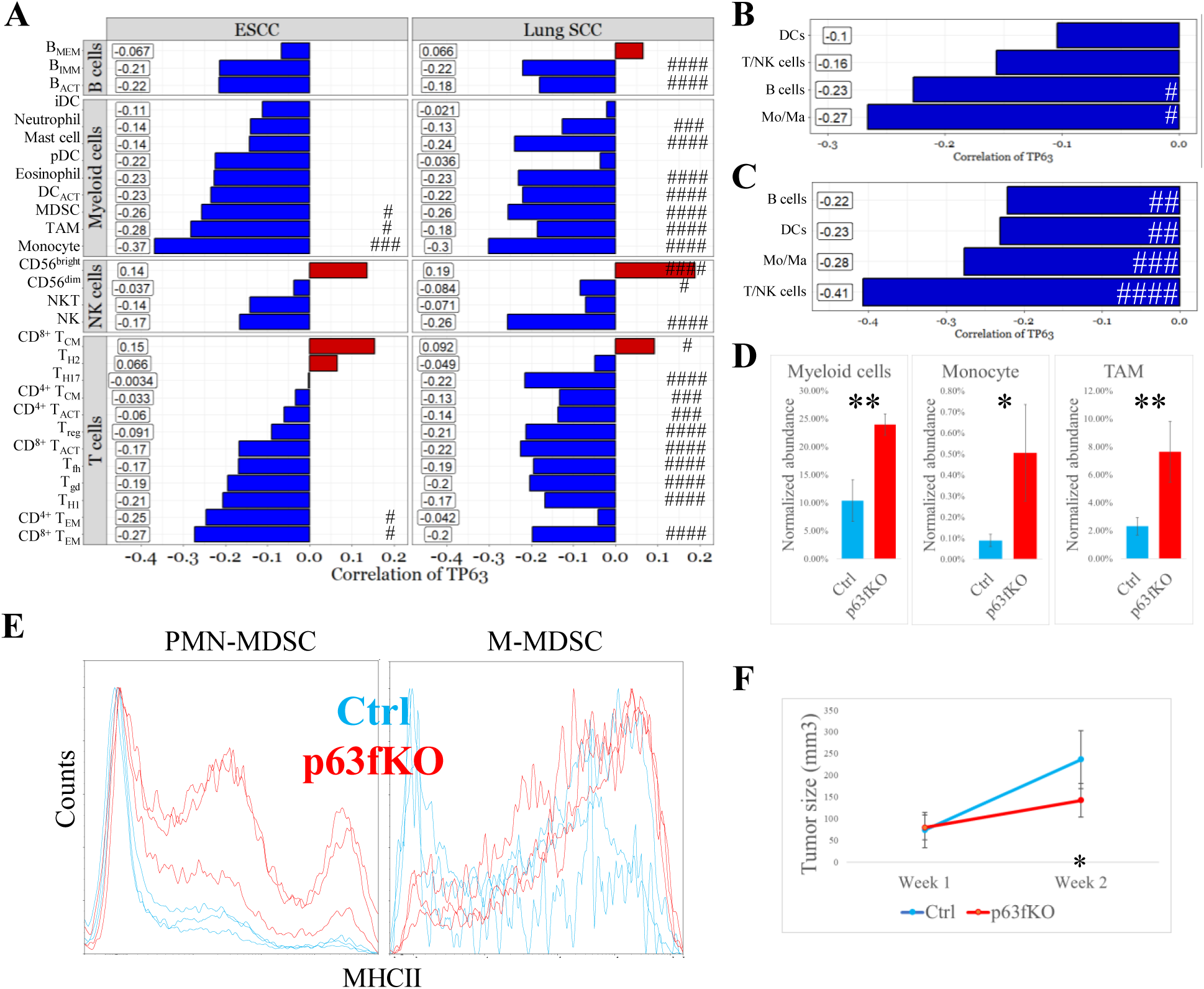
**A**) Negative correlations between *TP63* expression and multiple TIIC signatures are observed in the non-deconvolution TCGA immune cell profiles. B_MEM_: memory B cells; B_IMM_: immature B cells; B_ACT_: activated B cells; iDC: interstitial dendritic cells; pDC: plasmacytoid DC; DC_ACT_: activated DC; T_CM_: central memory T cells; T_H2_: T helper 2 cells; T_H17_: T helper 17 cells; T_ACT_: activated T cells; T_reg_: regulatory T cells; T_fh_: T follicular helper cells; T_gd_: γδ T cells; T_H1_: T helper 1 cells; T_EM_: effector memory T cells. **B** and **C**) Deconvolution analyses on bulk ESCC RNA sequencing data (TCGA ESCC dataset; B) and ESCC microarray data (GSE53624; C) reveal negative correlations between cancer cell *TP63* expression and major TIIC relative abundance. **D**) Elevated proportions of tumor-infiltrating mouse myeloid cells, monocytes, and TAM, as normalized to human cancer cells, are observed in ΔNp63-depleted KSYE450 CDXs on nude mice. N=3 in each group. **E**) Tumor-infiltrating M-MDSCs and PMN-MDSCs with increased MHCII expression were observed in ΔNp63-depleted KSYE450 CDXs on nude mice. **F**) ΔNp63-depleted KYSE450 cells inoculated in NOD-SCID mice display significant but greatly attenuated tumor suppression compared to the tumorigenicity profile observed in nude mice (Fig. 1C). ####, adjusted p-value < 0.001; ###, adjusted p-value < 0.01; ##, adjusted p-value < 0.05; #, adjusted p-value < 0.1.

The latest deconvolution-based cellular profiling method was also employed to decompose immune cell identities and estimate relative cell abundance for public TCGA ESCC bulk RNA-sequencing and microarray datasets. Consistently, cancer cell *TP63* expression level showed a significant negative correlation to T/NK cells, B cells, and tumor-infiltrating monocytes/TAMs in both datasets (Fig. 6B and 6C).

These data indicate that cancer cell *TP63* expression negatively correlates with the abundance of TIICs in patient samples, which suggests that cancer cells with discrete ΔNp63 expression vary in ability to attract immune-cell infiltrates, likely due to IFN-I signaling-mediated cytokine secretions (5). Low ΔNp63 expression in cancer cells may lead to the formation of TIIC-rich tumor mass.

### ΔNp63 depletion results in increased infiltration of reprogrammed myeloid cells

We observed negative correlations between tumor-infiltrating monocytes/TAM signatures and cancer cell *TP63* expression in clinical samples. Consistently, in cell line-derived xenografts (CDXs) examined by flow cytometry following tumor dissociation, a significantly increased proportion of the Cd45^+^Cd49b^-^ myeloid compartment was observed upon doxycycline-induced cancer cell ΔNp63 depletion (Fig. 6D). Specifically, increased Ly6g^-^Ly6c^hi^F4/80^-^ tumor-infiltrating monocyte and Ly6g^-^Ly6c^lo^F4/80^hi^ TAM populations were observed.

Besides tumor-infiltrating monocytes/TAMs, myeloid-derived suppressor cells (MDSCs) also play crucial roles in tumor development and tumor microenvironment (TME) modulation (49–51). Consistent with the Ly6g^-^ tumor-infiltrating monocytes/TAM population, the monocyte-derived Ly6g^-^Ly6c^hi^F4/80^+^ tumor-infiltrating monocytic MDSCs (M-MDSCs) (52) showed a trend of increased infiltration in xenografts of ΔNp63-depleted cells; in contrast, the proportion of Ly6g^+^ cells, commonly recognized as granulocyte-derived tumor-infiltrating polymorphonuclear MDSCs (PMN-MDSCs), remained largely unaltered (Supplementary Fig. 4A). Interestingly, both MDSC populations demonstrated increased proportions of MHCII^hi^ cells in ΔNp63-depleted xenografts (Fig. 6E). This is consistent with a previous study showing that dsRNA analog polyI:C treatment increased MHCII expression in tumor-infiltrating MDSCs to attenuate the immunosuppressive activity of MDSCs (53). The study also showed that polyI:C treatment polarized various myeloid cells into a tumor-suppressive state. These data suggested that ΔNp63 expression modulates intratumoral myeloid recruitment and reprogramming towards a pro-cancer state through suppressing IFN-I signaling.

Much more significant *in vivo* effects upon cancer cell ΔNp63 depletion were observed when inoculated on nude mice (Fig. 1C), as compared to those in the *in vitro* cultures (Fig. 1E). Given the increased myeloid cell infiltration in nude mice xenografts upon ΔNp63 depletion, we hypothesized that the reprogrammed myeloid population plays a significant anticancer role. Immunocompromised NOD/SCID mice differ from athymic nude mice, showing impaired tissue myeloid maturation and mononuclear phagocyte functions (54,55). The ΔNp63-depleted cells were inoculated on NOD/SCID mice. Increased monocyte infiltration in ΔNp63-depleted NOD/SCID xenografts was consistently observed (Supplementary Fig. 4B). However, the TAM population was unaltered, likely reflecting the myeloid maturation defect of NOD/SCID mice. Furthermore, a dampened effect upon ΔNp63 depletion was observed compared to that on athymic mice (Fig. 6F), supporting the hypothesis that reprogrammed myeloid cells, including TAMs, play a critical role in mediating ΔNp63 depletion-induced tumor suppression.

### ΔNp63 modulates MHC class I expression in cancer cells

Antigen processing and presentation by cancer cell MHCI molecules are crucial for antitumor immune surveillance. This is regulated by IFN-I signaling (5,8) and potentiates cytotoxic immune cell recruitment and activation (56,57). Consistently, genes encoding MHCI molecules HLA-A/B/C/F were upregulated in *in vitro* ΔNp63-depleted ESCC cells, along with genes encoding peptide transporters TAP1 and TAP2 (Supplementary Table 15). Flow cytometric analysis confirmed that ΔNp63 regulates HLA-A/B/C cell surface expression in a STAT1-dependent manner, as STAT1 depletion dramatically rescues the reduced HLA expression upon ΔNp63 depletion (Fig. 7A). Furthermore, flow cytometric analysis on the CDX models further showed increased HLA-A/B/C cell surface expression in ΔNp63-depleted ESCC cells from dissociated CDXs on both mouse models tested (Fig. 7B).

**Figure 7.**
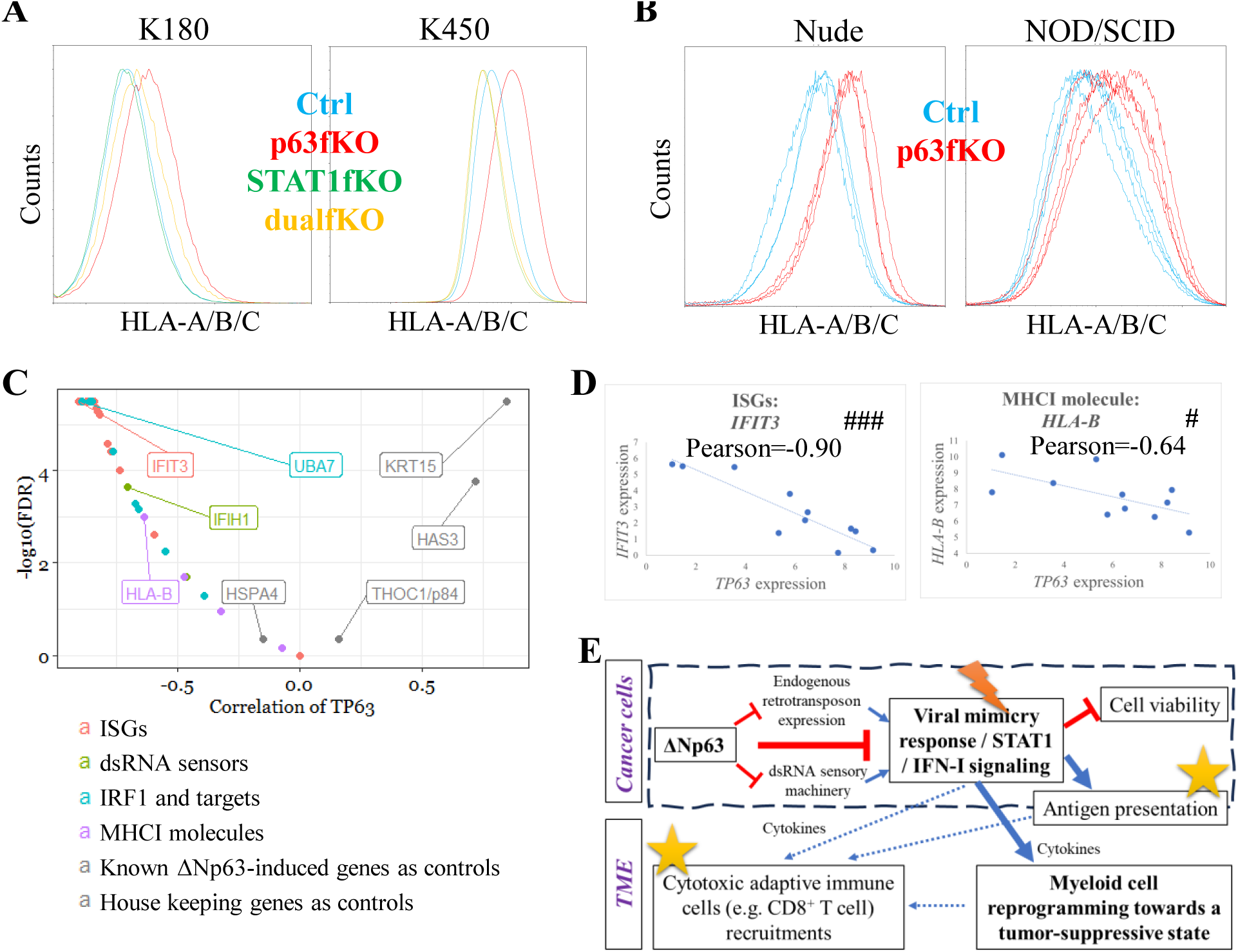
**A**) Cancer cell surface HLA-A/B/C expressions are upregulated in ΔNp63-depleted K180 and K450 cells *in vitro*, which is dramatically rescued upon ΔNp63/STAT1 dual-depletion (dualfKO). **B**) Cancer cell surface HLA-A/B/C expressions are upregulated in ΔNp63-depleted KSYE450 CDXs on both nude and NOD/SCID (N=4) mice models. **C**) The ISGs and related genes regulated by ΔNp63 in ESCC cell lines and xenografts show significant negative correlations with *TP63* expression in CCLE *TP63^+^* lung SCC cell lines (N=11). As controls, known ΔNp63 target genes *KRT15* and *HAS3* show significant positive correlations to *TP63* expression, while housekeeping genes *HSPA4* and *THOC1* show no significant correlations. Genes from different categories detailed in Supplementary Table 16, analyzed by Pearson correlation. **D**) Selected representative negative correlations between *TP63*/*IFIT3* and *TP63*/*HLA-B*, from C. **E**) Summary diagram of the novel oncogenic roles of ΔNp63. Lighting sign indicates a pharmaceutical opportunity for viral mimicry boosting treatment; Stars suggest a pharmaceutical opportunity for ICB therapy.

### *TP63* expression negatively correlates with ISG expression in lung SCC cell lines and with TIIC signatures in lung SCC patient samples

To extend our findings in ESCC to other SCCs, we examined *TP63* in lung SCC, the cancer type sharing similar tissue differentiation and molecular characteristics with ESCC (58) and explicitly expressing ΔNp63 (59). By exploring the public bulk transcriptomic data of a panel of lung SCC cell lines (60), we observed that the majority of ISGs and IFN-I signaling-related genes regulated by ΔNp63 in ESCC cell lines and xenografts also showed specific and significant negative correlations with *TP63* expression in *TP63^+^* lung SCC cell lines (Fig. 7C and 7D; Supplementary Table 16).

Furthermore, the correlation between *TP63* and TIIC infiltration in lung SCC was analyzed. We explored the TCGA data-derived non-deconvolution metagene immune cell abundance signature profile (47). Remarkably, significant negative correlations between *TP63* expression and TIIC signatures were observed in lung SCC clinical samples similar to ESCC (Fig. 6A). Among tumor-infiltrating lymphocytes (TILs), molecular signatures of several CD8^+^ T cell populations, including activated CD8^+^ and effector memory CD8^+^ T cells, showed substantial negative correlations with *TP63* expression. Consistently, *TP63* expression negatively correlated with the tumor-infiltrating monocytes/TAM signatures. These data strongly suggested that ΔNp63 expression in lung SCC cells plays a similar role in suppressing IFN-I signaling and TIIC infiltration.

## DISCUSSION

In this study, we demonstrated that p63/ΔNp63 exerts a previously undefined oncogenic role in SCC (Fig. 7E). ΔNp63 regulates several key aspects of cancer progression, including cancer stem cell maintenance and drug resistance (61). Our study revealed novel cancer-specific functions of ΔNp63 in suppressing cancer cell viral mimicry response and remodeling the TME towards an oncogenic state. In brief, ΔNp63 represses endogenous retrotransposon expression and dsRNA sensing, which restricts cancer cell viral mimicry response and affects both the cancer cell and the TME, in alignment with previous studies on cancer cell viral mimicry response and antitumor immunity in breast cancer, colorectal cancer, melanoma, and ovarian cancer (4–7). In cancer cells, ΔNp63 maintains cell viability, likely through inhibiting STAT1-mediated cell death (62). In the TME, cancer cell ΔNp63 suppresses antitumor TME generation, which may depend on STAT1-IFN-I signaling-mediated immune-regulatory cytokine secretion (7,63).

Human ESCC samples frequently display homozygous loss of the 9p21.3 region, harboring classic tumor suppressors *CDKN2A*/*B* (32,33). Two recent studies revealed an additional selective advantage of losing 9p21.3 in cancer cells, attributed to the IFN-I gene cluster in the region (34,35). Specific disruption of the cluster resulted in substantial changes in infiltrating immune cells and escape from CD8^+^ T cell surveillance. Since the genetic loss of the IFN-I gene cluster is irreversible, interferon production-independent IFN-I signaling activation triggered by viral mimicry response supplies additional anticancer mechanisms.

Cancer cells exhibit elevated retrotransposon expression and activity than normal cells, contributing to cancer initiation and development (64). This is usually accompanied by suppressed sublethal tumor-suppressive IFN-I signaling activation. We showed that enhanced IFN-I signaling triggered by dsRNA stress (expression and sensing) due to experimental manipulation (e.g., ΔNp63 depletion) reduces cancer cell viability. The cells may further develop exquisite sensitivity to additional anticancer treatments that exaggerate the increase of dsRNA expression, leading to maximized viral mimicry response activity and cell death. Drug response profiling on the ESCC PDO panel suggests that cancer cells with a specific genetic or transcriptomic predisposition (e.g., low ΔNp63 expression) exhibit hypersensitivity to viral mimicry-boosting treatments (e.g., polyI:C and Decitabine), in alignment with previous studies (8). Therefore, we hypothesize that ESCC cases with low basal ΔNp63 protein expression, accounting for around 20-40% of all clinical cases (65,66), may be considered “primed” for cancer cell-targeted viral mimicry response boosting and the resultant tumor suppression (Fig. 5H). A similar scenario may apply to lung SCC cases as well. Future studies are needed to explore the patient stratification potential of ΔNp63 expression and the utilization of boosters.

Cancer cell viral mimicry response and IFN-I signaling activation promote the generation of a tumor-suppressive TME and modulate antitumor immunity (4–7). Our analyses on TIIC signatures of bulk transcriptomic data consistently showed that cancer cell *TP63* expression level negatively correlates with the multiple signatures of TILs and tumor-infiltrating myeloid cells in patient samples of ESCC and lung SCC. A recent pioneering study subtyping ESCC utilizing integrated multi-omics profiling described a substantial CD8^+^ T cells- and CD68^+^ myeloid cells-enriched “immune modulation” subtype that specifically possesses low *TP63* expression (67), consistent with our analysis. The study also indicated that patients of the “immune modulation” subtype respond better to immune checkpoint blockade (ICB) treatments than other subtypes. Another milestone multi-omics study subtyping lung SCC identified a “classical” subtype featuring amplification and expression of *TP63*/ΔNp63 along with downregulation of immune signaling (59). Immunotherapy has been increasingly explored and utilized in ESCC and lung SCC management (68,69). Identifying patients responsive to immunotherapy is crucial. Our analysis provides a mechanistic insight into these clinical subtyping analyses. It suggests that *TP63*/ΔNp63 expression in ESCC and lung SCC cancer cells contributes to generating an immunosuppressive and low TIIC microenvironment. Future studies are urged to demonstrate *TP63*/ΔNp63 expression as a candidate prognostic and predictive biomarker for ICB immunotherapy.

Compared to studies on TILs, tumor-infiltrating myeloid cells have received less attention. We specifically showed that depletion of ΔNp63 leads to the accumulation of tumor-suppressive reprogrammed myeloid cells in the TME of ESCC xenografts. MDSCs with increased MHCII expression were observed in ESCC xenografts upon cancer cell ΔNp63 depletion, consistently indicating reprogramming toward tumor suppression (53). Together with the regulations of cancer cell MHCI expression, we hypothesize that the regulation of myeloid recruitment and reprogramming by ΔNp63 mediate the influence of anticancer immunity in ESCC (Fig. 7E). Our analysis also suggests that more attention be paid to myeloid cell reprogramming and heterogeneous subgroup analysis in addition to general myeloid cell recruitment in future studies.

A previous study analyzing ICB resistance in melanoma patients showed that ICB-induced *F-Box and WD repeat domain containing 7* (*FBXW7*) inactivation repressed dsRNA sensing, IFN-I signaling induction, and MHCI expression in cancer cells (70). Inactivation of *FBXW7* also altered the tumor immune microenvironment, including decreased CD8^+^ T cell and macrophage infiltration. Our current findings phenocopy these data. Intriguing, FBXW7, as an E3-ubiquitin ligase, induces the degradation of p63/ΔNp63 (71), which is expressed in melanoma (72). Increased p63/ΔNp63 protein expression following *FBXW7* inactivation may, therefore, mediate the IFN-I signaling repression and TIIC reprogramming observed in melanoma patients, as we showed in the current study in ESCC. This further suggests a broader role of p63/ΔNp63 in regulating viral mimicry response and TIIC infiltration in other cancer types beyond SCCs.

Our multi-method analyses in *in vitro* cultures, xenograft models, and human tumor tissue samples demonstrate that ΔNp63 restricts endogenous retrotransposon expression and retrotransposon-induced viral mimicry response and modulates both the cancer cell and the TME. This suggests a translational potential to stratify ESCC and lung SCC patients for viral mimicry-boosting targeted therapy. ΔNp63 and druggable targets upstream of ΔNp63, regulating its expression and function (61), may also be considered in combination therapies. The biobankable panel of ESCC organoid cultures established will facilitate future studies.

## ACKNOWLEDGEMENTS

We acknowledge DSMZ (German Collection of Microorganisms and Cell Culture) for the KYSE cell lines. We thank Prof. Gopesh Srivastava and the late Prof. George Tsao for the cell lines. We acknowledge the University of Hong Kong Li Ka Shing Faculty of Medicine Centre for PanorOmic Sciences for providing flow cytometry, live-cell bioluminescence imaging facilities, and mycoplasma screening service. We acknowledge Food and Health Bureau Hong Kong (Health and Medical Research Fund 6171566 to VZY) and Research Grants Council Hong Kong (Theme-based Research Scheme T12-701/17-R to MLL) for funding support.

## AUTHOR CONTRIBUTIONS

Conceptualization: VZY, MLL

Methodology: VZY, SSS, BCL, GZH

Investigation: VZY, SSS, BCL, GZH, CWW, MKC, LKC, KX, ZZT, IYW, CLW, DKC, FSC, BTL, KL, AWL, AKL, DLK, SL, MLL

Visualization: VZY, SSS, CWW, LKC

Funding acquisition: VZY, MLL

Project administration: SL, MLL

Supervision: VZY, SL, MLL

Writing – original draft: VZY, SSS

Writing – review & editing: VZY, JMK, AWL, AKL, WD, MLL

