## Supplementary Figures for "ΔNp63-restricted viral mimicry response impedes cancer cell viability and remodels tumor microenvironment in esophageal squamous cell carcinoma"

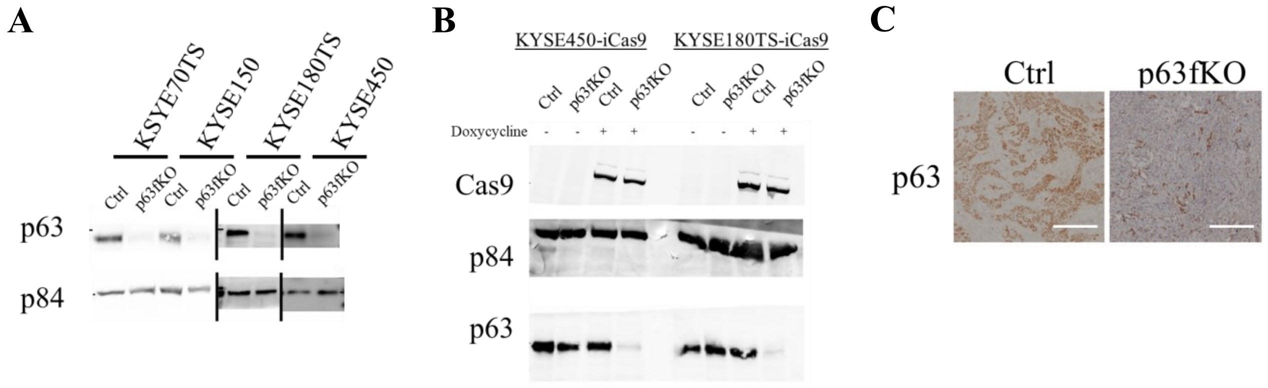


**Supplementary Figure 1. A**) WB analysis confirms the significant depletion of the endogenous ΔNp63 protein expressions in ESCC cell lines by CRISPR-mediated protein depletion. **B** and **C**) Verification of doxycycline-inducible ΔNp63 depletion in ESCC cell lines by *in vitro* WB analysis (B) and by IHC staining on CDXs (C, scale bar = 200μm ).

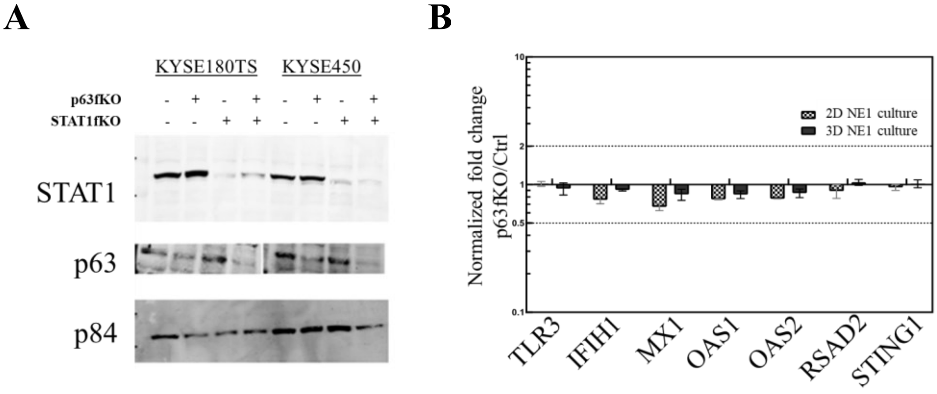


**Supplementary Figure 2. A**) WB analysis verifies the ΔNp63 and STAT1 depletions in ESCC cell lines. **B**) RNA expressions of type I IFN-related genes in NE1 cell line upon ΔNp63 depletion. Fold change (p63fKO/Ctrl) ≥ 2 indicates significant upregulation (upper dotted line), whereas fold change (p63fKO/Ctrl) ≤ 0.5 indicates significant downregulation (lower dotted line). None of the gene expressions was significantly altered.

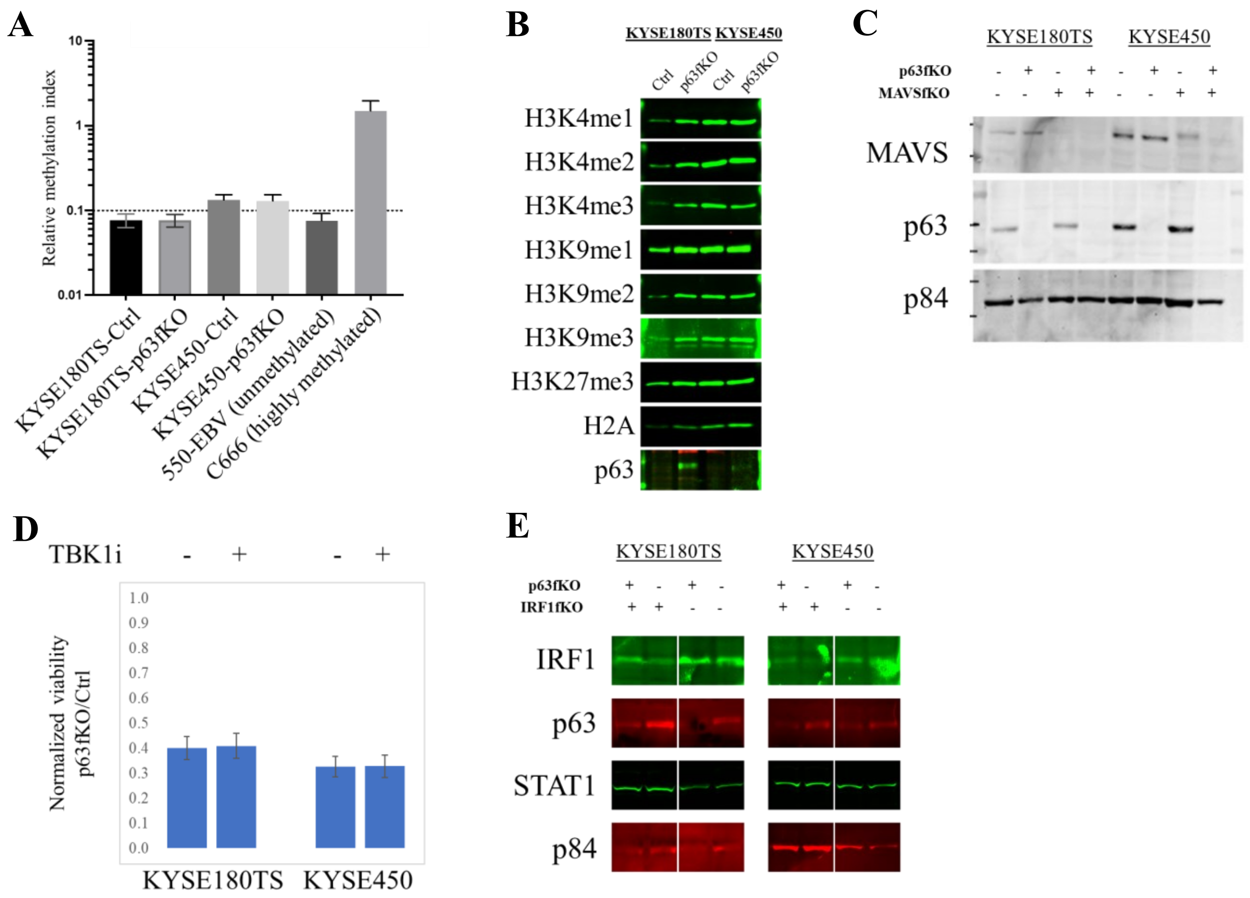


**Supplementary Figure 3. A**) Global genomic DNA methylation level indicated by LINE1 relative methylation level in ESCC cell lines upon ΔNp63 depletion with reference to the hypomethylated EBV infected 550 cell line and the hypermethylated C666 cell line (*84*). **B**) Levels of histone modification marks in ESCC cell lines upon p63 depletion by WB analysis. Expression of Histone H2A was used as loading control. **C**) WB analysis verifies the ΔNp63 and MAVS depletions in ESCC cell lines. **D**) TBK1 inhibition (TBKi) by small chemical molecule (MRT67307, 1μM) does not rescue the reduced viability upon p63 depletion. Shown are the normalized viability of ΔNp63-depleted cells to the corresponding control cells. **E**) WB analysis verifies the ΔNp63 and IRF1 depletions in ESCC cell lines.

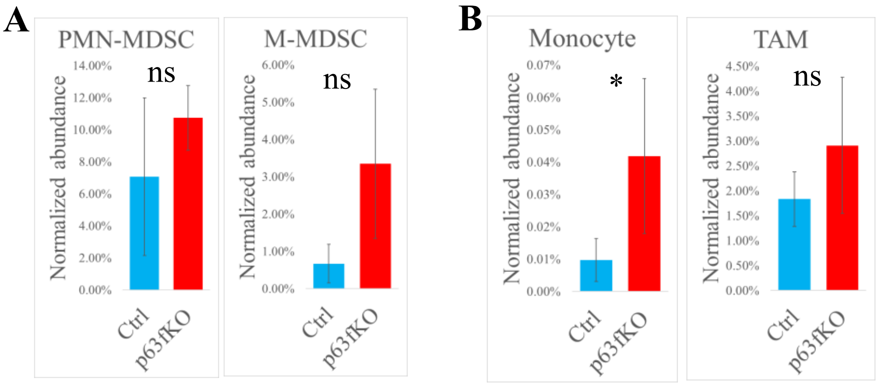


**Supplementary Figure 4. A**) Quantification of tumor-infiltrating M-MDSCs and PMN-MDSCs, as normalized to human cancer cells, in ΔNp63-depleted KSYE450 CDXs on nude mice. **B**) Quantification of tumor-infiltrating monocyte and TAMs, as normalized to human cancer cells, in ΔNp63-depleted KSYE450 CDXs on NOD/SCID mice. *, p-value<0.05; ns: not statistically significant.
