## Supplementary Tables for "ΔNp63-restricted viral mimicry response impedes cancer cell viability and remodels tumor microenvironment in esophageal squamous cell carcinoma"

Supplementary Table 1

Transcript-specific analysis of *TP63* variant expressions in hNEEO, PDO (PDO-ESCC and PDO-EAC), immortalized normal esophageal epithelial cell lines (2DCL-NE), ESCC cell lines (2DCL-ESCC), and three public ESCC transcriptomic datasets. TP63-203, encoding ΔNp63α, is the dominant variant expresssed. Other variants show minimal expressions.

| | | ***hNEEO vs PDO-ESCC*** | | | | | |
|---|---|---|---|---|---|---|---|
| Transcript ID | Encoded protein isoforms | P-value | FDR | Ratio | Fold change | LSMean(hNEEO) | LSMean(PDO-ESCC) |
| **TP63-201** | TAp63 alpha | 1.64E-01 | 4.24E-01 | 0.51 | -1.97 | 0.93 | 1.82 |
| **TP63-202** | TAp63 delta | 2.08E-01 | 4.89E-01 | 0.43 | -2.31 | 0.56 | 1.29 |
| **TP63-203** | **DeltaNp63 alpha** | **2.29E-01** | **5.18E-01** | **0.48** | **-2.09** | **259.25** | **540.59** |
| **TP63-205** | DeltaNp63 delta | 6.80E-02 | 2.43E-01 | 1.94 | 1.94 | 0.01 | 0.01 |
| **TP63-206** | DeltaNp63 beta | 3.46E-01 | 6.48E-01 | 0.44 | -2.28 | 0.05 | 0.11 |
| **TP63-207** | TAp63 gamma | 1.73E-01 | 4.40E-01 | 0.29 | -3.42 | 0.00 | 0.01 |
| **TP63-208** | unknown | 1.30E-02 | 7.64E-02 | 0.13 | -7.53 | 0.13 | 0.96 |
| **TP63-209** | DeltaNp63 gamma | 1.08E-01 | 3.29E-01 | 0.38 | -2.63 | 1.07 | 2.82 |
| **TP63-210** | DeltaNp63 epsilon | 4.06E-01 | 7.05E-01 | 0.51 | -1.95 | 0.02 | 0.05 |
| **TP63-211** | DeltaNp63 epsilon | 4.71E-01 | 7.59E-01 | 0.01 | -68.53 | 0.00 | 0.05 |
| **TP63-212** | DeltaNp63 epsilon | 3.41E-01 | 6.43E-01 | 0.35 | -2.83 | 22.48 | 63.63 |
| **TP63-213** | No protein | 5.61E-01 | 8.23E-01 | 0.51 | -1.98 | 4.98 | 9.86 |

| *hNEEO vs PDO-EAC* | | | | | | *2DCL-NE vs* | | |
|---|---|---|---|---|---|---|---|---|
| P-value | FDR | Ratio | Fold change | LSMean(hNEEO) | LSMean(PDO-EAC) | P-value | FDR | Ratio |
| 1.48E-12 | 6.68E-10 | 262.87 | 262.87 | 0.93 | 0.00 | 0.14 | 0.62 | 1.20 |
| 1.35E-02 | 1.18E-01 | 10.83 | 10.83 | 0.56 | 0.05 | 0.10 | 0.52 | 0.23 |
| **6.89E-25** | **3.02E-21** | **4528.42** | **4528.42** | **259.25** | **0.06** | **0.17** | **0.66** | **1.01** |
| 9.79E-04 | 1.81E-02 | 139.98 | 139.98 | 0.01 | 0.00 | 0.00 | 0.04 | 966.34 |
| 5.43E-02 | 2.89E-01 | 2.91 | 2.91 | 0.05 | 0.02 | 0.20 | 0.70 | 2.80 |
| 1.04E-02 | 9.94E-02 | 18.47 | 18.47 | 0.00 | 0.00 | 0.61 | 1.00 | 0.16 |
| 1.99E-02 | 1.53E-01 | 1268.51 | 1268.51 | 0.13 | 0.00 | 0.67 | 1.00 | 0.85 |
| 7.16E-13 | 3.55E-10 | 10737.61 | 10737.61 | 1.07 | 0.00 | 0.99 | 1.00 | 0.15 |
| 5.45E-04 | 1.17E-02 | 247.38 | 247.38 | 0.02 | 0.00 | 0.01 | 0.18 | 1.01 |
| 6.24E-01 | 9.59E-01 | 1.74 | 1.74 | 0.00 | 0.00 | 0.57 | 1.00 | 1.87 |
| 7.78E-13 | 3.81E-10 | 224409.10 | 224409.10 | 22.48 | 0.00 | 0.41 | 0.92 | 1.72 |
| 3.66E-09 | 6.42E-07 | 49812.79 | 49812.79 | 4.98 | 0.00 | 0.30 | 0.83 | 0.92 |

| *2DCL-ESCC* | | | *Patient normal tissue vs ESCC tissue (GSE111011)* | | | | | |
|---|---|---|---|---|---|---|---|---|
| Fold change | LSMean(2DCL-NE) | LSMean(2DCL-ESCC) | P-value | FDR | Ratio | Fold change | LSMean(Patient normal tissue ) | LSMean(Patient ESCC tissue ) |
| 1.20 | 8.12 | 6.75 | 3.93E-02 | 3.60E-01 | 0.18 | -5.43 | 1.80 | 9.76 |
| -4.42 | 0.22 | 0.95 | 2.59E-01 | 8.48E-01 | 0.24 | -4.20 | 0.11 | 0.44 |
| **1.01** | **186.26** | **185.06** | **1.60E-01** | **7.04E-01** | **0.32** | **-3.15** | **192.07** | **604.93** |
| 966.34 | 0.21 | 0.00 | 9.56E-01 | 1.00E+00 | 0.93 | -1.08 | 0.00 | 0.00 |
| 2.80 | 0.05 | 0.02 | 1.68E-01 | 7.18E-01 | 0.01 | -197.83 | 0.00 | 0.04 |
| -6.11 | 0.05 | 0.31 | 1.26E-02 | 1.91E-01 | 0.07 | -13.59 | 0.03 | 0.40 |
| -1.17 | 0.17 | 0.20 | 3.93E-01 | 9.58E-01 | 0.23 | -4.34 | 0.21 | 0.93 |
| -6.55 | 0.50 | 3.30 | 3.38E-01 | 9.23E-01 | 0.37 | -2.69 | 8.48 | 22.82 |
| 1.01 | 1.58 | 1.56 | 7.05E-03 | 1.35E-01 | 0.04 | -23.66 | 0.01 | 0.18 |
| 1.87 | 0.00 | 0.00 | 3.73E-02 | 3.51E-01 | 0.11 | -8.97 | 0.01 | 0.13 |
| 1.72 | 15.57 | 9.06 | 4.51E-01 | 9.84E-01 | 0.36 | -2.76 | 8.66 | 23.88 |
| -1.09 | 3.72 | 4.04 | 5.42E-01 | 1.00E+00 | 0.38 | -2.64 | 7.48 | 19.77 |

| *Patient normal tissue vs ESCC tissue (GSE149609)* | | | | | | *Patient normal tissue vs ES* | | |
|---|---|---|---|---|---|---|---|---|
| P-value | FDR | Ratio | Fold change | LSMean(Patient normal tissue ) | LSMean(Patient ESCC tissue ) | P-value | FDR | Ratio |
| 1.29E-01 | 5.23E-01 | 0.25 | -3.95 | 3.69 | 14.58 | 2.55E-01 | 6.04E-01 | 0.30 |
| 7.67E-01 | 9.97E-01 | 0.45 | -2.23 | 0.12 | 0.26 | 1.56E-01 | 4.71E-01 | 0.31 |
| **2.04E-01** | **6.44E-01** | **0.32** | **-3.16** | **118.08** | **372.83** | **2.61E-01** | **6.10E-01** | **0.33** |
| 9.95E-01 | 1.00E+00 | 1.00 | -1.00 | 0.00 | 0.00 | 1.11E-01 | 3.89E-01 | 0.01 |
| 4.41E-01 | 8.52E-01 | 0.00 | -369.81 | 0.00 | 0.04 | 9.42E-01 | 1.00E+00 | 0.58 |
| 1.78E-02 | 1.87E-01 | 0.18 | -5.43 | 0.00 | 0.02 | 2.41E-02 | 1.54E-01 | 0.26 |
| 2.64E-01 | 7.17E-01 | 0.18 | -5.48 | 0.05 | 0.27 | 2.99E-02 | 1.77E-01 | 0.07 |
| 3.48E-02 | 2.72E-01 | 0.19 | -5.24 | 1.25 | 6.54 | 1.36E-01 | 4.36E-01 | 0.27 |
| 3.55E-01 | 7.91E-01 | 0.28 | -3.58 | 0.00 | 0.01 | 3.47E-01 | 6.98E-01 | 0.39 |
| 4.14E-01 | 8.35E-01 | 0.01 | -177.57 | 0.00 | 0.02 | 1.00E+00 | 1.00E+00 | 1.00 |
| 5.29E-01 | 8.93E-01 | 0.24 | -4.21 | 1.91 | 8.06 | 8.04E-01 | 9.69E-01 | 0.36 |
| 8.29E-01 | 1.00E+00 | 0.44 | -2.30 | 0.82 | 1.88 | 3.59E-01 | 7.09E-01 | 0.28 |

| 'CC tissue (PRJNA298963) | | |
|---|---|---|
| Fold change | LSMean(Patient normal tissue ) | LSMean(Patient ESCC tissue ) |
| -3.32 | 4.77 | 15.80 |
| -3.19 | 0.03 | 0.09 |
| **-3.05** | **135.23** | **412.55** |
| -145.37 | 0.00 | 0.01 |
| -1.72 | 0.01 | 0.02 |
| -3.92 | 0.00 | 0.01 |
| -15.32 | 0.02 | 0.27 |
| -3.73 | 1.20 | 4.49 |
| -2.57 | 0.00 | 0.00 |
| 1.00 | 0.00 | 0.00 |
| -2.80 | 1.27 | 3.54 |
| -3.58 | 0.51 | 1.82 |

Supplementary Table 2

Summary of differentially-expressed genes upon ΔNp63 depletion in ESCC cell lines KYSE180 and KYSE450. IRF1 target genes are highlighted.

| Gene names | P-value (KYSE180-CTRL vs KYSE180-p63fKO) | FDR step up (KYSE180-CTRL vs KYSE180-p63fKO) | Ratio (KYSE180-CTRL vs KYSE180-p63fKO) | Fold change (KYSE180-CTRL vs KYSE180-p63fKO) | LSMean (KYSE180-CTRL) (KYSE180-CTRL vs KYSE180-p63fKO) | LSMean(KYSE180-p63fKO) (KYSE180-CTRL vs KYSE180-p63fKO) | P-value (KYSE450-CTRL vs KYSE450-p63fKO) | FDR step up (KYSE450-CTRL vs KYSE450-p63fKO) | Ratio (KYSE450-CTRL vs KYSE450-p63fKO) | Fold change (KYSE450-CTRL vs KYSE450-p63fKO) | LSMean (KYSE450-CTRL) (KYSE450-CTRL vs KYSE450-p63fKO) | LSMean(KYSE450-p63fKO) (KYSE450-CTRL vs KYSE450-p63fKO) |
|---|---|---|---|---|---|---|---|---|---|---|---|---|
| ***Up-regulated upon ΔNp63 depletion (FDR<0.05; Ratio<0.5; LSMean(p63fKO)>10)*** | | | | | | | | | | | | |
| MUC16 | 7.09E-12 | 6.63E-08 | 0.12 | -8.66 | 41.83 | 362.17 | 3.80E-14 | 1.07E-09 | 0.01 | -126.82 | 58.25 | 7386.92 |
| OAS2 | 2.28E-10 | 5.80E-07 | 0.28 | -3.61 | 189.72 | 685.54 | 8.70E-11 | 4.98E-07 | 0.23 | -4.44 | 131.83 | 585.16 |
| DDX58 | 7.73E-10 | 1.08E-06 | 0.20 | -4.98 | 22.13 | 110.27 | 9.44E-09 | 3.23E-06 | 0.34 | -2.97 | 114.71 | 341.20 |
| IFI44L | 9.48E-10 | 1.14E-06 | 0.28 | -3.57 | 69.23 | 247.41 | 1.61E-09 | 1.50E-06 | 0.31 | -3.23 | 155.58 | 503.13 |
| LAMP3 | 4.54E-09 | 2.83E-06 | 0.30 | -3.32 | 61.87 | 205.30 | 6.05E-08 | 8.57E-06 | 0.45 | -2.23 | 50.16 | 111.82 |
| WARS1 | 4.54E-09 | 2.83E-06 | 0.23 | -4.26 | 51.55 | 219.83 | 3.71E-09 | 2.26E-06 | 0.22 | -4.47 | 171.26 | 765.31 |
| ITGB8 | 4.15E-09 | 2.83E-06 | 0.43 | -2.34 | 306.16 | 717.76 | 3.96E-10 | 7.92E-07 | 0.29 | -3.41 | 147.57 | 502.85 |
| PARP14 | 7.17E-09 | 3.94E-06 | 0.40 | -2.48 | 269.94 | 670.11 | 2.07E-09 | 1.76E-06 | 0.33 | -3.01 | 108.01 | 325.44 |
| IGFBP5 | 9.39E-09 | 4.62E-06 | 0.14 | -7.03 | 30.11 | 211.62 | 6.40E-10 | 1.06E-06 | 0.05 | -19.20 | 7.33 | 140.67 |
| CCL5 | 1.03E-08 | 4.78E-06 | 0.16 | -6.21 | 8.61 | 53.48 | 3.08E-07 | 2.26E-05 | 0.34 | -2.93 | 11.03 | 32.33 |
| IFIT3 | 1.16E-08 | 5.01E-06 | 0.23 | -4.44 | 42.16 | 187.02 | 2.00E-08 | 4.45E-06 | 0.25 | -3.93 | 44.16 | 173.56 |
| ZNFX1 | 1.27E-08 | 5.31E-06 | 0.47 | -2.12 | 86.94 | 184.12 | 6.88E-09 | 2.78E-06 | 0.44 | -2.28 | 133.77 | 305.27 |
| IFI27 | 1.60E-08 | 6.24E-06 | 0.31 | -3.24 | 17.82 | 57.66 | 6.61E-10 | 1.06E-06 | 0.15 | -6.85 | 24.33 | 166.76 |
| ENC1 | 2.35E-08 | 8.29E-06 | 0.20 | -4.96 | 23.24 | 115.31 | 9.77E-09 | 3.26E-06 | 0.16 | -6.28 | 10.90 | 68.44 |
| HERC6 | 2.37E-08 | 8.29E-06 | 0.37 | -2.67 | 62.32 | 166.39 | 1.58E-09 | 1.50E-06 | 0.22 | -4.45 | 46.96 | 209.06 |
| ISG15 | 2.47E-08 | 8.34E-06 | 0.35 | -2.85 | 31.11 | 88.68 | 5.94E-09 | 2.70E-06 | 0.27 | -3.69 | 31.75 | 117.25 |
| IFIT2 | 2.63E-08 | 8.56E-06 | 0.28 | -3.62 | 17.49 | 63.31 | 1.49E-08 | 3.83E-06 | 0.25 | -4.07 | 33.89 | 138.01 |
| ABCC3 | 2.79E-08 | 8.89E-06 | 0.34 | -2.96 | 45.53 | 134.67 | 1.91E-08 | 4.33E-06 | 0.32 | -3.16 | 26.77 | 84.55 |
| PARP9 | 4.20E-08 | 1.20E-05 | 0.41 | -2.45 | 257.52 | 631.47 | 3.55E-08 | 6.13E-06 | 0.40 | -2.51 | 98.34 | 246.76 |
| SHROOM3 | 4.41E-08 | 1.22E-05 | 0.46 | -2.17 | 65.61 | 142.58 | 4.54E-08 | 7.28E-06 | 0.46 | -2.16 | 99.73 | 215.85 |
| NRP2 | 5.66E-08 | 1.38E-05 | 0.46 | -2.20 | 42.73 | 93.90 | 2.30E-09 | 1.79E-06 | 0.27 | -3.65 | 115.70 | 422.14 |
| FGF2 | 5.82E-08 | 1.40E-05 | 0.14 | -7.01 | 6.31 | 44.27 | 1.38E-07 | 1.38E-05 | 0.18 | -5.49 | 11.00 | 60.33 |
| MBOAT1 | 6.04E-08 | 1.44E-05 | 0.44 | -2.25 | 59.33 | 133.56 | 1.05E-09 | 1.20E-06 | 0.22 | -4.58 | 31.92 | 146.32 |
| IFIT1 | 6.78E-08 | 1.56E-05 | 0.35 | -2.88 | 82.41 | 237.55 | 4.60E-08 | 7.30E-06 | 0.32 | -3.08 | 73.78 | 227.49 |
| B4GALT5 | 7.10E-08 | 1.62E-05 | 0.46 | -2.18 | 137.28 | 299.23 | 7.22E-09 | 2.81E-06 | 0.33 | -3.04 | 107.85 | 327.65 |
| HEATR5A | 8.58E-08 | 1.83E-05 | 0.44 | -2.28 | 76.49 | 174.52 | 4.37E-09 | 2.37E-06 | 0.27 | -3.71 | 24.11 | 89.35 |
| PODXL | 9.90E-08 | 2.01E-05 | 0.42 | -2.36 | 53.23 | 125.74 | 3.29E-09 | 2.20E-06 | 0.23 | -4.30 | 10.62 | 45.72 |

| | | | | | | | | | | | | |
|---|---|---|---|---|---|---|---|---|---|---|---|---|
| ZDHHC21 | 1.02E-07 | 2.05E-05 | 0.40 | -2.52 | 21.11 | 53.13 | 3.21E-07 | 2.30E-05 | 0.46 | -2.16 | 48.42 | 104.78 |
| DDX60L | 1.08E-07 | 2.09E-05 | 0.39 | -2.56 | 29.89 | 76.66 | 7.21E-09 | 2.81E-06 | 0.24 | -4.20 | 24.42 | 102.66 |
| TNFRSF21 | 1.60E-07 | 2.63E-05 | 0.39 | -2.59 | 29.74 | 77.05 | 1.13E-09 | 1.22E-06 | 0.13 | -7.80 | 28.58 | 222.96 |
| SLFN5 | 1.62E-07 | 2.64E-05 | 0.47 | -2.14 | 190.51 | 407.40 | 8.64E-10 | 1.10E-06 | 0.18 | -5.55 | 62.58 | 347.21 |
| UNC5C | 1.73E-07 | 2.74E-05 | 0.16 | -6.09 | 5.22 | 31.83 | 2.76E-07 | 2.10E-05 | 0.19 | -5.36 | 7.85 | 42.03 |
| EPB41L4A | 1.73E-07 | 2.74E-05 | 0.33 | -3.00 | 34.93 | 104.77 | 4.79E-09 | 2.44E-06 | 0.15 | -6.81 | 3.47 | 23.63 |
| RSAD2 | 1.78E-07 | 2.78E-05 | 0.03 | -35.39 | 1.04 | 36.86 | 7.63E-07 | 4.10E-05 | 0.06 | -17.38 | 1.30 | 22.58 |
| PLSCR1 | 1.99E-07 | 2.99E-05 | 0.44 | -2.29 | 143.51 | 328.61 | 3.00E-08 | 5.65E-06 | 0.33 | -3.04 | 68.00 | 206.84 |
| OASL | 2.36E-07 | 3.31E-05 | 0.28 | -3.60 | 21.05 | 75.79 | 7.06E-08 | 9.34E-06 | 0.21 | -4.72 | 48.78 | 230.11 |
| CYBRD1 | 2.63E-07 | 3.53E-05 | 0.47 | -2.13 | 28.04 | 59.74 | 1.08E-08 | 3.34E-06 | 0.29 | -3.47 | 28.26 | 98.07 |
| TRIM22 | 3.25E-07 | 4.08E-05 | 0.17 | -5.85 | 6.78 | 39.66 | 6.53E-08 | 8.76E-06 | 0.10 | -9.78 | 11.76 | 115.06 |
| AGR2 | 3.33E-07 | 4.09E-05 | 0.13 | -7.92 | 4.42 | 34.98 | 1.13E-07 | 1.22E-05 | 0.08 | -11.78 | 49.69 | 585.45 |
| MUC4 | 3.46E-07 | 4.17E-05 | 0.25 | -3.97 | 6.33 | 25.12 | 3.79E-10 | 7.92E-07 | 0.02 | -52.94 | 3.05 | 161.60 |
| LGALS9 | 3.50E-07 | 4.17E-05 | 0.32 | -3.15 | 11.92 | 37.54 | 1.61E-06 | 6.59E-05 | 0.41 | -2.47 | 17.43 | 43.01 |
| SNTB1 | 3.50E-07 | 4.17E-05 | 0.19 | -5.26 | 8.90 | 46.81 | 9.66E-07 | 4.66E-05 | 0.24 | -4.12 | 8.52 | 35.05 |
| CMPK2 | 3.79E-07 | 4.35E-05 | 0.09 | -11.34 | 3.98 | 45.16 | 1.11E-06 | 5.14E-05 | 0.13 | -7.64 | 4.22 | 32.25 |
| CACNG4 | 3.91E-07 | 4.47E-05 | 0.21 | -4.78 | 4.52 | 21.62 | 3.19E-07 | 2.30E-05 | 0.20 | -5.01 | 7.25 | 36.34 |
| PLAAT3 | 3.96E-07 | 4.52E-05 | 0.32 | -3.15 | 20.60 | 64.94 | 8.56E-10 | 1.10E-06 | 0.05 | -19.79 | 1.87 | 36.98 |
| FMN1 | 4.33E-07 | 4.74E-05 | 0.30 | -3.35 | 9.84 | 32.94 | 2.79E-09 | 2.12E-06 | 0.07 | -14.05 | 6.29 | 88.34 |
| AL138689.2 | 4.29E-07 | 4.74E-05 | 0.28 | -3.56 | 10.59 | 37.71 | 2.43E-06 | 8.73E-05 | 0.38 | -2.63 | 4.29 | 11.26 |
| LMO7 | 4.58E-07 | 4.84E-05 | 0.39 | -2.56 | 26.25 | 67.30 | 6.95E-09 | 2.78E-06 | 0.16 | -6.08 | 35.23 | 214.19 |
| XAF1 | 4.56E-07 | 4.84E-05 | 0.14 | -7.18 | 21.35 | 153.21 | 1.62E-07 | 1.51E-05 | 0.10 | -10.08 | 6.66 | 67.17 |
| FREM2 | 5.03E-07 | 5.10E-05 | 0.40 | -2.51 | 25.18 | 63.28 | 3.80E-08 | 6.38E-06 | 0.25 | -3.97 | 18.04 | 71.68 |
| HERC5 | 5.06E-07 | 5.10E-05 | 0.47 | -2.12 | 36.17 | 76.72 | 6.28E-07 | 3.54E-05 | 0.48 | -2.07 | 87.99 | 182.06 |
| MACC1 | 5.99E-07 | 5.73E-05 | 0.46 | -2.16 | 81.27 | 175.62 | 2.18E-08 | 4.67E-06 | 0.27 | -3.65 | 24.93 | 90.97 |
| TSHZ2 | 6.14E-07 | 5.83E-05 | 0.40 | -2.51 | 30.65 | 77.09 | 8.18E-07 | 4.25E-05 | 0.41 | -2.42 | 22.73 | 55.00 |
| TMTC1 | 6.20E-07 | 5.84E-05 | 0.44 | -2.26 | 63.56 | 143.49 | 2.83E-08 | 5.63E-06 | 0.27 | -3.74 | 23.45 | 87.76 |
| PLXNA2 | 6.30E-07 | 5.89E-05 | 0.11 | -9.43 | 4.70 | 44.31 | 3.33E-06 | 1.08E-04 | 0.18 | -5.68 | 2.28 | 12.95 |
| LURAP1L | 6.58E-07 | 6.03E-05 | 0.42 | -2.36 | 19.87 | 46.94 | 3.39E-08 | 5.98E-06 | 0.26 | -3.91 | 37.18 | 145.32 |
| FOXA1 | 7.30E-07 | 6.45E-05 | 0.37 | -2.69 | 12.03 | 32.38 | 1.24E-07 | 1.28E-05 | 0.27 | -3.69 | 31.29 | 115.55 |
| TRANK1 | 8.16E-07 | 6.87E-05 | 0.41 | -2.43 | 29.31 | 71.36 | 1.88E-08 | 4.28E-06 | 0.20 | -4.95 | 9.55 | 47.32 |
| GAB2 | 9.96E-07 | 8.03E-05 | 0.20 | -5.12 | 5.48 | 28.08 | 4.89E-05 | 7.07E-04 | 0.42 | -2.40 | 11.62 | 27.85 |
| SRGAP1 | 1.04E-06 | 8.19E-05 | 0.47 | -2.14 | 54.93 | 117.79 | 1.10E-07 | 1.21E-05 | 0.34 | -2.96 | 70.62 | 208.72 |
| SEMA3A | 1.12E-06 | 8.62E-05 | 0.46 | -2.18 | 34.72 | 75.77 | 1.34E-08 | 3.76E-06 | 0.21 | -4.75 | 20.65 | 98.12 |
| HDAC9 | 1.19E-06 | 9.10E-05 | 0.39 | -2.54 | 4.11 | 10.45 | 3.94E-06 | 1.23E-04 | 0.46 | -2.16 | 48.69 | 105.38 |

| | | | | | | | | | | | | |
|---|---|---|---|---|---|---|---|---|---|---|---|---|
| PLAAT4 | 1.24E-06 | 9.39E-05 | 0.16 | -6.44 | 3.60 | 23.21 | 3.74E-08 | 6.33E-06 | 0.04 | -24.85 | 2.11 | 52.55 |
| SCG2 | 1.32E-06 | 9.79E-05 | 0.08 | -12.30 | 3.11 | 38.22 | 1.31E-06 | 5.78E-05 | 0.08 | -12.38 | 5.41 | 66.99 |
| DLX6-AS1 | 1.32E-06 | 9.79E-05 | 0.45 | -2.20 | 28.37 | 62.44 | 9.01E-08 | 1.08E-05 | 0.30 | -3.33 | 14.36 | 47.83 |
| BTC | 1.35E-06 | 9.92E-05 | 0.15 | -6.48 | 1.59 | 10.28 | 2.16E-06 | 8.06E-05 | 0.18 | -5.66 | 1.92 | 10.88 |
| NAALADL2 | 1.39E-06 | 1.00E-04 | 0.27 | -3.66 | 17.26 | 63.17 | 8.40E-07 | 4.31E-05 | 0.25 | -4.04 | 8.11 | 32.81 |
| GPAT3 | 1.46E-06 | 1.04E-04 | 0.44 | -2.26 | 29.17 | 65.98 | 3.52E-06 | 1.13E-04 | 0.49 | -2.03 | 9.34 | 19.00 |
| MYO5B | 1.55E-06 | 1.08E-04 | 0.31 | -3.27 | 8.75 | 28.61 | 5.42E-08 | 7.87E-06 | 0.14 | -7.40 | 15.96 | 118.04 |
| KRT4 | 1.73E-06 | 1.16E-04 | 0.12 | -8.37 | 1.71 | 14.31 | 2.34E-07 | 1.90E-05 | 0.05 | -18.51 | 4.32 | 79.94 |
| HECW2 | 1.73E-06 | 1.16E-04 | 0.07 | -13.66 | 1.10 | 14.97 | 5.55E-06 | 1.58E-04 | 0.11 | -8.90 | 1.56 | 13.86 |
| SAMD9L | 1.79E-06 | 1.19E-04 | 0.33 | -3.03 | 40.82 | 123.49 | 1.94E-08 | 4.36E-06 | 0.11 | -9.40 | 9.15 | 85.99 |
| SPDYE16 | 2.23E-06 | 1.39E-04 | 0.49 | -2.04 | 25.61 | 52.22 | 1.64E-06 | 6.67E-05 | 0.47 | -2.12 | 44.08 | 93.26 |
| OAS1 | 2.62E-06 | 1.54E-04 | 0.49 | -2.06 | 71.92 | 147.86 | 1.46E-07 | 1.44E-05 | 0.32 | -3.11 | 55.76 | 173.39 |
| NCOA7 | 2.65E-06 | 1.55E-04 | 0.40 | -2.53 | 17.66 | 44.64 | 2.48E-06 | 8.85E-05 | 0.39 | -2.56 | 68.41 | 175.13 |
| SAMHD1 | 2.83E-06 | 1.62E-04 | 0.47 | -2.12 | 44.78 | 94.88 | 5.07E-08 | 7.60E-06 | 0.24 | -4.10 | 19.13 | 78.41 |
| TRIB1 | 2.89E-06 | 1.63E-04 | 0.49 | -2.05 | 16.81 | 34.41 | 2.95E-06 | 9.93E-05 | 0.49 | -2.04 | 17.07 | 34.84 |
| LINC00472 | 2.88E-06 | 1.63E-04 | 0.38 | -2.60 | 29.22 | 75.93 | 1.24E-07 | 1.28E-05 | 0.21 | -4.79 | 15.72 | 75.34 |
| HPSE | 3.05E-06 | 1.70E-04 | 0.43 | -2.35 | 33.61 | 78.82 | 1.80E-06 | 7.08E-05 | 0.40 | -2.52 | 18.20 | 45.95 |
| PLEKHA7 | 3.13E-06 | 1.74E-04 | 0.38 | -2.65 | 19.04 | 50.42 | 1.41E-08 | 3.77E-06 | 0.10 | -9.69 | 5.98 | 57.90 |
| CENPT | 4.15E-06 | 2.06E-04 | 0.28 | -3.55 | 12.32 | 43.81 | 4.58E-05 | 6.72E-04 | 0.42 | -2.36 | 12.27 | 29.01 |
| SLC2A13 | 4.24E-06 | 2.08E-04 | 0.45 | -2.24 | 28.37 | 63.63 | 2.10E-06 | 7.90E-05 | 0.41 | -2.47 | 42.87 | 105.81 |
| AMOT | 4.37E-06 | 2.12E-04 | 0.41 | -2.41 | 5.08 | 12.27 | 1.65E-07 | 1.52E-05 | 0.23 | -4.38 | 8.08 | 35.40 |
| CDC42EP3 | 4.41E-06 | 2.13E-04 | 0.40 | -2.50 | 95.57 | 238.61 | 1.22E-06 | 5.48E-05 | 0.33 | -3.06 | 41.77 | 127.62 |
| FAR2P2 | 4.66E-06 | 2.20E-04 | 0.36 | -2.77 | 8.49 | 23.56 | 6.21E-06 | 1.71E-04 | 0.38 | -2.65 | 7.11 | 18.85 |
| LMO4 | 4.68E-06 | 2.20E-04 | 0.50 | -2.01 | 30.06 | 60.48 | 1.97E-06 | 7.57E-05 | 0.45 | -2.22 | 29.46 | 65.51 |
| MAP1B | 5.12E-06 | 2.37E-04 | 0.47 | -2.14 | 225.60 | 482.76 | 7.90E-07 | 4.18E-05 | 0.36 | -2.77 | 19.73 | 54.63 |
| MAP2 | 5.31E-06 | 2.39E-04 | 0.39 | -2.55 | 65.95 | 167.99 | 2.34E-07 | 1.90E-05 | 0.22 | -4.64 | 4.66 | 21.62 |
| PARP12 | 5.44E-06 | 2.43E-04 | 0.46 | -2.16 | 24.78 | 53.61 | 5.64E-07 | 3.26E-05 | 0.33 | -3.00 | 18.91 | 56.83 |
| LINC00641 | 6.17E-06 | 2.63E-04 | 0.42 | -2.35 | 25.79 | 60.73 | 4.91E-06 | 1.45E-04 | 0.41 | -2.44 | 14.38 | 35.03 |
| NEB | 9.45E-06 | 3.54E-04 | 0.26 | -3.82 | 9.34 | 35.67 | 1.38E-04 | 1.46E-03 | 0.42 | -2.38 | 16.49 | 39.26 |
| SH2D4A | 1.01E-05 | 3.70E-04 | 0.45 | -2.22 | 16.59 | 36.83 | 6.71E-06 | 1.81E-04 | 0.43 | -2.35 | 10.59 | 24.88 |
| DLX5 | 1.03E-05 | 3.72E-04 | 0.39 | -2.54 | 24.90 | 63.32 | 1.47E-06 | 6.20E-05 | 0.28 | -3.54 | 4.42 | 15.65 |
| LINC00342 | 1.10E-05 | 3.86E-04 | 0.23 | -4.26 | 3.03 | 12.90 | 2.80E-06 | 9.57E-05 | 0.16 | -6.08 | 12.45 | 75.72 |
| SGPP2 | 1.12E-05 | 3.89E-04 | 0.34 | -2.90 | 4.05 | 11.77 | 1.73E-06 | 6.89E-05 | 0.24 | -4.23 | 10.94 | 46.32 |
| AC092329.4 | 1.12E-05 | 3.89E-04 | 0.40 | -2.50 | 6.70 | 16.72 | 2.27E-06 | 8.32E-05 | 0.31 | -3.25 | 4.86 | 15.80 |
| ID3 | 1.18E-05 | 4.03E-04 | 0.38 | -2.64 | 18.86 | 49.78 | 2.96E-05 | 4.96E-04 | 0.43 | -2.31 | 10.60 | 24.51 |
| PLCE1 | 1.20E-05 | 4.07E-04 | 0.40 | -2.51 | 30.61 | 76.83 | 1.07E-05 | 2.47E-04 | 0.39 | -2.56 | 9.98 | 25.49 |

| | | | | | | | | | | | |
|---|---|---|---|---|---|---|---|---|---|---|---|
| CPEB4 | 1.27E-05 | 4.21E-04 | 0.48 | -2.08 | 20.99 | 43.57 | 1.19E-07 | 1.26E-05 | 0.22 | -4.58 | 16.49 | 75.50 |
| MYO5C | 1.42E-05 | 4.54E-04 | 0.43 | -2.34 | 41.22 | 96.52 | 6.77E-07 | 3.77E-05 | 0.25 | -3.97 | 7.47 | 29.71 |
| MX1 | 1.76E-05 | 5.31E-04 | 0.31 | -3.21 | 196.48 | 630.61 | 3.37E-04 | 2.82E-03 | 0.50 | -2.02 | 6.26 | 12.61 |
| SYNM | 1.87E-05 | 5.54E-04 | 0.46 | -2.18 | 13.57 | 29.57 | 9.98E-06 | 2.35E-04 | 0.42 | -2.36 | 16.17 | 38.21 |
| IFI35 | 1.91E-05 | 5.61E-04 | 0.44 | -2.30 | 23.29 | 53.49 | 5.88E-06 | 1.65E-04 | 0.36 | -2.75 | 12.19 | 33.54 |
| KRT80 | 1.95E-05 | 5.71E-04 | 0.18 | -5.48 | 14.45 | 79.20 | 1.40E-07 | 1.40E-05 | 0.03 | -39.60 | 1.62 | 64.22 |
| PIK3R3 | 2.16E-05 | 6.06E-04 | 0.39 | -2.58 | 5.98 | 15.43 | 1.27E-05 | 2.77E-04 | 0.36 | -2.82 | 16.35 | 46.03 |
| SP110 | 2.21E-05 | 6.13E-04 | 0.39 | -2.57 | 23.38 | 60.20 | 6.95E-05 | 9.06E-04 | 0.46 | -2.17 | 17.38 | 37.76 |
| ARHGAP27P1-BPTFP1-KPNA2P3 | 2.25E-05 | 6.19E-04 | 0.46 | -2.17 | 7.29 | 15.79 | 1.19E-05 | 2.65E-04 | 0.42 | -2.36 | 7.34 | 17.31 |
| AL137782.1 | 2.46E-05 | 6.52E-04 | 0.35 | -2.87 | 3.70 | 10.60 | 1.71E-06 | 6.85E-05 | 0.20 | -5.07 | 6.46 | 32.73 |
| USP18 | 2.47E-05 | 6.52E-04 | 0.37 | -2.70 | 12.63 | 34.11 | 2.57E-05 | 4.52E-04 | 0.38 | -2.66 | 11.32 | 30.16 |
| MIR99AHG | 2.60E-05 | 6.77E-04 | 0.33 | -3.00 | 13.97 | 41.93 | 1.54E-04 | 1.58E-03 | 0.44 | -2.27 | 11.71 | 26.64 |
| RDH10 | 2.62E-05 | 6.80E-04 | 0.45 | -2.21 | 13.45 | 29.69 | 4.51E-08 | 7.28E-06 | 0.12 | -8.64 | 6.35 | 54.85 |
| LYPD1 | 2.67E-05 | 6.89E-04 | 0.39 | -2.55 | 6.91 | 17.59 | 1.30E-06 | 5.74E-05 | 0.22 | -4.54 | 2.46 | 11.16 |
| SECTM1 | 2.73E-05 | 6.94E-04 | 0.48 | -2.10 | 6.84 | 14.35 | 7.21E-08 | 9.49E-06 | 0.15 | -6.68 | 3.32 | 22.16 |
| CICP27 | 3.22E-05 | 7.75E-04 | 0.38 | -2.66 | 6.46 | 17.19 | 2.29E-05 | 4.20E-04 | 0.36 | -2.80 | 6.67 | 18.69 |
| HSH2D | 3.28E-05 | 7.81E-04 | 0.47 | -2.14 | 10.75 | 22.96 | 8.07E-08 | 1.02E-05 | 0.14 | -7.11 | 3.15 | 22.42 |
| SMAD6 | 3.71E-05 | 8.52E-04 | 0.39 | -2.56 | 15.79 | 40.38 | 1.38E-05 | 2.95E-04 | 0.33 | -3.04 | 5.03 | 15.30 |
| TMC5 | 4.03E-05 | 8.94E-04 | 0.28 | -3.51 | 4.21 | 14.77 | 3.31E-08 | 5.98E-06 | 0.02 | -45.92 | 1.34 | 61.61 |
| CICP14 | 4.04E-05 | 8.95E-04 | 0.39 | -2.60 | 3.97 | 10.31 | 3.50E-06 | 1.12E-04 | 0.24 | -4.14 | 4.34 | 17.95 |
| NLRC5 | 4.07E-05 | 8.97E-04 | 0.37 | -2.70 | 10.75 | 28.97 | 3.29E-05 | 5.32E-04 | 0.35 | -2.83 | 7.84 | 22.14 |
| APOL6 | 4.53E-05 | 9.69E-04 | 0.48 | -2.10 | 83.93 | 176.10 | 5.36E-08 | 7.83E-06 | 0.12 | -8.57 | 8.02 | 68.70 |
| PPM1L | 5.45E-05 | 1.09E-03 | 0.47 | -2.11 | 11.72 | 24.77 | 5.17E-06 | 1.50E-04 | 0.34 | -2.98 | 17.88 | 53.31 |
| IER5L | 5.78E-05 | 1.15E-03 | 0.49 | -2.03 | 18.47 | 37.44 | 1.13E-06 | 5.19E-05 | 0.26 | -3.81 | 15.88 | 60.52 |
| MX2 | 6.43E-05 | 1.25E-03 | 0.35 | -2.83 | 18.46 | 52.29 | 4.28E-06 | 1.31E-04 | 0.19 | -5.15 | 20.94 | 107.85 |
| NBEA | 6.96E-05 | 1.32E-03 | 0.49 | -2.04 | 11.07 | 22.61 | 2.34E-05 | 4.25E-04 | 0.42 | -2.37 | 16.13 | 38.16 |
| CASZ1 | 8.55E-05 | 1.51E-03 | 0.46 | -2.19 | 27.84 | 61.03 | 3.27E-07 | 2.32E-05 | 0.15 | -6.75 | 2.49 | 16.80 |
| MISP | 1.05E-04 | 1.73E-03 | 0.46 | -2.16 | 19.87 | 42.88 | 5.85E-07 | 3.33E-05 | 0.17 | -5.92 | 8.64 | 51.11 |
| MYEF2 | 1.16E-04 | 1.84E-03 | 0.49 | -2.04 | 18.85 | 38.51 | 1.02E-04 | 1.19E-03 | 0.48 | -2.06 | 4.91 | 10.12 |
| MIR4435-2HG | 1.22E-04 | 1.90E-03 | 0.48 | -2.08 | 42.60 | 88.47 | 4.86E-06 | 1.44E-04 | 0.29 | -3.47 | 39.13 | 135.71 |
| AC025171.2 | 1.25E-04 | 1.93E-03 | 0.49 | -2.04 | 9.57 | 19.53 | 3.78E-05 | 5.89E-04 | 0.42 | -2.39 | 4.38 | 10.45 |
| IPO5P1 | 1.80E-04 | 2.49E-03 | 0.37 | -2.68 | 4.46 | 11.94 | 3.09E-04 | 2.64E-03 | 0.41 | -2.41 | 5.04 | 12.16 |
| APOL1 | 2.13E-04 | 2.81E-03 | 0.34 | -2.90 | 5.77 | 16.75 | 9.06E-06 | 2.21E-04 | 0.17 | -6.01 | 2.31 | 13.87 |
| CEACAM1 | 2.60E-04 | 3.22E-03 | 0.19 | -5.32 | 3.24 | 17.22 | 3.43E-05 | 5.48E-04 | 0.10 | -9.71 | 3.25 | 31.55 |
| CLEC7A | 2.83E-04 | 3.40E-03 | 0.44 | -2.27 | 6.73 | 15.29 | 5.40E-06 | 1.55E-04 | 0.21 | -4.76 | 5.31 | 25.31 |
| ZNF117 | 3.06E-04 | 3.58E-03 | 0.49 | -2.03 | 5.44 | 11.01 | 1.54E-06 | 6.44E-05 | 0.18 | -5.42 | 4.71 | 25.51 |
| ZSCAN31 | 3.12E-04 | 3.63E-03 | 0.40 | -2.50 | 6.33 | 15.80 | 3.24E-04 | 2.74E-03 | 0.40 | -2.49 | 6.66 | 16.62 |
| ABCA1 | 3.22E-04 | 3.71E-03 | 0.49 | -2.05 | 27.63 | 56.50 | 9.77E-08 | 1.12E-05 | 0.07 | -13.69 | 1.04 | 14.26 |
| GBP4 | 3.60E-04 | 3.99E-03 | 0.33 | -3.00 | 10.80 | 32.42 | 1.55E-05 | 3.19E-04 | 0.16 | -6.30 | 5.20 | 32.77 |

| | | | | | | | | | | | |
|---|---|---|---|---|---|---|---|---|---|---|---|
| RTKN2 | 3.61E-04 | 4.00E-03 | 0.42 | -2.38 | 12.64 | 30.03 | 8.31E-04 | 5.53E-03 | 0.47 | -2.14 | 4.75 | 10.16 |
| CD24 | 3.66E-04 | 4.03E-03 | 0.37 | -2.67 | 29.32 | 78.25 | 3.93E-07 | 2.55E-05 | 0.05 | -18.94 | 5.86 | 110.90 |
| KRT16 | 8.37E-04 | 7.27E-03 | 0.43 | -2.32 | 10.93 | 25.38 | 2.73E-06 | 9.38E-05 | 0.11 | -9.30 | 2.74 | 25.43 |
| IGSF10 | 1.31E-03 | 1.01E-02 | 0.40 | -2.50 | 23.38 | 58.51 | 3.30E-05 | 5.33E-04 | 0.18 | -5.44 | 3.38 | 18.39 |
| BSPRY | 1.52E-03 | 1.13E-02 | 0.38 | -2.66 | 4.95 | 13.20 | 1.41E-04 | 1.48E-03 | 0.23 | -4.26 | 3.98 | 16.94 |
| ASAH1 | 1.79E-03 | 1.28E-02 | 0.46 | -2.18 | 6.14 | 13.38 | 3.94E-04 | 3.13E-03 | 0.37 | -2.70 | 8.77 | 23.67 |
| GCNT2 | 2.79E-03 | 1.78E-02 | 0.49 | -2.05 | 8.18 | 16.79 | 9.86E-06 | 2.33E-04 | 0.16 | -6.43 | 2.52 | 16.19 |
| TIGAR | 4.55E-03 | 2.57E-02 | 0.44 | -2.30 | 6.41 | 14.72 | 1.89E-03 | 1.06E-02 | 0.38 | -2.62 | 4.82 | 12.61 |
| SULF2 | 5.69E-03 | 3.03E-02 | 0.29 | -3.43 | 4.24 | 14.57 | 3.54E-04 | 2.91E-03 | 0.13 | -7.57 | 2.32 | 17.54 |
| ***Down-regulated upon ΔNp63 depletion (FDR<0.05; Ratio>2; LSMean(ctrl)>10)*** | | | | | | | | | | | | |
| FAT2 | 3.01E-12 | 4.22E-08 | 7.99 | 7.99 | 374.48 | 46.84 | 1.38E-10 | 6.44E-07 | 3.16 | 3.16 | 1542.65 | 488.06 |
| DDIT4 | 3.69E-11 | 2.58E-07 | 6.59 | 6.59 | 757.35 | 114.93 | 1.07E-09 | 1.20E-06 | 3.06 | 3.06 | 1013.21 | 330.91 |
| TP63 | 1.13E-10 | 5.00E-07 | 4.29 | 4.29 | 731.16 | 170.27 | 1.15E-08 | 3.40E-06 | 2.04 | 2.04 | 448.74 | 220.20 |
| IER3 | 2.03E-10 | 5.70E-07 | 7.29 | 7.29 | 455.11 | 62.44 | 1.30E-09 | 1.30E-06 | 4.44 | 4.44 | 410.59 | 92.53 |
| EGFR | 5.16E-10 | 1.03E-06 | 2.23 | 2.23 | 3509.62 | 1577.35 | 1.29E-09 | 1.30E-06 | 2.00 | 2.00 | 7443.32 | 3718.02 |
| NECTIN1 | 7.53E-10 | 1.08E-06 | 3.78 | 3.78 | 1495.11 | 395.35 | 8.63E-10 | 1.10E-06 | 3.68 | 3.68 | 1148.96 | 312.48 |
| SFN | 8.48E-10 | 1.13E-06 | 3.36 | 3.36 | 219.12 | 65.25 | 1.48E-08 | 3.83E-06 | 2.18 | 2.18 | 107.52 | 49.39 |
| IRF6 | 9.73E-10 | 1.14E-06 | 3.08 | 3.08 | 303.28 | 98.50 | 8.87E-09 | 3.20E-06 | 2.22 | 2.22 | 318.34 | 143.28 |
| KRT5 | 1.28E-09 | 1.38E-06 | 2.12 | 2.12 | 1629.16 | 767.62 | 7.37E-10 | 1.09E-06 | 2.27 | 2.27 | 3054.38 | 1345.53 |
| WASF2 | 2.05E-09 | 1.98E-06 | 2.44 | 2.44 | 555.99 | 227.40 | 2.14E-09 | 1.77E-06 | 2.43 | 2.43 | 733.48 | 301.91 |
| IL6R | 2.10E-09 | 1.98E-06 | 4.67 | 4.67 | 150.97 | 32.32 | 1.44E-08 | 3.80E-06 | 3.14 | 3.14 | 43.44 | 13.84 |
| HAS3 | 2.78E-09 | 2.34E-06 | 7.23 | 7.23 | 119.08 | 16.46 | 9.27E-09 | 3.21E-06 | 5.16 | 5.16 | 241.28 | 46.78 |
| ADM | 3.47E-09 | 2.70E-06 | 8.43 | 8.43 | 101.37 | 12.03 | 7.42E-08 | 9.70E-06 | 3.77 | 3.77 | 359.34 | 95.44 |
| PTGS2 | 4.21E-09 | 2.83E-06 | 3.35 | 3.35 | 50.17 | 15.00 | 5.96E-09 | 2.70E-06 | 3.14 | 3.14 | 806.68 | 257.00 |
| CA9 | 7.58E-09 | 4.08E-06 | 16.50 | 16.50 | 96.53 | 5.85 | 5.86E-06 | 1.64E-04 | 2.70 | 2.70 | 499.40 | 185.30 |
| FRMD6 | 1.37E-08 | 5.65E-06 | 2.61 | 2.61 | 195.98 | 74.95 | 6.64E-09 | 2.78E-06 | 2.93 | 2.93 | 222.46 | 75.93 |
| SESN3 | 1.42E-08 | 5.78E-06 | 2.38 | 2.38 | 495.72 | 208.32 | 1.02E-08 | 3.26E-06 | 2.49 | 2.49 | 1303.00 | 522.72 |
| MYC | 2.77E-08 | 8.89E-06 | 2.41 | 2.41 | 224.13 | 92.81 | 6.72E-09 | 2.78E-06 | 3.00 | 3.00 | 562.01 | 187.22 |
| SERPINE1 | 3.33E-08 | 1.02E-05 | 18.08 | 18.08 | 87.09 | 4.82 | 2.56E-07 | 2.01E-05 | 8.14 | 8.14 | 18.75 | 2.30 |
| S100A2 | 4.59E-08 | 1.24E-05 | 2.98 | 2.98 | 165.30 | 55.38 | 2.44E-07 | 1.94E-05 | 2.32 | 2.32 | 56.79 | 24.45 |
| CNTN1 | 4.74E-08 | 1.27E-05 | 2.63 | 2.63 | 414.63 | 157.78 | 1.50E-07 | 1.45E-05 | 2.24 | 2.24 | 202.46 | 90.45 |
| ADAMTS1 | 7.37E-08 | 1.67E-05 | 4.05 | 4.05 | 60.27 | 14.86 | 1.96E-07 | 1.72E-05 | 3.33 | 3.33 | 48.69 | 14.63 |
| AHDC1 | 8.46E-08 | 1.83E-05 | 2.90 | 2.90 | 70.55 | 24.33 | 5.23E-07 | 3.09E-05 | 2.23 | 2.23 | 96.27 | 43.18 |
| CYP2S1 | 9.13E-08 | 1.90E-05 | 3.82 | 3.82 | 114.32 | 29.91 | 2.94E-07 | 2.20E-05 | 3.06 | 3.06 | 91.91 | 30.06 |
| FSCN1 | 1.09E-07 | 2.10E-05 | 2.77 | 2.77 | 574.25 | 207.49 | 1.19E-06 | 5.41E-05 | 2.01 | 2.01 | 343.43 | 170.73 |
| PGF | 1.42E-07 | 2.44E-05 | 17.93 | 17.93 | 37.67 | 2.10 | 1.47E-06 | 6.21E-05 | 7.42 | 7.42 | 30.20 | 4.07 |
| GBP6 | 1.59E-07 | 2.63E-05 | 2.71 | 2.71 | 101.21 | 37.39 | 7.58E-08 | 9.70E-06 | 3.05 | 3.05 | 90.33 | 29.57 |
| COL7A1 | 2.02E-07 | 2.99E-05 | 3.80 | 3.80 | 21.15 | 5.56 | 1.05E-06 | 4.93E-05 | 2.81 | 2.81 | 75.24 | 26.80 |
| GPC1 | 2.32E-07 | 3.30E-05 | 2.23 | 2.23 | 204.46 | 91.73 | 2.12E-07 | 1.79E-05 | 2.25 | 2.25 | 248.35 | 110.27 |
| OLFML2A | 2.34E-07 | 3.31E-05 | 2.21 | 2.21 | 152.59 | 69.05 | 1.48E-08 | 3.83E-06 | 3.38 | 3.38 | 36.75 | 10.86 |
| MFHAS1 | 2.59E-07 | 3.51E-05 | 3.43 | 3.43 | 128.52 | 37.52 | 3.66E-07 | 2.48E-05 | 3.22 | 3.22 | 75.30 | 23.42 |
| PER1 | 3.09E-07 | 3.93E-05 | 2.80 | 2.80 | 37.25 | 13.32 | 4.38E-07 | 2.70E-05 | 2.65 | 2.65 | 134.52 | 50.80 |
| FABP5 | 3.35E-07 | 4.09E-05 | 2.63 | 2.63 | 50.23 | 19.11 | 2.34E-06 | 8.51E-05 | 2.04 | 2.04 | 41.96 | 20.60 |
| TNFAIP8 | 3.78E-07 | 4.35E-05 | 2.07 | 2.07 | 49.64 | 23.92 | 1.62E-08 | 3.83E-06 | 3.29 | 3.29 | 93.29 | 28.32 |

| | | | | | | | | | | | |
|---|---|---|---|---|---|---|---|---|---|---|---|
| ARL4C | 4.60E-07 | 4.84E-05 | 2.18 | 2.18 | 30.07 | 13.80 | 5.64E-08 | 8.14E-06 | 2.95 | 2.95 | 242.13 | 82.16 |
| ARL4D | 4.72E-07 | 4.88E-05 | 4.86 | 4.86 | 73.12 | 15.05 | 8.84E-06 | 2.18E-04 | 2.68 | 2.68 | 85.91 | 32.04 |
| CEL | 9.08E-07 | 7.53E-05 | 83.64 | 83.64 | 93.17 | 1.11 | 1.26E-04 | 1.37E-03 | 7.70 | 7.70 | 22.53 | 2.93 |
| NGFR | 9.76E-07 | 7.93E-05 | 8.81 | 8.81 | 12.76 | 1.45 | 4.70E-05 | 6.87E-04 | 3.18 | 3.18 | 31.92 | 10.04 |
| MMP10 | 1.50E-06 | 1.05E-04 | 9.00 | 9.00 | 44.98 | 5.00 | 1.94E-04 | 1.88E-03 | 2.74 | 2.74 | 18.64 | 6.80 |
| CDH13 | 1.84E-06 | 1.21E-04 | 2.50 | 2.50 | 54.05 | 21.58 | 1.21E-06 | 5.47E-05 | 2.66 | 2.66 | 30.02 | 11.29 |
| PHLDB2 | 1.94E-06 | 1.25E-04 | 4.22 | 4.22 | 93.55 | 22.19 | 3.63E-06 | 1.15E-04 | 3.66 | 3.66 | 73.30 | 20.02 |
| ANXA8 | 1.96E-06 | 1.26E-04 | 2.34 | 2.34 | 33.45 | 14.30 | 9.01E-07 | 4.52E-05 | 2.61 | 2.61 | 42.36 | 16.26 |
| SNAI2 | 2.02E-06 | 1.29E-04 | 2.99 | 2.99 | 36.41 | 12.19 | 3.74E-07 | 2.49E-05 | 4.16 | 4.16 | 31.41 | 7.55 |
| LGALS1 | 2.31E-06 | 1.41E-04 | 4.25 | 4.25 | 81.56 | 19.21 | 1.00E-05 | 2.36E-04 | 3.13 | 3.13 | 18.51 | 5.91 |
| USP31 | 2.54E-06 | 1.51E-04 | 2.37 | 2.37 | 99.98 | 42.19 | 6.75E-07 | 3.77E-05 | 2.90 | 2.90 | 122.63 | 42.34 |
| NCS1 | 3.21E-06 | 1.77E-04 | 2.90 | 2.90 | 69.60 | 23.99 | 1.11E-05 | 2.54E-04 | 2.40 | 2.40 | 74.21 | 30.95 |
| SIK1B | 3.25E-06 | 1.78E-04 | 2.14 | 2.14 | 49.59 | 23.21 | 7.84E-07 | 4.17E-05 | 2.58 | 2.58 | 27.11 | 10.50 |
| NUAK2 | 3.70E-06 | 1.92E-04 | 2.85 | 2.85 | 29.70 | 10.42 | 2.86E-06 | 9.68E-05 | 2.97 | 2.97 | 15.48 | 5.22 |
| BTG2 | 3.94E-06 | 2.02E-04 | 2.93 | 2.93 | 19.44 | 6.64 | 4.63E-05 | 6.77E-04 | 2.07 | 2.07 | 18.49 | 8.94 |
| DKK3 | 4.09E-06 | 2.04E-04 | 11.41 | 11.41 | 25.31 | 2.22 | 1.59E-05 | 3.25E-04 | 7.16 | 7.16 | 64.53 | 9.01 |
| CLIP2 | 4.12E-06 | 2.05E-04 | 2.41 | 2.41 | 76.36 | 31.66 | 9.31E-07 | 4.61E-05 | 3.04 | 3.04 | 68.56 | 22.57 |
| GAL | 5.64E-06 | 2.48E-04 | 2.17 | 2.17 | 24.81 | 11.46 | 2.12E-06 | 7.94E-05 | 2.46 | 2.46 | 16.94 | 6.88 |
| DUSP9 | 5.83E-06 | 2.53E-04 | 3.71 | 3.71 | 18.21 | 4.91 | 4.38E-06 | 1.32E-04 | 3.96 | 3.96 | 13.17 | 3.33 |
| EFNB1 | 7.62E-06 | 3.06E-04 | 2.65 | 2.65 | 49.32 | 18.59 | 3.40E-05 | 5.44E-04 | 2.16 | 2.16 | 16.53 | 7.67 |
| DLK2 | 8.94E-06 | 3.40E-04 | 3.95 | 3.95 | 19.92 | 5.05 | 2.50E-05 | 4.43E-04 | 3.22 | 3.22 | 11.02 | 3.43 |
| ANXA8L1 | 1.07E-05 | 3.81E-04 | 2.50 | 2.50 | 26.81 | 10.71 | 9.58E-06 | 2.29E-04 | 2.55 | 2.55 | 55.72 | 21.81 |
| RAB30 | 1.11E-05 | 3.89E-04 | 2.95 | 2.95 | 26.28 | 8.90 | 6.91E-07 | 3.83E-05 | 5.38 | 5.38 | 116.89 | 21.72 |
| LPAR3 | 1.21E-05 | 4.11E-04 | 2.55 | 2.55 | 25.30 | 9.94 | 2.87E-05 | 4.86E-04 | 2.26 | 2.26 | 28.47 | 12.61 |
| SOCS2 | 1.99E-05 | 5.75E-04 | 2.09 | 2.09 | 55.56 | 26.60 | 5.45E-06 | 1.57E-04 | 2.48 | 2.48 | 82.02 | 33.09 |
| IGFBP7 | 2.81E-05 | 7.04E-04 | 2.73 | 2.73 | 28.93 | 10.59 | 5.83E-06 | 1.64E-04 | 3.63 | 3.63 | 24.10 | 6.64 |
| ATP8B3 | 4.05E-05 | 8.96E-04 | 3.76 | 3.76 | 28.60 | 7.60 | 2.15E-04 | 2.03E-03 | 2.71 | 2.71 | 13.69 | 5.05 |
| GPC3 | 4.63E-05 | 9.81E-04 | 2.11 | 2.11 | 13.18 | 6.26 | 3.96E-05 | 6.06E-04 | 2.13 | 2.13 | 73.02 | 34.24 |
| RGS2 | 7.81E-05 | 1.42E-03 | 2.29 | 2.29 | 11.43 | 4.99 | 1.21E-05 | 2.67E-04 | 3.07 | 3.07 | 16.93 | 5.52 |
| PLA2G4A | 1.03E-04 | 1.70E-03 | 2.16 | 2.16 | 19.18 | 8.88 | 1.21E-05 | 2.67E-04 | 2.94 | 2.94 | 56.54 | 19.25 |
| TNC | 1.44E-04 | 2.14E-03 | 2.71 | 2.71 | 57.82 | 21.33 | 3.95E-05 | 6.04E-04 | 3.49 | 3.49 | 43.78 | 12.53 |
| BNC1 | 1.48E-04 | 2.18E-03 | 3.30 | 3.30 | 19.70 | 5.97 | 2.19E-04 | 2.05E-03 | 3.14 | 3.14 | 37.76 | 12.04 |
| IKBIP | 1.51E-04 | 2.20E-03 | 2.06 | 2.06 | 28.42 | 13.81 | 1.30E-04 | 1.40E-03 | 2.09 | 2.09 | 23.44 | 11.23 |
| CNTNAP1 | 4.41E-04 | 4.58E-03 | 2.08 | 2.08 | 21.64 | 10.39 | 4.84E-05 | 7.01E-04 | 2.86 | 2.86 | 14.88 | 5.21 |
| AC009119.1 | 4.83E-04 | 4.84E-03 | 2.63 | 2.63 | 21.73 | 8.25 | 1.77E-04 | 1.76E-03 | 3.28 | 3.28 | 18.82 | 5.73 |
| RIN1 | 8.43E-04 | 7.30E-03 | 2.17 | 2.17 | 12.16 | 5.60 | 1.43E-03 | 8.48E-03 | 2.06 | 2.06 | 21.40 | 10.37 |
| TCF4 | 1.34E-03 | 1.04E-02 | 2.04 | 2.04 | 10.43 | 5.11 | 5.40E-04 | 3.97E-03 | 2.34 | 2.34 | 39.70 | 16.94 |
| RCOR2 | 1.49E-03 | 1.12E-02 | 2.68 | 2.68 | 16.39 | 6.12 | 7.39E-04 | 5.06E-03 | 3.13 | 3.13 | 11.85 | 3.79 |
| NPW | 2.37E-03 | 1.58E-02 | 2.43 | 2.43 | 10.43 | 4.30 | 4.13E-03 | 1.97E-02 | 2.23 | 2.23 | 14.28 | 6.42 |

Supplementary Table 3

Gene ontology annotations by GeneMANIA from differentially-expressed genes upon ΔNp63 depletion in ESCC cell lines KYSE180 and KYSE450. Highlighted are annotations related to IFN-I signaling.

| *False discovery rate* | *Coverage* | *Gene ontology annotation* |
|---|---|---|
| | | ***Input: up-regulated genes*** |
| 6.4E-29 | 25/88 | cellular response to type I interferon |
| 6.4E-29 | 25/88 | response to type I interferon |
| 4.7E-21 | 26/197 | response to virus |
| 9E-16 | 20/149 | regulation of viral life cycle |
| 2.4E-15 | 22/210 | regulation of viral process |
| 6.2E-15 | 22/221 | regulation of symbiotic process |
| 6.7E-15 | 17/102 | negative regulation of viral process |
| 4.3E-14 | 16/94 | regulation of viral genome replication |
| 2.8E-13 | 16/106 | viral genome replication |
| 1.1E-12 | 17/139 | response to interferon-gamma |
| 2.6E-12 | 21/267 | viral life cycle |
| 7.2E-12 | 14/87 | cellular response to interferon-gamma |
| 0.00013 | 13/255 | regulation of innate immune response |
| 0.0002 | 12/221 | negative regulation of cytokine production |
| 0.0038 | 8/115 | type I interferon production |
| 0.017 | 5/41 | negative regulation of innate immune response |
| 0.017 | 8/143 | interaction with host |
| 0.019 | 45037 | regulation of nuclease activity |
| 0.019 | 7/106 | regulation of type I interferon production |
| 0.019 | 8/149 | regulation of response to cytokine stimulus |
| 0.022 | 5/45 | modulation by symbiont of entry into host |
| 0.022 | 45039 | regulation of response to interferon-gamma |
| 0.022 | 45039 | interferon-gamma-mediated signaling pathway |
| 0.025 | 6/78 | negative regulation of response to biotic stimulus |
| 0.025 | 6/79 | entry into host |
| 0.03 | 5/50 | viral entry into host cell |
| 0.033 | 6/84 | nephron development |
| 0.039 | 9/222 | negative regulation of cell population proliferation |
| 0.046 | 45045 | protein ADP-ribosylation |
| 0.046 | 44996 | regulation of interferon-gamma-mediated signaling pathway |
| 0.056 | 4/31 | adenylyltransferase activity |
| 0.064 | 10/298 | angiogenesis |
| 0.07 | 6/100 | negative regulation of immune response |
| 0.07 | 44998 | ganglion development |
| 0.076 | 6/102 | movement in host environment |
| 0.078 | 6/103 | regulation of epithelial cell differentiation |
| 0.084 | 9/255 | epithelial cell migration |
| 0.084 | 9/256 | negative regulation of locomotion |
| 0.095 | 9/261 | epithelium migration |
| | | ***Input: up-regulated genes*** |
| 0.00022 | 12/292 | skin development |
| 0.00049 | 8/111 | keratinization |
| 0.0054 | 10/283 | epidermal cell differentiation |
| 0.01 | 5/45 | molting cycle |
| 0.035 | 8/220 | response to oxygen levels |
| 0.039 | 11-Mar | regulation of phospholipase A2 activity |
| 0.039 | 8/234 | keratinocyte differentiation |
| 0.039 | 6/111 | regulation of extrinsic apoptotic signaling pathway |

| | | |
|---|---|---|
| 0.059 | 13-Mar | negative regulation of lipase activity |
| 0.059 | 5/79 | entry into host |
| 0.059 | 5/81 | cell-substrate junction assembly |
| 0.059 | 8/270 | regulation of endopeptidase activity |
| 0.059 | 6/129 | negative regulation of peptidase activity |
| 0.066 | 7/208 | response to decreased oxygen levels |
| 0.066 | 4/45 | modulation by symbiont of entry into host |
| 0.066 | 5/87 | cell-substrate junction organization |
| 0.066 | 16-Mar | cell proliferation involved in kidney development |
| 0.069 | 6/146 | extrinsic apoptotic signaling pathway |
| 0.078 | 18-Mar | regulation of water loss via skin |
| 0.078 | 8/299 | regulation of peptidase activity |
| 0.082 | 5/96 | growth factor binding |
| 0.092 | 6/160 | cellular response to oxygen levels |
| 0.092 | 7/232 | regulation of epithelial cell proliferation |
| 0.095 | 5/102 | movement in host environment |
| 0.096 | 5/103 | Notch signaling pathway |

Supplementary Table 4

The Gene Ontology geneset enrichments by GSEA in ΔNp63-depleted cell lines show the enrichments of IFN-I signaling-related pathways (highlighted) upon ΔNp63 depletion. Shown are all genesets (geneset size > 20) significantly (p-value < 0.05) and negatively (normalized enrichment score < 0) associated with depleted p63 expression (p63fKO) as compared control p63 expression.

| *Gene set ID* | *Gene set description* | *Gene set size* | *Enrichment score (scrambled ctrl vs p63fKO)* | *zed enrichment score (scrambled ctrl vs* | *P-value* | *FDR* | *Leading edge size* |
|---|---|---|---|---|---|---|---|
| GOMF_NADPLUS_PROTEIN_ADP_RIBOSYLTRANSFERASE_ACTIVITY | http://www.gsea-msigdb.org/gsea/msigdb/human/geneset/GOMF_NADPLUS_PROTEIN_ADP_RIBOSYLTRANSFERASE_ACTIVITY | 25 | -0.69 | -2.28 | ###### | ###### | 7 |
| GOBP_NEGATIVE_REGULATION_OF_SMOOTH_MUSCLE_CELL_PROLIFERATION | http://www.gsea-msigdb.org/gsea/msigdb/human/geneset/GOBP_NEGATIVE_REGULATION_OF_SMOOTH_MUSCLE_CELL_PROLIFERATION | 44 | -0.53 | -2.12 | ###### | ###### | 10 |
| GOBP_REGULATION_OF_LIPOPOLYSACCHARIDE_MEDIATED_SIGNALING_PATHWAY | http://www.gsea-msigdb.org/gsea/msigdb/human/geneset/GOBP_REGULATION_OF_LIPOPOLYSACCHARIDE_MEDIATED_SIGNALING_PATHWAY | 21 | -0.66 | -2.11 | ###### | ###### | 6 |
| GOBP_REGULATION_OF_TYPE_I_INTERFERON_MEDIATED_SIGNALING_PATHWAY | http://www.gsea-msigdb.org/gsea/msigdb/human/geneset/GOBP_REGULATION_OF_TYPE_I_INTERFERON_MEDIATED_SIGNALING_PATHWAY | 40 | -0.56 | -2.03 | ###### | ###### | 7 |
| GOBP_NEGATIVE_REGULATION_OF_INNATE_IMMUNE_RESPONSE | http://www.gsea-msigdb.org/gsea/msigdb/human/geneset/GOBP_NEGATIVE_REGULATION_OF_INNATE_IMMUNE_RESPONSE | 64 | -0.53 | -2.00 | ###### | ###### | 16 |
| GOBP_RESPONSE_TO_TYPE_I_INTERFERON | http://www.gsea-msigdb.org/gsea/msigdb/human/geneset/GOBP_RESPONSE_TO_TYPE_I_INTERFERON | 68 | -0.61 | -1.95 | ###### | ###### | 16 |

| | | | | | | | |
|---|---|---|---|---|---|---|---|
| GOBP_DETECTION_OF_CHEMICAL_STIMULUS_INVOLVED_IN_SENSORY_PERCEPTION_OF_TASTE | http://www.gsea-msigdb.org/gsea/msigdb/human/geneset/GOBP_DETECTION_OF_CHEMICAL_STIMULUS_INVOLVED_IN_SENSORY_PERCEPTION_OF_TASTE | 25 | -0.69 | -1.92 | ###### | ###### | 9 |
| GOBP_NEGATIVE_REGULATION_OF_RESPONSE_TO_BIOTIC_STIMULUS | http://www.gsea-msigdb.org/gsea/msigdb/human/geneset/GOBP_NEGATIVE_REGULATION_OF_RESPONSE_TO_BIOTIC_STIMULUS | 104 | -0.52 | -1.91 | ###### | ###### | 35 |
| GOBP_INTERFERON_BETA_PRODUCTION | http://www.gsea-msigdb.org/gsea/msigdb/human/geneset/GOBP_INTERFERON_BETA_PRODUCTION | 50 | -0.47 | -1.90 | ###### | ###### | 8 |
| GOBP_DEFENSE_RESPONSE_TO_SYMBIONT | http://www.gsea-msigdb.org/gsea/msigdb/human/geneset/GOBP_DEFENSE_RESPONSE_TO_SYMBIONT | 256 | -0.46 | -1.87 | ###### | ###### | 71 |
| GOBP_TYPE_I_INTERFERON_PRODUCTION | http://www.gsea-msigdb.org/gsea/msigdb/human/geneset/GOBP_TYPE_I_INTERFERON_PRODUCTION | 89 | -0.41 | -1.87 | ###### | ###### | 13 |
| GOBP_POSITIVE_REGULATION_OF_SIGNALING_RECEPTOR_ACTIVITY | http://www.gsea-msigdb.org/gsea/msigdb/human/geneset/GOBP_POSITIVE_REGULATION_OF_SIGNALING_RECEPTOR_ACTIVITY | 37 | -0.53 | -1.85 | ###### | ###### | 3 |
| GOBP_POSITIVE_REGULATION_OF_INTERLEUKIN_10_PRODUCTION | http://www.gsea-msigdb.org/gsea/msigdb/human/geneset/GOBP_POSITIVE_REGULATION_OF_INTERLEUKIN_10_PRODUCTION | 32 | -0.65 | -1.84 | ###### | ###### | 11 |
| GOBP_NEGATIVE_REGULATION_OF_VIRAL_GENOME_REPLICATION | http://www.gsea-msigdb.org/gsea/msigdb/human/geneset/GOBP_NEGATIVE_REGULATION_OF_VIRAL_GENOME_REPLICATION | 51 | -0.62 | -1.81 | ###### | ###### | 21 |
| GOBP_RESPONSE_TO_VIRUS | http://www.gsea-msigdb.org/gsea/msigdb/human/geneset/GOBP_RESPONSE_TO_VIRUS | 340 | -0.39 | -1.78 | ###### | ###### | 55 |

| | | | | | | | |
|---|---|---|---|---|---|---|---|
| GOBP_CHLORIDE_TRANSPORT | http://www.gsea-msigdb.org/gsea/msigdb/human/geneset/GOBP_CHLORIDE_TRANSPORT | 79 | -0.50 | -1.77 | ###### | ###### | 27 |
| GOBP_NEGATIVE_REGULATION_OF_VIRAL_PROCESS | http://www.gsea-msigdb.org/gsea/msigdb/human/geneset/GOBP_NEGATIVE_REGULATION_OF_VIRAL_PROCESS | 80 | -0.58 | -1.77 | ###### | ###### | 38 |
| GOBP_REGULATION_OF_MAST_CELL_ACTIVATION | http://www.gsea-msigdb.org/gsea/msigdb/human/geneset/GOBP_REGULATION_OF_MAST_CELL_ACTIVATION | 31 | -0.49 | -1.76 | ###### | ###### | 6 |

Supplementary Table 5
Summary of the patient details for hNEEO and PDO establishments.

| *Name* | *Established organoid type* | *Patient tissue type* | *Sample acquisition method* | *Prior treatment* | *Stage* | *Culture medium* |
|---|---|---|---|---|---|---|
| 20200819T_K | hNEEO | Tumor tissue | Endoscopic examination | None | n.a. | hNEEO medium |
| EN18n | hNEEO | Adjacent normal tissue | Surgery | None | n.a. | hNEEO medium |
| EN23 | hNEEO | Tumor tissue | Endoscopic examination | None | n.a. | hNEEO medium |
| EN28 | hNEEO | Tumor tissue | Endoscopic examination | None | n.a. | hNEEO medium |
| EN29 | hNEEO | Tumor tissue | Endoscopic examination | None | n.a. | hNEEO medium |
| N200113n | hNEEO | Adjacent normal tissue | Surgery | None | n.a. | hNEEO medium |
| N200120n | hNEEO | Adjacent normal tissue | Surgery | None | n.a. | hNEEO medium |
| N20210203 | hNEEO | Adjacent normal tissue | Surgery | None | n.a. | hNEEO medium |
| ET1c3 | ESCC PDO | Tumor tissue | Surgery | None | pT1bN0 | PDO medium |
| ET2 | ESCC PDO | Tumor tissue | Surgery | None | pT1bN2 | PDO medium |
| ET3 | ESCC PDO | Tumor tissue | Surgery | None | pT2N0 | PDO medium |
| ET5 | ESCC PDO | Tumor tissue | Endoscopic examination | None | Stage III | PDO medium |
| ET6 | ESCC PDO | Tumor tissue | Surgery | None | pT3N2 | PDO medium |
| ET8 | ESCC PDO | Tumor tissue | Endoscopic examination | None | Stage IV | PDO medium |
| ET10P | ESCC PDO | Tumor tissue | Surgery | Neoadjuvant CROSS | pT3, N+ve | PDO medium |
| ET13 | ESCC PDO | Tumor tissue | Endoscopic examination | None | Unknown, N-ve | PDO medium |
| ET15 | ESCC PDO | Tumor tissue | Surgery | Neoadjuvant TPF | pT4N2 | PDO medium |
| ET20 | ESCC PDO | Tumor tissue | Endoscopic examination | None | pTis | PDO medium |
| ET22 | ESCC PDO | Tumor tissue | Surgery | None | pT3N0 | PDO medium |
| ET24 | ESCC PDO | Tumor tissue | Surgery | None | pT4bN0 | PDO medium |
| ET25 | ESCC PDO | Tumor tissue | Endoscopic examination | None | Unknown, N-ve | PDO medium |
| ET26 | ESCC PDO | Tumor tissue | Endoscopic examination | None | Stage III | PDO medium |
| ET27 | ESCC PDO | Tumor tissue | Endoscopic examination | None | Unknown | PDO medium |
| ET29 | ESCC PDO | Tumor tissue | Endoscopic examination | None | Unknown | PDO medium |
| ET32 | ESCC PDO | Tumor tissue | Endoscopic examination | None | Unknown | PDO medium |
| ET4 | EAC PDO | Tumor tissue | Endoscopic examination | None | pT1aN0 | PDO medium |

| | | | | | | |
|---|---|---|---|---|---|---|
| ET14 | EAC PDO | Tumor tissue | Endoscopic examination | None | pT3N2 | PDO medium |
| ET19 | EAC PDO | Tumor tissue | Endoscopic examination | None | pT3N3 | PDO medium |

Supplementary Table 6

The Gene Ontology geneset enrichments by GSEA in three ESCC PDOs with the lowest ΔNp63 expression (ΔNp63lo) versus three PDOs with the highest expression (ΔNp63hi) show the enrichments of IFN-I signaling-related pathways (highlighted) in ΔNp63lo PDOs as highlighted. Shown are all genesets (geneset size > 20) significantly (p-value < 0.05)and negatively (normalized enrichment score < 0) associated with lower p63 expression.

| *Gene set ID* | *Gene set description* | *Gene set size* | *Enrichment score (ΔNp63hi vs ΔNp63lo)* | *Normalized enrichment score (ΔNp63hi vs ΔNp63lo)* | *P-value* | *FDR* | *Leading edge size* |
|---|---|---|---|---|---|---|---|
| GOMF_HISTONE_LYSINE_N_METHYLTRANSFERASE_ACTIVITY | http://www.gsea-msigdb.org/gsea/msigdb/human/geneset/GOMF_HISTONE_LYSINE_N_METHYLTRANSFERASE_ACTIVITY | 48 | -0.48 | -1.95 | 4.00E-02 | 2.25E-01 | 14 |
| GOBP_POSITIVE_REGULATION_OF_RESPONSE_TO_CYTOKINE_STIMULUS | http://www.gsea-msigdb.org/gsea/msigdb/human/geneset/GOBP_POSITIVE_REGULATION_OF_RESPONSE_TO_CYTOKINE_STIMULUS | 47 | -0.53 | -1.91 | 4.26E-02 | 2.06E-01 | 13 |
| GOBP_INTERFERON_MEDIATED_SIGNALING_PATHWAY | http://www.gsea-msigdb.org/gsea/msigdb/human/geneset/GOBP_INTERFERON_MEDIATED_SIGNALING_PATHWAY | 66 | -0.56 | -1.90 | 4.44E-02 | 1.94E-01 | 25 |
| GOMF_LYSINE_N_METHYLTRANSFERASE_ACTIVITY | http://www.gsea-msigdb.org/gsea/msigdb/human/geneset/GOMF_LYSINE_N_METHYLTRANSFERASE_ACTIVITY | 55 | -0.45 | -1.88 | 4.00E-02 | 1.90E-01 | 14 |
| GOBP_POSITIVE_REGULATION_OF_INTERFERON_BETA_PRODUCTION | http://www.gsea-msigdb.org/gsea/msigdb/human/geneset/GOBP_POSITIVE_REGULATION_OF_INTERFERON_BETA_PRODUCTION | 33 | -0.63 | -1.88 | 4.26E-02 | 1.88E-01 | 10 |

| | | | | | | | |
|---|---|---|---|---|---|---|---|
| GOBP_ANTIGEN_PROCESSING_AND_PRESENTATION_OF_PEPTIDE_ANTIGEN | http://www.gsea-msigdb.org/gsea/msigdb/human/geneset/GOBP_ANTIGEN_PROCESSING_AND_PRESENTATION_OF_PEPTIDE_ANTIGEN | 49 | -0.57 | -1.88 | 4.26E-02 | 1.84E-01 | 22 |
| GOBP_RESPONSE_TO_TYPE_I_INTERFERON | http://www.gsea-msigdb.org/gsea/msigdb/human/geneset/GOBP_RESPONSE_TO_TYPE_I_INTERFERON | 58 | -0.53 | -1.85 | 2.13E-02 | 1.86E-01 | 28 |
| GOBP_ANTIGEN_PROCESSING_AND_PRESENTATION | http://www.gsea-msigdb.org/gsea/msigdb/human/geneset/GOBP_ANTIGEN_PROCESSING_AND_PRESENTATION | 74 | -0.48 | -1.84 | 1.96E-02 | 1.85E-01 | 23 |
| GOBP_INTERFERON_BETA_PRODUCTION | http://www.gsea-msigdb.org/gsea/msigdb/human/geneset/GOBP_INTERFERON_BETA_PRODUCTION | 49 | -0.52 | -1.83 | 4.08E-02 | 1.84E-01 | 15 |
| GOBP_POSITIVE_REGULATION_OF_TYROSINE_PHOSPHORYLATION_OF_STAT_PROTEIN | http://www.gsea-msigdb.org/gsea/msigdb/human/geneset/GOBP_POSITIVE_REGULATION_OF_TYROSINE_PHOSPHORYLATION_OF_STAT_PROTEIN | 27 | -0.59 | -1.82 | 4.35E-02 | 1.79E-01 | 9 |
| GOCC_LUMENAL_SIDE_OF_MEMBRANE | http://www.gsea-msigdb.org/gsea/msigdb/human/geneset/GOCC_LUMENAL_SIDE_OF_MEMBRANE | 23 | -0.61 | -1.82 | 4.08E-02 | 1.78E-01 | 10 |

| | | | | | | | |
|---|---|---|---|---|---|---|---|
| GOBP_ANTIGEN_PROCESSING_AND_PRESENTATION_OF_EXOGENOUS_PEPTIDE_ANTIGEN | http://www.gsea-msigdb.org/gsea/msigdb/human/geneset/GOBP_ANTIGEN_PROCESSING_AND_PRESENTATION_OF_EXOGENOUS_PEPTIDE_ANTIGEN | 23 | -0.61 | -1.81 | 2.38E-02 | 1.88E-01 | 10 |
| GOBP_STEROL_BIOSYNTHETIC_PROCESS | http://www.gsea-msigdb.org/gsea/msigdb/human/geneset/GOBP_STEROL_BIOSYNTHETIC_PROCESS | 50 | -0.57 | -1.80 | 3.77E-02 | 1.88E-01 | 20 |
| GOMF_HISTONE_METHYLTRANSFERASE_ACTIVITY | http://www.gsea-msigdb.org/gsea/msigdb/human/geneset/GOMF_HISTONE_METHYLTRANSFERASE_ACTIVITY | 59 | -0.41 | -1.78 | 3.92E-02 | 1.91E-01 | 19 |
| GOBP_IRON_ION_TRANSPORT | http://www.gsea-msigdb.org/gsea/msigdb/human/geneset/GOBP_IRON_ION_TRANSPORT | 43 | -0.53 | -1.78 | 4.26E-02 | 1.94E-01 | 13 |
| GOBP_REGULATION_OF_TYPE_I_INTERFERON_MEDIATED_SIGNALING_PATHWAY | http://www.gsea-msigdb.org/gsea/msigdb/human/geneset/GOBP_REGULATION_OF_TYPE_I_INTERFERON_MEDIATED_SIGNALING_PATHWAY | 37 | -0.56 | -1.78 | 2.27E-02 | 1.94E-01 | 18 |
| GOMF_RNA_POLYMERASE_CORE_ENZYME_BINDING | http://www.gsea-msigdb.org/gsea/msigdb/human/geneset/GOMF_RNA_POLYMERASE_CORE_ENZYME_BINDING | 39 | -0.51 | -1.78 | 0.00E+00 | 1.94E-01 | 4 |
| GOBP_DEFENSE_RESPONSE_TO_SYMBIONT | http://www.gsea-msigdb.org/gsea/msigdb/human/geneset/GOBP_DEFENSE_RESPONSE_TO_SYMBIONT | 211 | -0.44 | -1.77 | 2.08E-02 | 1.91E-01 | 83 |

| | | | | | | | |
|---|---|---|---|---|---|---|---|
| GOBP_ACUTE_INFLAMMATORY_RESPONSE | http://www.gsea-msigdb.org/gsea/msigdb/human/geneset/GOBP_ACUTE_INFLAMMATORY_RESPONSE | 45 | -0.52 | -1.77 | 4.17E-02 | 1.90E-01 | 17 |
| GOMF_PHOSPHATASE_INHIBITOR_ACTIVITY | http://www.gsea-msigdb.org/gsea/msigdb/human/geneset/GOMF_PHOSPHATASE_INHIBITOR_ACTIVITY | 28 | -0.50 | -1.76 | 4.26E-02 | 2.03E-01 | 11 |
| GOBP_T_CELL_MEDIATED_IMMUNITY | http://www.gsea-msigdb.org/gsea/msigdb/human/geneset/GOBP_T_CELL_MEDIATED_IMMUNITY | 69 | -0.48 | -1.75 | 2.13E-02 | 1.98E-01 | 28 |
| GOBP_ANTIGEN_PROCESSING_AND_PRESENTATION_OF_PEPTIDE_ANTIGEN_VIA_MHC_CLASS_I | http://www.gsea-msigdb.org/gsea/msigdb/human/geneset/GOBP_ANTIGEN_PROCESSING_AND_PRESENTATION_OF_PEPTIDE_ANTIGEN_VIA_MHC_CLASS_I | 31 | -0.66 | -1.75 | 2.04E-02 | 1.96E-01 | 17 |
| GOBP_ENDOPLASMIC_RETICULUM_TO_CYTOSOL_TRANSPORT | http://www.gsea-msigdb.org/gsea/msigdb/human/geneset/GOBP_ENDOPLASMIC_RETICULUM_TO_CYTOSOL_TRANSPORT | 26 | -0.57 | -1.75 | 0.00E+00 | 1.97E-01 | 14 |
| GOBP_MIRNA_PROCESSING | http://www.gsea-msigdb.org/gsea/msigdb/human/geneset/GOBP_MIRNA_PROCESSING | 40 | -0.51 | -1.74 | 0.00E+00 | 2.01E-01 | 8 |
| GOBP_PEPTIDYL_LYSINE_DIMETHYLATION | http://www.gsea-msigdb.org/gsea/msigdb/human/geneset/GOBP_PEPTIDYL_LYSINE_DIMETHYLATION | 22 | -0.55 | -1.74 | 4.65E-02 | 2.01E-01 | 6 |

| | | | | | | | |
|---|---|---|---|---|---|---|---|
| GOBP_GRANULOCYTE_DIFFERENTIATION | http://www.gsea-msigdb.org/gsea/msigdb/human/geneset/GOBP_GRANULOCYTE_DIFFERENTIATION | 23 | -0.49 | -1.74 | 4.55E-02 | 1.99E-01 | 7 |
| GOBP_ANOIKIS | http://www.gsea-msigdb.org/gsea/msigdb/human/geneset/GOBP_ANOIKIS | 31 | -0.54 | -1.72 | 0.00E+00 | 2.23E-01 | 9 |
| GOBP_ER_NUCLEUS_SIGNALING_PATHWAY | http://www.gsea-msigdb.org/gsea/msigdb/human/geneset/GOBP_ER_NUCLEUS_SIGNALING_PATHWAY | 46 | -0.49 | -1.71 | 2.13E-02 | 2.27E-01 | 25 |
| GOBP_LYSOSOMAL_LUMEN_ACIDIFICATION | http://www.gsea-msigdb.org/gsea/msigdb/human/geneset/GOBP_LYSOSOMAL_LUMEN_ACIDIFICATION | 21 | -0.55 | -1.71 | 1.79E-02 | 2.28E-01 | 8 |
| GOMF_PEPTIDASE_ACTIVATOR_ACTIVITY | http://www.gsea-msigdb.org/gsea/msigdb/human/geneset/GOMF_PEPTIDASE_ACTIVATOR_ACTIVITY | 44 | -0.45 | -1.71 | 4.35E-02 | 2.31E-01 | 14 |
| GOCC_SPECIFIC_GRANULE_LUMEN | http://www.gsea-msigdb.org/gsea/msigdb/human/geneset/GOCC_SPECIFIC_GRANULE_LUMEN | 36 | -0.52 | -1.70 | 0.00E+00 | 2.35E-01 | 16 |
| GOBP_POSITIVE_REGULATION_OF_TYPE_I_INTERFERON_PRODUCTION | http://www.gsea-msigdb.org/gsea/msigdb/human/geneset/GOBP_POSITIVE_REGULATION_OF_TYPE_I_INTERFERON_PRODUCTION | 52 | -0.53 | -1.70 | 4.08E-02 | 2.34E-01 | 13 |
| GOBP_NEGATIVE_REGULATION_OF_HEMOPOIESIS | http://www.gsea-msigdb.org/gsea/msigdb/human/geneset/GOBP_NEGATIVE_REGULATION_OF_HEMOPOIESIS | 61 | -0.38 | -1.70 | 0.00E+00 | 2.32E-01 | 14 |

| | | | | | | | |
|---|---|---|---|---|---|---|---|
| GOBP_TRANSITION_METAL_ION_TRANSPORT | http://www.gsea-msigdb.org/gsea/msigdb/human/geneset/GOBP_TRANSITION_METAL_ION_TRANSPORT | 72 | -0.44 | -1.69 | 2.00E-02 | 2.33E-01 | 13 |
| GOBP_POSITIVE_REGULATION_OF_INTERLEUKIN_12_PRODUCTION | http://www.gsea-msigdb.org/gsea/msigdb/human/geneset/GOBP_POSITIVE_REGULATION_OF_INTERLEUKIN_12_PRODUCTION | 21 | -0.51 | -1.69 | 3.77E-02 | 2.38E-01 | 5 |
| GOBP_TYROSINE_PHOSPHORYLATION_OF_STAT_PROTEIN | http://www.gsea-msigdb.org/gsea/msigdb/human/geneset/GOBP_TYROSINE_PHOSPHORYLATION_OF_STAT_PROTEIN | 41 | -0.47 | -1.69 | 0.00E+00 | 2.38E-01 | 10 |
| GOMF_N_METHYLTRANSFERASE_ACTIVITY | http://www.gsea-msigdb.org/gsea/msigdb/human/geneset/GOMF_N_METHYLTRANSFERASE_ACTIVITY | 86 | -0.37 | -1.68 | 2.17E-02 | 2.40E-01 | 15 |
| GOBP_CYTOKINE_MEDIATED_SIGNALING_PATHWAY | http://www.gsea-msigdb.org/gsea/msigdb/human/geneset/GOBP_CYTOKINE_MEDIATED_SIGNALING_PATHWAY | 277 | -0.37 | -1.68 | 0.00E+00 | 2.44E-01 | 69 |
| GOBP_NEGATIVE_REGULATION_OF_LYMPHOCYTE_ACTIVATION | http://www.gsea-msigdb.org/gsea/msigdb/human/geneset/GOBP_NEGATIVE_REGULATION_OF_LYMPHOCYTE_ACTIVATION | 85 | -0.36 | -1.68 | 2.17E-02 | 2.44E-01 | 24 |
| GOBP_RESPONSE_TO_VIRUS | http://www.gsea-msigdb.org/gsea/msigdb/human/geneset/GOBP_RESPONSE_TO_VIRUS | 285 | -0.40 | -1.67 | 4.17E-02 | 2.47E-01 | 119 |

| | | | | | | | |
|---|---|---|---|---|---|---|---|
| GOBP_PROTEIN_EXIT_FROM_ENDOPLASMIC_RETICULUM | http://www.gsea-msigdb.org/gsea/msigdb/human/geneset/GOBP_PROTEIN_EXIT_FROM_ENDOPLASMIC_RETICULUM | 43 | -0.43 | -1.67 | 4.35E-02 | 2.48E-01 | 18 |
| GOBP_ADAPTIVE_IMMUNE_RESPONSE | http://www.gsea-msigdb.org/gsea/msigdb/human/geneset/GOBP_ADAPTIVE_IMMUNE_RESPONSE | 230 | -0.36 | -1.67 | 2.13E-02 | 2.47E-01 | 68 |
| GOBP_IRON_ION_HOMEOSTASIS | http://www.gsea-msigdb.org/gsea/msigdb/human/geneset/GOBP_IRON_ION_HOMEOSTASIS | 72 | -0.44 | -1.67 | 4.08E-02 | 2.47E-01 | 29 |
| GOBP_ISOPRENOID_BIOSYNTHETIC_PROCESS | http://www.gsea-msigdb.org/gsea/msigdb/human/geneset/GOBP_ISOPRENOID_BIOSYNTHETIC_PROCESS | 24 | -0.59 | -1.67 | 3.77E-02 | 2.50E-01 | 11 |

Supplementary Table 7

The transcription factor target geneset enrichments by GSEA in ΔNp63-depleted cell lines show the enrichment of genes with STAT1 binding site (bolded) and with IRF1 binding site (highlighted) upon ΔNp63 depletion. Shown are all genesets (geneset size > 20) significantly (p-value < 0.05) and negatively (normalized enrichment score < 0) associated with depleted p63 expression (p63fKO) as compared control p63 expression.

| *Gene set ID* | *Gene set description* | *Gene set size* | *Enrichment score (scrambled ctrl vs p63fKO)* | *Normalized enrichment score (scrambled ctrl vs p63fKO)* | *P-value* | *FDR* | *Leading edge size* |
|---|---|---|---|---|---|---|---|
| FXR_Q3 | http://www.gsea-msigdb.org/gsea/msigdb/human/geneset/FXR_Q3 | 93 | -0.46 | -1.98 | 3.39E-02 | 2.10E-02 | 15 |
| FOXD3_TARGET_GENES | http://www.gsea-msigdb.org/gsea/msigdb/human/geneset/FOXD3_TARGET_GENES | 32 | -0.55 | -1.86 | 3.92E-02 | 7.33E-02 | 9 |
| PAX8_01 | http://www.gsea-msigdb.org/gsea/msigdb/human/geneset/PAX8_01 | 37 | -0.44 | -1.74 | 0.00E+00 | 1.34E-01 | 7 |
| **ISRE_01** | http://www.gsea-msigdb.org/gsea/msigdb/human/geneset/ISRE_01 | 215 | -0.39 | -1.71 | 1.67E-02 | 1.39E-01 | 51 |
| STTTCRNTTT_IRF_Q6 | http://www.gsea-msigdb.org/gsea/msigdb/human/geneset/STTTCRNTTT_IRF_Q6 | 173 | -0.40 | -1.63 | 1.72E-02 | 2.10E-01 | 43 |
| ICSBP_Q6 | http://www.gsea-msigdb.org/gsea/msigdb/human/geneset/ICSBP_Q6 | 223 | -0.31 | -1.59 | 3.33E-02 | 2.48E-01 | 34 |
| IRF_Q6 | http://www.gsea-msigdb.org/gsea/msigdb/human/geneset/IRF_Q6 | 211 | -0.38 | -1.52 | 1.69E-02 | 3.22E-01 | 55 |
| METTL14_TARGET_GENES | http://www.gsea-msigdb.org/gsea/msigdb/human/geneset/METTL14_TARGET_GENES | 120 | -0.33 | -1.52 | 4.08E-02 | 3.02E-01 | 36 |
| IRF1_01 | http://www.gsea-msigdb.org/gsea/msigdb/human/geneset/IRF1_01 | 214 | -0.33 | -1.50 | 1.82E-02 | 3.06E-01 | 48 |
| WYAAANNRNNNGCG_UNKNOWN | http://www.gsea-msigdb.org/gsea/msigdb/human/geneset/WYAAANNRNNNGCG_UNKNOWN | 54 | -0.34 | -1.47 | 4.65E-02 | 3.30E-01 | 5 |
| ZNF704_TARGET_GENES | http://www.gsea-msigdb.org/gsea/msigdb/human/geneset/ZNF704_TARGET_GENES | 68 | -0.37 | -1.43 | 4.08E-02 | 4.05E-01 | 10 |

| | | | | | | | |
|---|---|---|---|---|---|---|---|
| S8_01 | http://www.gsea-msigdb.org/gsea/msigdb/human/geneset/S8_01 | 195 | -0.32 | -1.42 | 1.89E-02 | 3.87E-01 | 40 |
| GATA4_Q3 | http://www.gsea-msigdb.org/gsea/msigdb/human/geneset/GATA4_Q3 | 216 | -0.26 | -1.42 | 3.92E-02 | 3.81E-01 | 48 |
| AREB6_02 | http://www.gsea-msigdb.org/gsea/msigdb/human/geneset/AREB6_02 | 228 | -0.29 | -1.41 | 1.79E-02 | 3.68E-01 | 41 |
| TGIF_01 | http://www.gsea-msigdb.org/gsea/msigdb/human/geneset/TGIF_01 | 224 | -0.29 | -1.40 | 1.85E-02 | 3.68E-01 | 50 |
| HSF1_01 | http://www.gsea-msigdb.org/gsea/msigdb/human/geneset/HSF1_01 | 235 | -0.26 | -1.40 | 1.75E-02 | 3.54E-01 | 33 |
| TTAYRTAA_E4BP4_01 | http://www.gsea-msigdb.org/gsea/msigdb/human/geneset/TTAYRTAA_E4BP4_01 | 236 | -0.26 | -1.39 | 2.04E-02 | 3.60E-01 | 42 |
| TAAWWATAG_RSRFC4_Q2 | http://www.gsea-msigdb.org/gsea/msigdb/human/geneset/TAAWWATAG_RSRFC4_Q2 | 146 | -0.29 | -1.37 | 2.00E-02 | 4.00E-01 | 34 |
| TCF11MAFG_01 | http://www.gsea-msigdb.org/gsea/msigdb/human/geneset/TCF11MAFG_01 | 182 | -0.28 | -1.35 | 5.00E-02 | 4.28E-01 | 27 |
| GATA6_01 | http://www.gsea-msigdb.org/gsea/msigdb/human/geneset/GATA6_01 | 214 | -0.26 | -1.34 | 2.17E-02 | 4.35E-01 | 39 |
| E4BP4_01 | http://www.gsea-msigdb.org/gsea/msigdb/human/geneset/E4BP4_01 | 198 | -0.25 | -1.29 | 5.00E-02 | 4.92E-01 | 27 |
| OCT1_04 | http://www.gsea-msigdb.org/gsea/msigdb/human/geneset/OCT1_04 | 198 | -0.28 | -1.28 | 3.92E-02 | 4.91E-01 | 50 |
| MEF2_Q6_01 | http://www.gsea-msigdb.org/gsea/msigdb/human/geneset/MEF2_Q6_01 | 200 | -0.31 | -1.28 | 2.08E-02 | 4.84E-01 | 47 |
| IRF7_01 | http://www.gsea-msigdb.org/gsea/msigdb/human/geneset/IRF7_01 | 227 | -0.26 | -1.27 | 4.08E-02 | 4.79E-01 | 42 |
| RSRFC4_Q2 | http://www.gsea-msigdb.org/gsea/msigdb/human/geneset/RSRFC4_Q2 | 177 | -0.26 | -1.24 | 2.22E-02 | 4.75E-01 | 37 |

| | | | | | | | |
|---|---|---|---|---|---|---|---|
| HNF6_Q6 | http://www.gsea-msigdb.org/gsea/msigdb/human/geneset/HNF6_Q6 | 195 | -0.25 | -1.23 | 3.51E-02 | 4.59E-01 | 30 |
| STAT_01 | http://www.gsea-msigdb.org/gsea/msigdb/human/geneset/STAT_01 | 217 | -0.26 | -1.22 | 4.44E-02 | 4.50E-01 | 32 |

Supplementary Table 8

The transcription factor target geneset enrichments by GSEA in three ESCC PDOs with the lowest ΔNp63 expression (ΔNp63lo) versus three PDOs with the highest expression (ΔNp63hi) show the enrichment of genes with STAT1 binding site (bolded) and with IRF1 binding site (highlighted) upon ΔNp63 depletion. Shown are all genesets (geneset size > 20) significantly (p-value < 0.05) and negatively (normalized enrichment score < 0) associated with depleted p63 expression (p63fKO) as compared control p63 expression.

| *Gene set ID* | *Gene set description* | *Gene set size* | *Enrichment score (ΔNp63hi vs ΔNp63lo)* | *Normalized enrichment score (ΔNp63hi vs ΔNp63lo)* | *P-value* | *FDR* | *Leading edge size* |
|---|---|---|---|---|---|---|---|
| MCRS1_TARGET_GENES | http://www.gsea-msigdb.org/gsea/msigdb/human/geneset/MCRS1_TARGET_GENES | 34 | -0.56 | -1.85 | 1.72E-02 | 9.39E-02 | 9 |
| STTTCRNTTT_IRF_Q6 | http://www.gsea-msigdb.org/gsea/msigdb/human/geneset/STTTCRNTTT_IRF_Q6 | 142 | -0.42 | -1.78 | 4.88E-02 | 1.14E-01 | 45 |
| CCANNAGRKGGC_UNKNOWN | http://www.gsea-msigdb.org/gsea/msigdb/human/geneset/CCANNAGRKGGC_UNKNOWN | 72 | -0.46 | -1.68 | 4.55E-02 | 3.89E-01 | 12 |
| ZNF165_TARGET_GENES | http://www.gsea-msigdb.org/gsea/msigdb/human/geneset/ZNF165_TARGET_GENES | 67 | -0.40 | -1.68 | 0.00E+00 | 2.92E-01 | 22 |
| IRF_Q6 | http://www.gsea-msigdb.org/gsea/msigdb/human/geneset/IRF_Q6 | 164 | -0.41 | -1.66 | 2.08E-02 | 2.71E-01 | 55 |
| ZNF26_TARGET_GENES | http://www.gsea-msigdb.org/gsea/msigdb/human/geneset/ZNF26_TARGET_GENES | 21 | -0.53 | -1.65 | 3.39E-02 | 2.44E-01 | 7 |
| NFKAPPAB_01 | http://www.gsea-msigdb.org/gsea/msigdb/human/geneset/NFKAPPAB_01 | 177 | -0.34 | -1.60 | 3.92E-02 | 3.30E-01 | 41 |
| KCCGNSWTTT_UNKNOWN | http://www.gsea-msigdb.org/gsea/msigdb/human/geneset/KCCGNSWTTT_UNKNOWN | 97 | -0.38 | -1.60 | 3.77E-02 | 3.30E-01 | 23 |
| AACWWCAANK_UNKNOWN | http://www.gsea-msigdb.org/gsea/msigdb/human/geneset/AACWWCAANK_UNKNOWN | 120 | -0.40 | -1.60 | 3.77E-02 | 3.10E-01 | 37 |
| **ISRE_01** | http://www.gsea-msigdb.org/gsea/msigdb/human/geneset/ISRE_01 | 177 | -0.37 | -1.57 | 2.13E-02 | 3.35E-01 | 51 |
| IRF2_01 | http://www.gsea-msigdb.org/gsea/msigdb/human/geneset/IRF2_01 | 91 | -0.35 | -1.57 | 0.00E+00 | 3.22E-01 | 25 |

| | | | | | | | |
|---|---|---|---|---|---|---|---|
| NFKAPPAB65_01 | http://www.gsea-msigdb.org/gsea/msigdb/human/geneset/NFKAPPAB65_01 | 169 | -0.35 | -1.56 | 4.35E-02 | 3.06E-01 | 43 |
| PTTG1_TARGET_GENES | http://www.gsea-msigdb.org/gsea/msigdb/human/geneset/PTTG1_TARGET_GENES | 56 | -0.40 | -1.54 | 0.00E+00 | 3.02E-01 | 15 |
| ZNF354B_TARGET_GENES | http://www.gsea-msigdb.org/gsea/msigdb/human/geneset/ZNF354B_TARGET_GENES | 172 | -0.41 | -1.54 | 3.57E-02 | 2.85E-01 | 50 |
| LMX1B_TARGET_GENES | http://www.gsea-msigdb.org/gsea/msigdb/human/geneset/LMX1B_TARGET_GENES | 21 | -0.54 | -1.50 | 0.00E+00 | 4.49E-01 | 11 |
| FOXC1_TARGET_GENES | http://www.gsea-msigdb.org/gsea/msigdb/human/geneset/FOXC1_TARGET_GENES | 55 | -0.40 | -1.49 | 2.04E-02 | 4.81E-01 | 17 |
| TAATTA_CHX10_01 | http://www.gsea-msigdb.org/gsea/msigdb/human/geneset/TAATTA_CHX10_01 | 499 | -0.28 | -1.48 | 4.17E-02 | 4.84E-01 | 113 |
| ZNF746_TARGET_GENES | http://www.gsea-msigdb.org/gsea/msigdb/human/geneset/ZNF746_TARGET_GENES | 59 | -0.37 | -1.47 | 4.35E-02 | 4.79E-01 | 14 |
| MSX1_TARGET_GENES | http://www.gsea-msigdb.org/gsea/msigdb/human/geneset/MSX1_TARGET_GENES | 167 | -0.33 | -1.47 | 2.04E-02 | 4.70E-01 | 62 |
| ZNF532_TARGET_GENES | http://www.gsea-msigdb.org/gsea/msigdb/human/geneset/ZNF532_TARGET_GENES | 54 | -0.37 | -1.46 | 3.51E-02 | 5.03E-01 | 18 |
| LBP1_Q6 | http://www.gsea-msigdb.org/gsea/msigdb/human/geneset/LBP1_Q6 | 142 | -0.36 | -1.46 | 4.00E-02 | 4.85E-01 | 41 |
| ARID3B_TARGET_GENES | http://www.gsea-msigdb.org/gsea/msigdb/human/geneset/ARID3B_TARGET_GENES | 102 | -0.37 | -1.45 | 2.27E-02 | 4.80E-01 | 25 |
| IRF1_Q6 | http://www.gsea-msigdb.org/gsea/msigdb/human/geneset/IRF1_Q6 | 168 | -0.34 | -1.45 | 2.33E-02 | 4.87E-01 | 43 |
| P300_01 | http://www.gsea-msigdb.org/gsea/msigdb/human/geneset/P300_01 | 198 | -0.39 | -1.45 | 4.65E-02 | 4.71E-01 | 52 |
| CRGAARNNNNCGA_UNKNOWN | http://www.gsea-msigdb.org/gsea/msigdb/human/geneset/CRGAARNNNNCGA_UNKNOWN | 44 | -0.46 | -1.44 | 0.00E+00 | 4.81E-01 | 16 |

| | | | | | | | |
|---|---|---|---|---|---|---|---|
| CHAF1A_TARGET_GENES | http://www.gsea-msigdb.org/gsea/msigdb/human/geneset/CHAF1A_TARGET_GENES | 79 | -0.40 | -1.43 | 4.26E-02 | 5.21E-01 | 23 |
| CREL_01 | http://www.gsea-msigdb.org/gsea/msigdb/human/geneset/CREL_01 | 180 | -0.31 | -1.43 | 0.00E+00 | 5.02E-01 | 42 |
| TFCP2_TARGET_GENES | http://www.gsea-msigdb.org/gsea/msigdb/human/geneset/TFCP2_TARGET_GENES | 83 | -0.40 | -1.42 | 2.08E-02 | 5.13E-01 | 22 |
| NFKB_Q6 | http://www.gsea-msigdb.org/gsea/msigdb/human/geneset/NFKB_Q6 | 189 | -0.32 | -1.42 | 3.92E-02 | 5.02E-01 | 45 |
| MAZ_Q6 | http://www.gsea-msigdb.org/gsea/msigdb/human/geneset/MAZ_Q6 | 151 | -0.36 | -1.42 | 0.00E+00 | 4.91E-01 | 53 |
| ICSBP_Q6 | http://www.gsea-msigdb.org/gsea/msigdb/human/geneset/ICSBP_Q6 | 183 | -0.33 | -1.41 | 2.27E-02 | 5.29E-01 | 65 |
| GABP_B | http://www.gsea-msigdb.org/gsea/msigdb/human/geneset/GABP_B | 251 | -0.32 | -1.41 | 4.17E-02 | 5.44E-01 | 76 |
| MED16_TARGET_GENES | http://www.gsea-msigdb.org/gsea/msigdb/human/geneset/MED16_TARGET_GENES | 23 | -0.46 | -1.41 | 4.26E-02 | 5.29E-01 | 14 |
| CCTNTMAGA_UNKNOWN | http://www.gsea-msigdb.org/gsea/msigdb/human/geneset/CCTNTMAGA_UNKNOWN | 97 | -0.32 | -1.40 | 4.44E-02 | 5.22E-01 | 29 |
| LEF1_Q2 | http://www.gsea-msigdb.org/gsea/msigdb/human/geneset/LEF1_Q2 | 155 | -0.32 | -1.40 | 4.08E-02 | 5.22E-01 | 45 |
| RYCACNNRNNRNCAG_UNKNOWN | http://www.gsea-msigdb.org/gsea/msigdb/human/geneset/RYCACNNRNNRNCAG_UNKNOWN | 55 | -0.39 | -1.37 | 4.55E-02 | 6.26E-01 | 11 |
| IK1_01 | http://www.gsea-msigdb.org/gsea/msigdb/human/geneset/IK1_01 | 198 | -0.28 | -1.37 | 4.65E-02 | 6.36E-01 | 59 |
| GATA6_01 | http://www.gsea-msigdb.org/gsea/msigdb/human/geneset/GATA6_01 | 153 | -0.29 | -1.36 | 4.44E-02 | 6.27E-01 | 31 |
| HOXC13_TARGET_GENES | http://www.gsea-msigdb.org/gsea/msigdb/human/geneset/HOXC13_TARGET_GENES | 104 | -0.35 | -1.36 | 4.35E-02 | 6.36E-01 | 24 |
| PAX4_04 | http://www.gsea-msigdb.org/gsea/msigdb/human/geneset/PAX4_04 | 153 | -0.30 | -1.36 | 2.44E-02 | 6.38E-01 | 46 |
| NCOA2_TARGET_GENES | http://www.gsea-msigdb.org/gsea/msigdb/human/geneset/NCOA2_TARGET_GENES | 374 | -0.28 | -1.33 | 4.17E-02 | 6.91E-01 | 116 |

| | | | | | | | |
|---|---|---|---|---|---|---|---|
| PAX4_02 | http://www.gsea-msigdb.org/gsea/msigdb/human/geneset/PAX4_02 | 143 | -0.31 | -1.33 | 2.44E-02 | 6.83E-01 | 43 |
| STAT5A_04 | http://www.gsea-msigdb.org/gsea/msigdb/human/geneset/STAT5A_04 | 136 | -0.29 | -1.32 | 0.00E+00 | 7.29E-01 | 35 |
| IRF7_01 | http://www.gsea-msigdb.org/gsea/msigdb/human/geneset/IRF7_01 | 175 | -0.31 | -1.32 | 0.00E+00 | 7.19E-01 | 47 |
| ZNF707_TARGET_GENES | http://www.gsea-msigdb.org/gsea/msigdb/human/geneset/ZNF707_TARGET_GENES | 27 | -0.36 | -1.31 | 0.00E+00 | 7.14E-01 | 7 |
| TERF1_TARGET_GENES | http://www.gsea-msigdb.org/gsea/msigdb/human/geneset/TERF1_TARGET_GENES | 185 | -0.28 | -1.31 | 2.27E-02 | 7.03E-01 | 47 |
| P53_02 | http://www.gsea-msigdb.org/gsea/msigdb/human/geneset/P53_02 | 179 | -0.34 | -1.31 | 5.00E-02 | 7.11E-01 | 29 |
| TAAYNRNNTCC_UNKNOWN | http://www.gsea-msigdb.org/gsea/msigdb/human/geneset/TAAYNRNNTCC_UNKNOWN | 107 | -0.35 | -1.31 | 0.00E+00 | 7.00E-01 | 34 |
| DLX2_TARGET_GENES | http://www.gsea-msigdb.org/gsea/msigdb/human/geneset/DLX2_TARGET_GENES | 204 | -0.31 | -1.30 | 4.17E-02 | 7.16E-01 | 69 |
| AREB6_04 | http://www.gsea-msigdb.org/gsea/msigdb/human/geneset/AREB6_04 | 173 | -0.30 | -1.30 | 2.44E-02 | 7.01E-01 | 31 |
| MEIS1_01 | http://www.gsea-msigdb.org/gsea/msigdb/human/geneset/MEIS1_01 | 165 | -0.27 | -1.30 | 2.13E-02 | 6.98E-01 | 28 |
| RFX1_01 | http://www.gsea-msigdb.org/gsea/msigdb/human/geneset/RFX1_01 | 188 | -0.26 | -1.29 | 4.35E-02 | 7.12E-01 | 53 |
| CEBPB_02 | http://www.gsea-msigdb.org/gsea/msigdb/human/geneset/CEBPB_02 | 191 | -0.27 | -1.29 | 2.17E-02 | 7.22E-01 | 35 |
| SOX10_TARGET_GENES | http://www.gsea-msigdb.org/gsea/msigdb/human/geneset/SOX10_TARGET_GENES | 180 | -0.26 | -1.28 | 2.17E-02 | 7.33E-01 | 44 |
| MYOD_Q6 | http://www.gsea-msigdb.org/gsea/msigdb/human/geneset/MYOD_Q6 | 161 | -0.30 | -1.27 | 1.89E-02 | 7.63E-01 | 36 |
| ZNF16_TARGET_GENES | http://www.gsea-msigdb.org/gsea/msigdb/human/geneset/ZNF16_TARGET_GENES | 53 | -0.41 | -1.27 | 4.65E-02 | 7.59E-01 | 17 |
| RTTTNNNYTGGM_UNKNOWN | http://www.gsea-msigdb.org/gsea/msigdb/human/geneset/RTTTNNNYTGGM_UNKNOWN | 99 | -0.31 | -1.27 | 0.00E+00 | 7.54E-01 | 32 |

| | | | | | | | |
|---|---|---|---|---|---|---|---|
| MEIS1BHOXA9_01 | http://www.gsea-msigdb.org/gsea/msigdb/human/geneset/MEIS1BHOXA9_01 | 83 | -0.30 | -1.26 | 4.55E-02 | 7.55E-01 | 13 |
| TOP2B_TARGET_GENES | http://www.gsea-msigdb.org/gsea/msigdb/human/geneset/TOP2B_TARGET_GENES | 499 | -0.28 | -1.26 | 2.17E-02 | 7.58E-01 | 158 |
| NR1D1_TARGET_GENES | http://www.gsea-msigdb.org/gsea/msigdb/human/geneset/NR1D1_TARGET_GENES | 105 | -0.28 | -1.25 | 4.88E-02 | 7.60E-01 | 15 |
| STAT5A_03 | http://www.gsea-msigdb.org/gsea/msigdb/human/geneset/STAT5A_03 | 182 | -0.28 | -1.25 | 2.17E-02 | 7.57E-01 | 32 |
| CREBP1_01 | http://www.gsea-msigdb.org/gsea/msigdb/human/geneset/CREBP1_01 | 142 | -0.27 | -1.24 | 2.63E-02 | 7.70E-01 | 30 |
| TTAYRTAA_E4BP4_01 | http://www.gsea-msigdb.org/gsea/msigdb/human/geneset/TTAYRTAA_E4BP4_01 | 201 | -0.27 | -1.23 | 2.38E-02 | 8.04E-01 | 45 |
| HMGB2_TARGET_GENES | http://www.gsea-msigdb.org/gsea/msigdb/human/geneset/HMGB2_TARGET_GENES | 406 | -0.27 | -1.22 | 2.17E-02 | 8.01E-01 | 86 |
| NKX2_8_TARGET_GENES | http://www.gsea-msigdb.org/gsea/msigdb/human/geneset/NKX2_8_TARGET_GENES | 29 | -0.36 | -1.21 | 0.00E+00 | 8.06E-01 | 13 |
| NKX25_01 | http://www.gsea-msigdb.org/gsea/msigdb/human/geneset/NKX25_01 | 71 | -0.25 | -1.20 | 4.35E-02 | 8.02E-01 | 19 |
| EFC_Q6 | http://www.gsea-msigdb.org/gsea/msigdb/human/geneset/EFC_Q6 | 183 | -0.27 | -1.19 | 4.88E-02 | 8.18E-01 | 59 |
| TFIIA_Q6 | http://www.gsea-msigdb.org/gsea/msigdb/human/geneset/TFIIA_Q6 | 189 | -0.25 | -1.18 | 4.26E-02 | 8.19E-01 | 50 |
| IRF1_01 | http://www.gsea-msigdb.org/gsea/msigdb/human/geneset/IRF1_01 | 168 | -0.25 | -1.18 | 0.00E+00 | 8.19E-01 | 40 |
| HOXA10_TARGET_GENES | http://www.gsea-msigdb.org/gsea/msigdb/human/geneset/HOXA10_TARGET_GENES | 229 | -0.28 | -1.18 | 0.00E+00 | 8.18E-01 | 46 |
| STAT4_01 | http://www.gsea-msigdb.org/gsea/msigdb/human/geneset/STAT4_01 | 172 | -0.24 | -1.16 | 0.00E+00 | 8.44E-01 | 51 |
| ZNF197_TARGET_GENES | http://www.gsea-msigdb.org/gsea/msigdb/human/geneset/ZNF197_TARGET_GENES | 483 | -0.23 | -1.15 | 4.26E-02 | 8.35E-01 | 73 |

| | | | | | | | |
|---|---|---|---|---|---|---|---|
| YGCANTGCR_UNKNOWN | http://www.gsea-msigdb.org/gsea/msigdb/human/geneset/YGCANTGCR_UNKNOWN | 100 | -0.26 | -1.15 | 4.88E-02 | 8.68E-01 | 21 |
| CEBPA_01 | http://www.gsea-msigdb.org/gsea/msigdb/human/geneset/CEBPA_01 | 183 | -0.24 | -1.15 | 0.00E+00 | 8.63E-01 | 41 |
| CEBP_01 | http://www.gsea-msigdb.org/gsea/msigdb/human/geneset/CEBP_01 | 173 | -0.25 | -1.14 | 4.00E-02 | 8.59E-01 | 29 |
| ALPHACP1_01 | http://www.gsea-msigdb.org/gsea/msigdb/human/geneset/ALPHACP1_01 | 189 | -0.23 | -1.12 | 4.17E-02 | 9.00E-01 | 53 |
| CDPCR1_01 | http://www.gsea-msigdb.org/gsea/msigdb/human/geneset/CDPCR1_01 | 78 | -0.25 | -1.10 | 2.17E-02 | 9.04E-01 | 17 |
| NKX3A_01 | http://www.gsea-msigdb.org/gsea/msigdb/human/geneset/NKX3A_01 | 136 | -0.24 | -1.10 | 4.17E-02 | 9.01E-01 | 31 |

Supplementary Table 9

The Gene Ontology geneset enrichments by GSEA in three hNEEOs with the lowest ΔNp63 expression (ΔNp63lo) versus three hNEEOs with the highest expression (ΔNp63hi). Shown are all genesets related to IFN-I signaling that show significant enrichments in ΔNp63-depleted ESCC cell line and ESCC PDO analyses.

| *Gene set ID* | *Gene set description* | *Gene set size* | *Enrichment score (scrambled ctrl vs p63fKO)* | *Normalized enrichment score (scrambled ctrl vs p63fKO)* | *P-value* | *FDR* | *Leading edge size* |
|---|---|---|---|---|---|---|---|
| GOBP_ANTIGEN_PROCESSING_AND_PRESENTATION_OF_ENDOGENOUS_PEPTIDE_ANTIGEN | http://www.gsea-msigdb.org/gsea/msigdb/human/geneset/GOBP_ANTIGEN_PROCESSING_AND_PRESENTATION_OF_ENDOGENOUS_PEPTIDE_ANTIGEN | 23 | -0.39 | -1.29 | 7.14E-02 | 9.22E-01 | 6 |
| GOBP_ANTIGEN_PROCESSING_AND_PRESENTATION_VIA_MHC_CLASS_IB | http://www.gsea-msigdb.org/gsea/msigdb/human/geneset/GOBP_ANTIGEN_PROCESSING_AND_PRESENTATION_VIA_MHC_CLASS_IB | 21 | -0.41 | -1.32 | 7.27E-02 | 8.98E-01 | 7 |
| GOBP_ANTIGEN_PROCESSING_AND_PRESENTATION_OF_EXOGENOUS_ANTIGEN | http://www.gsea-msigdb.org/gsea/msigdb/human/geneset/GOBP_ANTIGEN_PROCESSING_AND_PRESENTATION_OF_EXOGENOUS_ANTIGEN | 26 | 0.33 | 1.16 | 7.41E-02 | 6.70E-01 | 7 |
| GOBP_ANTIGEN_PROCESSING_AND_PRESENTATION_OF_ENDOGENOUS_ANTIGEN | http://www.gsea-msigdb.org/gsea/msigdb/human/geneset/GOBP_ANTIGEN_PROCESSING_AND_PRESENTATION_OF_ENDOGENOUS_ANTIGEN | 26 | -0.36 | -1.13 | 2.04E-01 | 1.00E+00 | 10 |
| GOBP_INTERFERON_BETA_PRODUCTION | http://www.gsea-msigdb.org/gsea/msigdb/human/geneset/GOBP_INTERFERON_BETA_PRODUCTION | 48 | 0.42 | 1.05 | 2.60E-01 | 7.64E-01 | 22 |
| GOBP_REGULATION_OF_TYPE_I_INTERFERON_MEDIATED_SIGNALING_PATHWAY | http://www.gsea-msigdb.org/gsea/msigdb/human/geneset/GOBP_REGULATION_OF_TYPE_I_INTERFERON_MEDIATED_SIGNALING_PATHWAY | 37 | 0.40 | 0.95 | 5.00E-01 | 8.41E-01 | 13 |

| | | | | | | | |
|---|---|---|---|---|---|---|---|
| GOBP_INTERFERON_MEDIATED_SIGNALING_PATHWAY | http://www.gsea-msigdb.org/gsea/msigdb/human/geneset/GOBP_INTERFERON_MEDIATED_SIGNALING_PATHWAY | 69 | 0.30 | 0.84 | 5.80E-01 | 9.19E-01 | 8 |
| GOBP_ANTIGEN_PROCESSING_AND_PRESENTATION_OF_PEPTIDE_ANTIGEN_VIA_MHC_CLASS_I | http://www.gsea-msigdb.org/gsea/msigdb/human/geneset/GOBP_ANTIGEN_PROCESSING_AND_PRESENTATION_OF_PEPTIDE_ANTIGEN_VIA_MHC_CLASS_I | 33 | -0.27 | -0.92 | 6.04E-01 | 1.00E+00 | 8 |
| GOBP_ANTIGEN_PROCESSING_AND_PRESENTATION_OF_EXOGENOUS_PEPTIDE_ANTIGEN | http://www.gsea-msigdb.org/gsea/msigdb/human/geneset/GOBP_ANTIGEN_PROCESSING_AND_PRESENTATION_OF_EXOGENOUS_PEPTIDE_ANTIGEN | 21 | 0.29 | 0.92 | 6.30E-01 | 8.63E-01 | 6 |
| GOBP_TYPE_I_INTERFERON_PRODUCTION | http://www.gsea-msigdb.org/gsea/msigdb/human/geneset/GOBP_TYPE_I_INTERFERON_PRODUCTION | 85 | 0.37 | 1.03 | 6.40E-01 | 7.81E-01 | 36 |
| GOBP_RESPONSE_TO_VIRUS | http://www.gsea-msigdb.org/gsea/msigdb/human/geneset/GOBP_RESPONSE_TO_VIRUS | 289 | 0.24 | 0.81 | 7.00E-01 | 9.29E-01 | 68 |
| GOBP_ANTIGEN_PROCESSING_AND_PRESENTATION_OF_PEPTIDE_ANTIGEN | http://www.gsea-msigdb.org/gsea/msigdb/human/geneset/GOBP_ANTIGEN_PROCESSING_AND_PRESENTATION_OF_PEPTIDE_ANTIGEN | 47 | -0.20 | -0.86 | 7.41E-01 | 1.00E+00 | 9 |
| GOBP_POSITIVE_REGULATION_OF_INTERFERON_BETA_PRODUCTION | http://www.gsea-msigdb.org/gsea/msigdb/human/geneset/GOBP_POSITIVE_REGULATION_OF_INTERFERON_BETA_PRODUCTION | 33 | 0.38 | 0.89 | 7.80E-01 | 8.86E-01 | 14 |
| GOBP_NEGATIVE_REGULATION_OF_VIRAL_GENOME_REPLICATION | http://www.gsea-msigdb.org/gsea/msigdb/human/geneset/GOBP_NEGATIVE_REGULATION_OF_VIRAL_GENOME_REPLICATION | 46 | -0.25 | -0.67 | 8.24E-01 | 1.00E+00 | 6 |

| | | | | | | | |
|---|---|---|---|---|---|---|---|
| GOBP_NEGATIVE_REGULATION_OF_INNATE_IMMUNE_RESPONSE | http://www.gsea-msigdb.org/gsea/msigdb/human/geneset/GOBP_NEGATIVE_REGULATION_OF_INNATE_IMMUNE_RESPONSE | 56 | -0.22 | -0.78 | 8.39E-01 | 1.00E+00 | 19 |
| GOBP_POSITIVE_REGULATION_OF_TYPE_I_INTERFERON_PRODUCTION | http://www.gsea-msigdb.org/gsea/msigdb/human/geneset/GOBP_POSITIVE_REGULATION_OF_TYPE_I_INTERFERON_PRODUCTION | 53 | 0.34 | 0.88 | 8.40E-01 | 8.93E-01 | 23 |
| GOBP_NEGATIVE_REGULATION_OF_VIRAL_PROCESS | http://www.gsea-msigdb.org/gsea/msigdb/human/geneset/GOBP_NEGATIVE_REGULATION_OF_VIRAL_PROCESS | 69 | 0.18 | 0.52 | 9.80E-01 | 9.97E-01 | 29 |
| GOBP_RESPONSE_TO_TYPE_I_INTERFERON | http://www.gsea-msigdb.org/gsea/msigdb/human/geneset/GOBP_RESPONSE_TO_TYPE_I_INTERFERON | 61 | 0.29 | 0.69 | 1.00E+00 | 9.74E-01 | 23 |
| GOBP_ANTIGEN_PROCESSING_AND_PRESENTATION | http://www.gsea-msigdb.org/gsea/msigdb/human/geneset/GOBP_ANTIGEN_PROCESSING_AND_PRESENTATION | 75 | 0.15 | 0.76 | 1.00E+00 | 9.54E-01 | 8 |

Supplementary Table 10

Expressions of *IFNA1* and *IFNK* in ESCC PDO and cell line panels, and in K180 and K450 depleted of ΔNp63.

| | Expression in unmanipulated PDO/CLs | | Expression in manipulated K180 | | | | Expression in manipulated K450 | | | |
|---|---|---|---|---|---|---|---|---|---|---|
| Gene name | LSMean (ESCC PDO) | LSMean(2DCL-ESCC) | P-value | FDR | LSMean(Ctrl) | LSMean(p63fKO) | P-value | FDR | LSMean(Ctrl) | LSMean(p63fKO) |
| IFNA1 | 0.11 | 0.01 | 8.29E-01 | 9.46E-01 | 0.19 | 0.11 | 1.88E-01 | ####### | 0.38 | 0.40 |
| IFNK | 0.00 | 0.00 | 4.41E-01 | 6.40E-01 | 0.25 | 0.43 | 4.22E-01 | ####### | 0.16 | 0.53 |

Supplementary Table 11
Summary of differentially expressed endogenous retrotransposons by TEcounts.

| | *gene/TE* | *baseMean* | *log2FoldChange* | *lfcSE* | *stat* | *pvalue* | *padj* | *Log(Ctrl1)* | *Log(Ctrl2)* | *Log(p63fKO1)* | *Log(p63fKO2)* | *Types* |
|---|---|---|---|---|---|---|---|---|---|---|---|---|
| **KYSE180-p63fKO/Ctrl** | | | | | | | | | | | | |
| 1 | hAT-N1_Mam:hAT-Tip100:DNA | 47.826 | 3.9206 | 0.515 | 7.61 | 2.69E-14 | 1.64E-12 | 4.605 | 4.7 | 5.8578 | 5.84728 | DNA |
| 2 | Charlie2b:hAT-Charlie:DNA | 317.5 | 1.4609 | 0.139 | 10.5 | 9.28E-26 | 1.15E-23 | 7.61 | 7.521 | 8.7592 | 8.74812 | DNA |
| 3 | MER103C:hAT-Charlie:DNA | 100.7 | 0.9824 | 0.251 | 3.91 | 9.16E-05 | 0.001554 | 6.452 | 6.141 | 6.9105 | 6.86225 | DNA |
| 4 | MER30:hAT-Charlie:DNA | 294.49 | 0.5709 | 0.137 | 4.15 | 3.27E-05 | 0.000624 | 7.938 | 7.953 | 8.4042 | 8.41661 | DNA |
| 5 | MER75:PiggyBac:DNA | 225.04 | 0.5249 | 0.164 | 3.2 | 0.0013668 | 0.016609 | 7.541 | 7.638 | 7.8792 | 8.10518 | DNA |
| 6 | Tigger1:TcMar-Tigger:DNA | 571.67 | 0.395 | 0.102 | 3.87 | 0.0001107 | 0.001827 | 8.938 | 9.003 | 9.2976 | 9.347 | DNA |
| 7 | Tigger4b:TcMar-Tigger:DNA | 340.15 | -0.4626 | 0.128 | -3.6 | 0.0003123 | 0.004592 | 8.592 | 8.581 | 8.1765 | 8.22489 | DNA |
| 8 | MER102c:hAT-Charlie:DNA | 200.29 | -0.7365 | 0.168 | -4.37 | 1.23E-05 | 0.000257 | 7.833 | 7.934 | 7.3715 | 7.29304 | DNA |
| 9 | L1PREC2:L1:LINE | 496.07 | 2.4262 | 0.127 | 19.1 | 1.56E-81 | 9.25E-79 | 7.43 | 7.611 | 9.5574 | 9.61413 | LINE |
| 10 | L2-3_Crp:L2:LINE | 60.255 | 2.291 | 0.355 | 6.45 | 1.09E-10 | 4.69E-09 | 5.286 | 5.168 | 6.3142 | 6.0806 | LINE |
| 11 | L1PA15:L1:LINE | 459.96 | 1.6346 | 0.282 | 5.8 | 6.63E-09 | 2.39E-07 | 8.158 | 7.702 | 9.4213 | 9.27242 | LINE |
| 12 | L1MCc:L1:LINE | 48.979 | 1.1258 | 0.353 | 3.19 | 0.001411 | 0.017054 | 5.317 | 5.344 | 5.7039 | 5.87177 | LINE |
| 13 | L1P1:L1:LINE | 1905.3 | 1.0268 | 0.065 | 15.8 | 1.52E-56 | 4.92E-54 | 10.31 | 10.33 | 11.262 | 11.3323 | LINE |
| 14 | L1PA2:L1:LINE | 12676 | 0.8938 | 0.038 | 23.3 | 3.72E-120 | ######## | 13.17 | 13.07 | 14 | 14.0042 | LINE |
| 15 | L1HS:L1:LINE | 11010 | 0.891 | 0.038 | 23.6 | 1.18E-123 | ######## | 12.95 | 12.89 | 13.781 | 13.8137 | LINE |

| | | | | | | | | | | | | |
|---|---|---|---|---|---|---|---|---|---|---|---|---|
| 16 | L1PA3:L1:LINE | 13405 | 0.8144 | 0.038 | 21.5 | 1.18E-102 | ######## | 13.28 | 13.22 | 14.02 | 14.09 | LINE |
| 17 | L1P3:L1:LINE | 100.03 | 0.809 | 0.25 | 3.24 | 0.0011906 | 0.014741 | 6.507 | 6.203 | 6.8288 | 6.85724 | LINE |
| 18 | L1PA10:L1:LINE | 102.45 | 0.7856 | 0.235 | 3.34 | 0.0008496 | 0.01104 | 6.452 | 6.361 | 6.8678 | 6.88959 | LINE |
| 19 | L1PA4:L1:LINE | 14806 | 0.7742 | 0.073 | 10.5 | 5.94E-26 | 7.45E-24 | 13.49 | 13.35 | 14.195 | 14.1744 | LINE |
| 20 | L1PA5:L1:LINE | 4021.1 | 0.7155 | 0.102 | 7.03 | 2.07E-12 | 1.07E-10 | 11.68 | 11.48 | 12.266 | 12.2882 | LINE |
| 21 | L1MA5:L1:LINE | 202.71 | 0.7071 | 0.168 | 4.2 | 2.65E-05 | 0.000516 | 7.282 | 7.437 | 7.9154 | 7.87442 | LINE |
| 22 | L1PA7:L1:LINE | 1827 | 0.6856 | 0.073 | 9.4 | 5.32E-21 | 5.07E-19 | 10.57 | 10.36 | 11.117 | 11.1257 | LINE |
| 23 | L1MA8:L1:LINE | 226.73 | 0.6128 | 0.166 | 3.69 | 0.0002281 | 0.003502 | 7.535 | 7.584 | 7.889 | 8.16544 | LINE |
| 24 | L1PA6:L1:LINE | 810.8 | 0.6033 | 0.098 | 6.14 | 8.01E-10 | 3.15E-08 | 9.482 | 9.225 | 9.9312 | 9.88454 | LINE |
| 25 | L1M3:L1:LINE | 150.65 | 0.5977 | 0.193 | 3.1 | 0.0019261 | 0.022157 | 7.021 | 6.987 | 7.3674 | 7.468 | LINE |
| 26 | L1MEc:L1:LINE | 233.79 | 0.4978 | 0.161 | 3.09 | 0.0019823 | 0.022678 | 7.71 | 7.601 | 7.9282 | 8.15054 | LINE |
| 27 | L1MB8:L1:LINE | 479.87 | 0.4497 | 0.114 | 3.94 | 8.20E-05 | 0.001412 | 8.742 | 8.646 | 9.1602 | 9.00762 | LINE |
| 28 | L1MB5:L1:LINE | 414.55 | 0.4288 | 0.118 | 3.63 | 0.0002794 | 0.004172 | 8.47 | 8.524 | 8.9019 | 8.82752 | LINE |
| 29 | L1ME3E:L1:LINE | 380.78 | 0.3506 | 0.121 | 2.89 | 0.0038866 | 0.039805 | 8.4 | 8.429 | 8.7233 | 8.70108 | LINE |
| 30 | L1MC5a:L1:LINE | 1056.9 | 0.3076 | 0.084 | 3.64 | 0.0002711 | 0.004063 | 9.964 | 9.824 | 10.236 | 10.1214 | LINE |
| 31 | L1M5:L1:LINE | 1986.1 | 0.1921 | 0.062 | 3.1 | 0.0019324 | 0.022213 | 10.89 | 10.83 | 11.048 | 11.04 | LINE |
| 32 | L2c:L2:LINE | 4382.3 | 0.161 | 0.05 | 3.21 | 0.001319 | 0.016122 | 12.06 | 11.97 | 12.157 | 12.1896 | LINE |
| 33 | L1M4c:L1:LINE | 3194.2 | -0.3369 | 0.057 | -5.96 | 2.52E-09 | 9.48E-08 | 11.85 | 11.74 | 11.449 | 11.4873 | LINE |
| 34 | L1MB2:L1:LINE | 1359.1 | -0.3625 | 0.077 | -4.72 | 2.40E-06 | 5.74E-05 | 10.64 | 10.49 | 10.247 | 10.2043 | LINE |
| 35 | L4_A_Mam:RTE-X:LINE | 170.92 | -0.5782 | 0.18 | -3.21 | 0.0013423 | 0.016368 | 7.595 | 7.61 | 7.1402 | 7.23151 | LINE |
| 36 | HAL1:L1:LINE | 1053.2 | -0.6026 | 0.084 | -7.21 | 5.43E-13 | 2.97E-11 | 10.27 | 10.31 | 9.6474 | 9.81021 | LINE |
| 37 | MER72:ERV1:LTR | 67.375 | 1.8271 | 0.326 | 5.61 | 1.99E-08 | 6.69E-07 | 5.632 | 5.349 | 6.2974 | 6.40966 | LTR |
| 38 | LTR26:ERV1:LTR | 16.775 | 1.8269 | 0.642 | 2.85 | 0.0044346 | 0.044308 | 3.81 | 3.883 | 4.2262 | 4.10997 | LTR |
| 39 | LTR57:ERVL:LTR | 32.708 | 1.7656 | 0.448 | 3.94 | 8.08E-05 | 0.001394 | 4.662 | 4.692 | 5.2409 | 5.17227 | LTR |
| 40 | MER31B:ERV1:LTR | 36.37 | 1.6556 | 0.425 | 3.9 | 9.63E-05 | 0.00162 | 4.894 | 4.756 | 5.3571 | 5.37353 | LTR |

| | | | | | | | | | | | | |
|---|---|---|---|---|---|---|---|---|---|---|---|---|
| 41 | MER21C:ERVL:LTR | 397.14 | 1.3923 | 0.127 | 10.9 | 6.81E-28 | 9.27E-26 | 7.961 | 7.846 | 9.0244 | 9.13033 | LTR |
| 42 | MER34-int:ERV1:LTR | 54.772 | 1.1583 | 0.333 | 3.48 | 0.0004957 | 0.006884 | 5.499 | 5.429 | 5.9004 | 6.03261 | LTR |
| 43 | MER92B:ERV1:LTR | 92.577 | 0.8273 | 0.256 | 3.24 | 0.0012054 | 0.014889 | 6.303 | 6.206 | 6.8382 | 6.61226 | LTR |
| 44 | ERV3-16A3_I-int:ERVL:LTR | 647.24 | 0.7797 | 0.098 | 7.99 | 1.31E-15 | 8.79E-14 | 8.904 | 8.974 | 9.636 | 9.64384 | LTR |
| 45 | LTR16B2:ERVL:LTR | 94.878 | 0.7649 | 0.252 | 3.04 | 0.002385 | 0.026459 | 6.375 | 6.244 | 6.8556 | 6.64641 | LTR |
| 46 | MER4B-int:ERV1:LTR | 141.69 | 0.7168 | 0.198 | 3.61 | 0.0003036 | 0.004484 | 6.844 | 6.897 | 7.3596 | 7.35522 | LTR |
| 47 | HERVK9-int:ERVK:LTR | 158.48 | 0.6737 | 0.19 | 3.55 | 0.000382 | 0.005487 | 7.015 | 7.065 | 7.445 | 7.58189 | LTR |
| 48 | MLT1L:ERVL-MaLR:LTR | 487.71 | 0.5024 | 0.118 | 4.25 | 2.17E-05 | 0.000431 | 8.815 | 8.556 | 9.093 | 9.16231 | LTR |
| 49 | MLT1A0:ERVL-MaLR:LTR | 538.48 | 0.47 | 0.106 | 4.43 | 9.41E-06 | 0.000202 | 8.841 | 8.854 | 9.3182 | 9.20497 | LTR |
| 50 | LTR46-int:ERV1:LTR | 546.67 | 0.4601 | 0.105 | 4.38 | 1.21E-05 | 0.000254 | 8.902 | 8.846 | 9.2335 | 9.32697 | LTR |
| 51 | MSTB1:ERVL-MaLR:LTR | 438.28 | 0.4427 | 0.117 | 3.8 | 0.0001474 | 0.002358 | 8.598 | 8.541 | 9.0118 | 8.8875 | LTR |
| 52 | MER34B:ERV1:LTR | 276.7 | 0.4051 | 0.141 | 2.87 | 0.0040624 | 0.041204 | 7.944 | 7.93 | 8.2596 | 8.2669 | LTR |
| 53 | MLT1J:ERVL-MaLR:LTR | 760.8 | 0.321 | 0.09 | 3.58 | 0.000349 | 0.005064 | 9.401 | 9.432 | 9.6919 | 9.72658 | LTR |
| 54 | LTR12C:ERV1:LTR | 718.39 | 0.2972 | 0.093 | 3.2 | 0.0013743 | 0.016681 | 9.392 | 9.301 | 9.6167 | 9.61468 | LTR |
| 55 | MER34B-int:ERV1:LTR | 1616.4 | -0.2322 | 0.071 | -3.27 | 0.001094 | 0.013675 | 10.77 | 10.76 | 10.469 | 10.6152 | LTR |
| 56 | LTR7Y:ERV1:LTR | 938.48 | -0.3463 | 0.082 | -4.21 | 2.57E-05 | 0.000502 | 10.04 | 10.01 | 9.687 | 9.72147 | LTR |
| 57 | MER11B:ERVK:LTR | 12633 | -0.592 | 0.039 | -15.4 | 2.61E-53 | 7.91E-51 | 13.91 | 13.86 | 13.344 | 13.2609 | LTR |
| 58 | HERVH-int:ERV1:LTR | 8656.8 | -0.5956 | 0.043 | -13.7 | 1.07E-42 | 2.44E-40 | 13.39 | 13.29 | 12.742 | 12.7715 | LTR |
| 59 | MER65C:ERV1:LTR | 196.68 | -0.8536 | 0.175 | -4.88 | 1.06E-06 | 2.73E-05 | 7.818 | 7.952 | 7.153 | 7.3452 | LTR |

| | | | | | | | | | | | | |
|---|---|---|---|---|---|---|---|---|---|---|---|---|
| 60 | LTR45C:ERV1:LTR | 46.399 | -1.6277 | 0.394 | -4.14 | 3.55E-05 | 0.000669 | 5.673 | 5.815 | 5.2543 | 4.96906 | LTR |
| 61 | MER41C:ERV1:LTR | 39.745 | -2.3692 | 0.436 | -5.44 | 5.47E-08 | 1.74E-06 | 5.439 | 5.624 | 4.7558 | 4.77368 | LTR |
| 62 | SVA_F:SVA:Retroposon | 4606.6 | -0.1524 | 0.046 | -3.33 | 0.00088 | 0.011361 | 12.23 | 12.26 | 12.095 | 12.0922 | Retroposon |
| 63 | 7SK:RNA:RNA | 158499 | -0.1944 | 0.021 | -9.19 | 3.85E-20 | 3.51E-18 | 17.35 | 17.39 | 17.174 | 17.1744 | RNA |
| 64 | AluYm1:Alu:SINE | 322.73 | 0.3887 | 0.132 | 2.95 | 0.0031498 | 0.033424 | 8.136 | 8.186 | 8.4601 | 8.50745 | SINE |
| 65 | AluJo:Alu:SINE | 3651 | -0.186 | 0.05 | -3.7 | 0.0002144 | 0.003311 | 11.94 | 11.9 | 11.759 | 11.7226 | SINE |
| 66 | AluSq:Alu:SINE | 1649.9 | -0.2223 | 0.076 | -2.91 | 0.0036549 | 0.037864 | 10.88 | 10.69 | 10.529 | 10.6247 | SINE |
| 67 | AluJb:Alu:SINE | 11395 | -0.3594 | 0.036 | -10 | 9.74E-24 | 1.08E-21 | 13.62 | 13.66 | 13.281 | 13.2956 | SINE |
| 68 | AluYj4:Alu:SINE | 300.95 | -0.4101 | 0.14 | -2.92 | 0.0034874 | 0.036428 | 8.349 | 8.425 | 7.9556 | 8.14465 | SINE |
| **KYSE450-p63fKO/Ctrl** | | | | | | | | | | | | |
| 1 | Charlie4z:hAT-Charlie:DNA | 63.821 | 3.9912 | 0.453 | 8.81 | 1.22E-18 | 5.21E-17 | 4.736 | 4.689 | 6.4446 | 6.32539 | DNA |
| 2 | MER117:hAT-Charlie:DNA | 15.559 | 3.8583 | 0.893 | 4.32 | 1.55E-05 | 0.000184 | 3.51 | 3.464 | 4.1481 | 4.05161 | DNA |
| 3 | MamTip3:hAT-Tip100:DNA | 27.357 | 2.3331 | 0.518 | 4.5 | 6.75E-06 | 8.56E-05 | 4.292 | 4.267 | 5.0402 | 4.92489 | DNA |
| 4 | Charlie18a:hAT-Charlie:DNA | 97.617 | 1.9879 | 0.262 | 7.59 | 3.26E-14 | 1.02E-12 | 5.805 | 5.764 | 7.0285 | 7.00173 | DNA |
| 5 | MamRep137:TcMar-Tigger:DNA | 75.363 | 1.583 | 0.295 | 5.37 | 7.99E-08 | 1.36E-06 | 5.705 | 5.607 | 6.6638 | 6.43841 | DNA |
| 6 | MER91B:hAT-Tip100:DNA | 26.824 | 1.3491 | 0.475 | 2.84 | 0.0044831 | 0.029753 | 4.478 | 4.464 | 4.9052 | 4.91103 | DNA |
| 7 | MER30:hAT-Charlie:DNA | 253.25 | 1.2752 | 0.152 | 8.4 | 4.50E-17 | 1.77E-15 | 7.392 | 7.286 | 8.4234 | 8.36588 | DNA |
| 8 | Charlie9:hAT-Charlie:DNA | 391.25 | 1.0728 | 0.12 | 8.92 | 4.62E-19 | 2.04E-17 | 8.061 | 8.041 | 9.0021 | 8.99107 | DNA |

| | | | | | | | | | | | | |
|---|---|---|---|---|---|---|---|---|---|---|---|---|
| 9 | MER63B:hAT-Blackjack:DNA | 90.544 | 0.7229 | 0.255 | 2.83 | 0.0046551 | 0.0307 | 6.339 | 6.134 | 6.7538 | 6.63862 | DNA |
| 10 | MER5B:hAT-Charlie:DNA | 127.42 | 0.6607 | 0.215 | 3.07 | 0.0021209 | 0.015601 | 6.59 | 6.856 | 7.2097 | 7.18556 | DNA |
| 11 | Charlie8:hAT-Charlie:DNA | 220.95 | 0.4902 | 0.157 | 3.11 | 0.0018499 | 0.013837 | 7.633 | 7.504 | 7.9923 | 7.94728 | DNA |
| 12 | MER1B:hAT-Charlie:DNA | 220.33 | 0.4274 | 0.157 | 2.72 | 0.0064479 | 0.0403 | 7.537 | 7.653 | 7.9354 | 7.95209 | DNA |
| 13 | Charlie6:hAT-Charlie:DNA | 312.56 | -0.4489 | 0.133 | -3.38 | 0.0007223 | 0.006067 | 8.467 | 8.463 | 8.0048 | 8.1499 | DNA |
| 14 | Tigger4b:TcMar-Tigger:DNA | 507.28 | -0.6121 | 0.108 | -5.67 | 1.41E-08 | 2.63E-07 | 9.25 | 9.219 | 8.7643 | 8.58561 | DNA |
| 15 | L1MDb:L1:LINE | 126.82 | 3.2144 | 0.273 | 11.8 | 6.04E-32 | 5.05E-30 | 5.571 | 5.438 | 7.5325 | 7.50475 | LINE |
| 16 | L1PREC2:L1:LINE | 355.53 | 2.8952 | 0.153 | 18.9 | 1.09E-79 | 3.39E-77 | 6.731 | 6.762 | 9.1417 | 9.16376 | LINE |
| 17 | L1PA10:L1:LINE | 157.34 | 1.9007 | 0.204 | 9.32 | 1.13E-20 | 5.42E-19 | 6.447 | 6.312 | 7.756 | 7.78201 | LINE |
| 18 | L1M4a1:L1:LINE | 15.36 | 1.8972 | 0.67 | 2.83 | 0.0046451 | 0.030649 | 3.734 | 3.622 | 4.1178 | 4.01055 | LINE |
| 19 | L1PBa1:L1:LINE | 25.918 | 1.5725 | 0.498 | 3.16 | 0.001579 | 0.01207 | 4.432 | 4.32 | 4.9316 | 4.80015 | LINE |
| 20 | L2-3_Crp:L2:LINE | 206.31 | 1.5138 | 0.171 | 8.88 | 6.90E-19 | 3.01E-17 | 6.892 | 6.977 | 8.1479 | 8.11303 | LINE |
| 21 | L1PA15:L1:LINE | 240.24 | 0.9704 | 0.159 | 6.11 | 9.93E-10 | 2.12E-08 | 7.572 | 7.291 | 8.2423 | 8.23676 | LINE |
| 22 | L1P3:L1:LINE | 149.57 | 0.8246 | 0.212 | 3.88 | 0.0001037 | 0.001049 | 6.98 | 6.751 | 7.6541 | 7.28684 | LINE |
| 23 | L1MC5a:L1:LINE | 861.51 | 0.8039 | 0.084 | 9.58 | 1.00E-21 | 5.05E-20 | 9.273 | 9.367 | 10.099 | 10.0511 | LINE |
| 24 | L1MD:L1:LINE | 86.496 | 0.7096 | 0.261 | 2.71 | 0.006637 | 0.041256 | 6.125 | 6.242 | 6.7209 | 6.51834 | LINE |
| 25 | L1HS:L1:LINE | 5527 | 0.6366 | 0.04 | 15.9 | 5.06E-57 | 9.54E-55 | 12.09 | 12.07 | 12.73 | 12.6924 | LINE |
| 26 | L1PA2:L1:LINE | 7521.2 | 0.5681 | 0.036 | 15.8 | 5.01E-56 | 9.18E-54 | 12.58 | 12.55 | 13.142 | 13.1169 | LINE |
| 27 | L1PA3:L1:LINE | 7660.7 | 0.4967 | 0.036 | 13.9 | 1.26E-43 | 1.52E-41 | 12.64 | 12.63 | 13.145 | 13.1098 | LINE |

| | | | | | | | | | | | | |
|---|---|---|---|---|---|---|---|---|---|---|---|---|
| 28 | MamRTE1:RTE-Bo.B:LINE | 755.68 | 0.4953 | 0.091 | 5.47 | 4.59E-08 | 7.99E-07 | 9.223 | 9.394 | 9.761 | 9.78344 | LINE |
| 29 | L1P1:L1:LINE | 1009.7 | 0.4656 | 0.081 | 5.77 | 7.99E-09 | 1.54E-07 | 9.663 | 9.816 | 10.217 | 10.1459 | LINE |
| 30 | L1PA6:L1:LINE | 515.68 | 0.4621 | 0.104 | 4.43 | 9.49E-06 | 0.000118 | 8.824 | 8.741 | 9.2343 | 9.17175 | LINE |
| 31 | L1M5:L1:LINE | 670.27 | 0.4391 | 0.092 | 4.79 | 1.70E-06 | 2.39E-05 | 9.171 | 9.167 | 9.5478 | 9.60482 | LINE |
| 32 | L2c:L2:LINE | 3934.8 | 0.4258 | 0.046 | 9.32 | 1.20E-20 | 5.76E-19 | 11.72 | 11.71 | 12.103 | 12.1679 | LINE |
| 33 | L1PA5:L1:LINE | 2542.5 | 0.4067 | 0.097 | 4.21 | 2.61E-05 | 0.000297 | 11.19 | 11 | 11.499 | 11.4937 | LINE |
| 34 | L1ME3A:L1:LINE | 343.33 | 0.3859 | 0.128 | 3.02 | 0.0024902 | 0.01796 | 8.253 | 8.232 | 8.5043 | 8.65287 | LINE |
| 35 | L1PA4:L1:LINE | 7566.5 | 0.3685 | 0.037 | 9.84 | 7.77E-23 | 4.21E-21 | 12.7 | 12.69 | 13.088 | 13.0228 | LINE |
| 36 | L1ME3E:L1:LINE | 511.1 | 0.3587 | 0.108 | 3.32 | 0.0008962 | 0.007367 | 8.908 | 8.734 | 9.1946 | 9.10466 | LINE |
| 37 | L2a:L2:LINE | 4139.7 | -0.3359 | 0.045 | -7.52 | 5.66E-14 | 1.73E-12 | 12.18 | 12.16 | 11.808 | 11.8728 | LINE |
| 38 | LTR43-int:ERV1:LTR | 5.6077 | 5.9163 | 1.719 | 3.44 | 0.0005775 | 0.004977 | 2.271 | 2.274 | 2.5366 | 2.53468 | LTR |
| 39 | LOR1a:ERV1:LTR | 15.874 | 3.8668 | 0.896 | 4.32 | 1.60E-05 | 0.000189 | 3.437 | 3.57 | 4.1667 | 4.08852 | LTR |
| 40 | MER31A:ERV1:LTR | 38.041 | 2.8891 | 0.477 | 6.06 | 1.34E-09 | 2.82E-08 | 4.541 | 4.466 | 5.5838 | 5.46371 | LTR |
| 41 | MER72:ERV1:LTR | 74.777 | 2.1001 | 0.306 | 6.87 | 6.54E-12 | 1.73E-10 | 5.413 | 5.468 | 6.5318 | 6.6498 | LTR |
| 42 | LTR57:ERVL:LTR | 23.335 | 1.8499 | 0.536 | 3.45 | 0.0005562 | 0.004828 | 4.129 | 4.254 | 4.7448 | 4.68425 | LTR |
| 43 | MER21C:ERVL:LTR | 332.79 | 1.4476 | 0.133 | 10.9 | 1.65E-27 | 1.11E-25 | 7.602 | 7.604 | 8.8515 | 8.84159 | LTR |
| 44 | LTR7Y:ERV1:LTR | 2508.5 | 1.3008 | 0.054 | 24.1 | 1.29E-128 | ######## | 10.51 | 10.53 | 11.788 | 11.7841 | LTR |
| 45 | HERVH-int:ERV1:LTR | 13143 | 1.164 | 0.082 | 14.1 | 3.34E-45 | 4.35E-43 | 13.06 | 12.92 | 14.184 | 14.1044 | LTR |
| 46 | LTR6A:ERV1:LTR | 178.8 | 0.9189 | 0.177 | 5.19 | 2.13E-07 | 3.43E-06 | 7.038 | 7.096 | 7.7432 | 7.81503 | LTR |
| 47 | MER34C2:ERV1:LTR | 83.135 | 0.8612 | 0.262 | 3.29 | 0.0010147 | 0.008204 | 6.061 | 6.086 | 6.6067 | 6.58756 | LTR |
| 48 | LTR5B:ERVK:LTR | 102.55 | 0.8001 | 0.236 | 3.39 | 0.0007106 | 0.005978 | 6.304 | 6.446 | 6.9005 | 6.90786 | LTR |
| 49 | LTR7:ERV1:LTR | 566.7 | 0.7845 | 0.1 | 7.82 | 5.16E-15 | 1.74E-13 | 8.766 | 8.711 | 9.4804 | 9.43144 | LTR |
| 50 | MSTB:ERVL-MaLR:LTR | 132.67 | 0.7841 | 0.211 | 3.72 | 0.0002014 | 0.001919 | 6.845 | 6.602 | 7.329 | 7.25574 | LTR |

| | | | | | | | | | | | | |
|---|---|---|---|---|---|---|---|---|---|---|---|---|
| 51 | LTR16:ERVL:LTR | 108.48 | 0.7434 | 0.238 | 3.13 | 0.00177 | 0.013332 | 6.304 | 6.625 | 6.9497 | 6.99803 | LTR |
| 52 | MSTD:ERVL-MaLR:LTR | 150.23 | 0.5829 | 0.196 | 2.97 | 0.0029646 | 0.020881 | 7.039 | 6.94 | 7.5247 | 7.31681 | LTR |
| 53 | HERVK3-int:ERVK:LTR | 146.72 | 0.5827 | 0.193 | 3.02 | 0.0025148 | 0.018123 | 6.966 | 6.951 | 7.3983 | 7.38184 | LTR |
| 54 | MER11B:ERVK:LTR | 248.94 | 0.4714 | 0.147 | 3.2 | 0.0013693 | 0.010663 | 7.774 | 7.719 | 8.174 | 8.10384 | LTR |
| 55 | MLT1B:ERVL-MaLR:LTR | 337.6 | 0.4697 | 0.133 | 3.52 | 0.0004336 | 0.003848 | 8.3 | 8.04 | 8.5603 | 8.61332 | LTR |
| 56 | MER11C:ERVK:LTR | 444.76 | 0.4618 | 0.113 | 4.1 | 4.21E-05 | 0.000461 | 8.565 | 8.582 | 9.0506 | 8.91983 | LTR |
| 57 | LTR12C:ERV1:LTR | 420.14 | 0.4593 | 0.114 | 4.03 | 5.63E-05 | 0.000602 | 8.46 | 8.527 | 8.9188 | 8.8863 | LTR |
| 58 | MLT1K:ERVL-MaLR:LTR | 270.96 | 0.3925 | 0.143 | 2.75 | 0.0059106 | 0.037524 | 7.956 | 7.849 | 8.1829 | 8.28809 | LTR |
| 59 | MER65C:ERV1:LTR | 447.34 | -0.7927 | 0.115 | -6.91 | 4.99E-12 | 1.33E-10 | 9.189 | 9.026 | 8.4057 | 8.39543 | LTR |
| 60 | MLT1J1:ERVL-MaLR:LTR | 104.01 | -0.937 | 0.238 | -3.94 | 8.26E-05 | 0.000853 | 6.866 | 7.036 | 6.279 | 6.38663 | LTR |
| 61 | HUERS-P3-int:ERV1:LTR | 83.602 | -1.1153 | 0.267 | -4.17 | 3.03E-05 | 0.000342 | 6.694 | 6.604 | 5.893 | 6.05289 | LTR |
| 62 | LTR21A:ERV1:LTR | 462.86 | -1.2585 | 0.114 | -11.1 | 1.45E-28 | 1.04E-26 | 9.251 | 9.339 | 8.2017 | 8.13221 | LTR |
| 63 | MER34-int:ERV1:LTR | 334.34 | -1.3604 | 0.132 | -10.3 | 8.79E-25 | 5.29E-23 | 8.84 | 8.825 | 7.6113 | 7.71063 | LTR |
| 64 | LTR30:ERV1:LTR | 24.14 | -1.4284 | 0.507 | -2.82 | 0.0048731 | 0.031944 | 4.772 | 4.715 | 4.3756 | 4.24934 | LTR |
| 65 | LTR45B:ERV1:LTR | 70.604 | -1.8237 | 0.307 | -5.93 | 2.97E-09 | 6.07E-08 | 6.419 | 6.525 | 5.5535 | 5.39628 | LTR |
| 66 | SVA_C:SVA:Retroposon | 213.96 | 0.4777 | 0.168 | 2.85 | 0.0043761 | 0.029168 | 7.675 | 7.366 | 7.9035 | 7.93278 | Retroposon |
| 67 | SVA_D:SVA:Retroposon | 638.45 | 0.3084 | 0.093 | 3.3 | 0.0009567 | 0.007788 | 9.176 | 9.16 | 9.4307 | 9.47586 | Retroposon |
| 68 | 7SK:RNA:RNA | 181873 | 0.1023 | 0.022 | 4.74 | 2.16E-06 | 2.98E-05 | 17.41 | 17.43 | 17.502 | 17.5433 | RNA |
| 69 | FAM:Alu:SINE | 352.25 | 6.7324 | 0.414 | 16.3 | 2.17E-59 | 4.44E-57 | 5.082 | 5.286 | 9.1769 | 9.2646 | SINE |
| 70 | MIR1_Amn:MIR:SINE | 116.83 | 0.6902 | 0.219 | 3.15 | 0.0016487 | 0.01253 | 6.545 | 6.652 | 7.0995 | 7.05185 | SINE |
| 71 | MIR3:MIR:SINE | 512.39 | 0.5631 | 0.109 | 5.19 | 2.15E-07 | 3.46E-06 | 8.819 | 8.611 | 9.2306 | 9.23339 | SINE |
| 72 | AluYb8:Alu:SINE | 200.44 | 0.5273 | 0.168 | 3.14 | 0.0016994 | 0.012873 | 7.441 | 7.392 | 7.9291 | 7.73863 | SINE |

| | | | | | | | | | | | | |
|---|---|---|---|---|---|---|---|---|---|---|---|---|
| 73 | AluJo:Alu:SINE | 1810.6 | 0.4912 | 0.06 | 8.21 | 2.29E-16 | 8.59E-15 | 10.58 | 10.55 | 11.024 | 11.0554 | SINE |
| 74 | AluSc8:Alu:SINE | 598.12 | 0.4253 | 0.096 | 4.42 | 9.65E-06 | 0.000119 | 9.004 | 9.024 | 9.4011 | 9.40939 | SINE |
| 75 | AluY:Alu:SINE | 3313.2 | 0.3432 | 0.054 | 6.36 | 1.96E-10 | 4.49E-09 | 11.54 | 11.49 | 11.919 | 11.7814 | SINE |
| 76 | AluSc:Alu:SINE | 1482.7 | 0.2569 | 0.065 | 3.93 | 8.54E-05 | 0.000879 | 10.39 | 10.42 | 10.622 | 10.6812 | SINE |
| 77 | AluSq2:Alu:SINE | 2440.7 | 0.2361 | 0.055 | 4.32 | 1.57E-05 | 0.000186 | 11.15 | 11.12 | 11.403 | 11.3224 | SINE |
| 78 | AluSz:Alu:SINE | 3912.4 | 0.2159 | 0.044 | 4.91 | 9.29E-07 | 1.37E-05 | 11.82 | 11.83 | 12.042 | 12.03 | SINE |
| 79 | AluSg:Alu:SINE | 1450.8 | 0.2024 | 0.065 | 3.1 | 0.0019353 | 0.014392 | 10.39 | 10.41 | 10.575 | 10.6174 | SINE |
| 80 | MIR:MIR:SINE | 1948.5 | 0.1556 | 0.058 | 2.68 | 0.0073512 | 0.044875 | 10.87 | 10.83 | 11.011 | 10.9923 | SINE |
| 81 | AluJr:Alu:SINE | 2347.4 | 0.1515 | 0.055 | 2.78 | 0.005511 | 0.035356 | 11.13 | 11.12 | 11.301 | 11.235 | SINE |
| 82 | AluJb:Alu:SINE | 3431.5 | 0.1435 | 0.046 | 3.1 | 0.001954 | 0.014504 | 11.69 | 11.66 | 11.813 | 11.8139 | SINE |
| 83 | AluSq:Alu:SINE | 710.59 | -0.3114 | 0.09 | -3.48 | 0.0005041 | 0.004409 | 9.574 | 9.646 | 9.3349 | 9.30383 | SINE |

Supplementary Table 12

Summary of the expressions of RISC component genes in ΔNp63-depleted cell lines.

| *Gene name* | *P-value* | *FDR* | *LSMean(Ctrl)* | *LSMean(p63fKO)* | *LSMean(KYSE180-CTRL)* | *LSMean(KYSE180-p63fKO)* | *P-value* | *FDR* | *LSMean(Ctrl)* | *LSMean(p63fKO)* | *LSMean(KYSE450-CTRL)* | *LSMean(KYSE450-p63fKO)* |
|---|---|---|---|---|---|---|---|---|---|---|---|---|
| AGO1 | ####### | 6.11E-01 | 0.97 | -1.03 | 135.30 | 139.84 | 2.43E-03 | 1.29E-02 | 0.84 | -1.20 | 78.68 | 94.07 |
| AGO2 | ####### | 3.64E-02 | 0.87 | -1.15 | 178.39 | 204.92 | 1.14E-02 | 4.37E-02 | 1.14 | 1.14 | 157.48 | 138.74 |
| AGO3 | ####### | 5.97E-02 | 0.90 | -1.12 | 50.19 | 56.03 | 7.00E-01 | 8.56E-01 | 1.01 | 1.01 | 70.61 | 69.67 |
| AGO4 | ####### | 3.25E-01 | 1.10 | 1.10 | 30.49 | 27.76 | 5.84E-03 | 2.58E-02 | 0.79 | -1.27 | 26.60 | 33.66 |
| DICER1 | ####### | 7.78E-02 | 0.89 | -1.13 | 135.88 | 153.48 | 1.65E-01 | 3.22E-01 | 0.94 | -1.06 | 99.05 | 105.43 |
| TARBP2 | ####### | 9.94E-01 | 0.98 | -1.02 | 11.01 | 11.20 | 8.05E-02 | 1.97E-01 | 0.79 | -1.26 | 13.86 | 17.44 |

Supplementary Table 13

The transcription factor target geneset enrichments by GSEA in macrophges from cancer cell-TP63 high patient samples vs those from TP63 low samples show the enrichment of TFs related to macrophage polarization. Shown are top 5 significant TF target genesets.

| *Gene Set Name [# Genes]* | *# Genes in Overlap* | *p-value* | *FDR* | *Remarks* |
|---|---|---|---|---|
| **from macrophage genes negatively correlated with cancer cell TP63 expression (N=99)** | | | | |
| SETD7_TARGET_GENES [1002] | 16 | 1.47E-09 | 1.64E-06 | Macrophage SETD7 is involved in M1 macrophage activation and inflammatory mediator expression by promoting NF-κB signaling. |
| IRF_Q6 [244] | 9 | 6.46E-09 | 3.60E-06 | Macrophage IRF1 expression is required for M1 macrophage polarization and the antitumor functions. |
| ATF6_TARGET_GENES [1080] | 14 | 2.20E-07 | 6.26E-05 | Macrophage ATF6 expression faciliates inflammatory mediator expression and secretion. |
| CREB3L4_TARGET_GENES [1441] | 16 | 2.25E-07 | 6.26E-05 | Expression of CREB3L4 is classified in TNF signaling pathway in M1 polarization; no functional analysis found. |
| BARX1_TARGET_GENES [1285] | 14 | 1.73E-06 | 3.38E-04 | No study related to macrophage found. |
| **from macrophage genes positively correlated with cancer cell TP63 expression (N=33)** | | | | |
| PAX4_01 [268] | 5 | 1.97E-06 | 2.20E-03 | No study related to macrophage found. |
| MYCMAX_03 [264] | 4 | 5.17E-05 | 1.59E-02 | Inhibition of MYC/MAX dimerization leads to suppression of macrophage M2 alternative activation genes. |
| PRKDC_TARGET_GENES [887] | 6 | 5.56E-05 | 1.59E-02 | No study related to macrophage in cancer found. |
| CACCCBINDINGFACTOR_Q6 [271] | 4 | 5.72E-05 | 1.59E-02 | No study related to macrophage found. |
| ZNF407_TARGET_GENES [1996] | 8 | 1.13E-04 | 1.97E-02 | No study related to macrophage found. |

Supplementary Table 14

Summary of the expressions of genes encoding MHC class I molecules and antigen presenting proteins in ΔNp63-depleted cell lines. Highlighted are genes showing significant (FDR<0.05 in both cell lines) up-regulations upon ΔNp63 depletion.

| *Gene name* | *P-value* | *FDR* | *LSMean(Ctrl)* | *LSMean(p63fKO)* | *LSMean(KYSE180-CTRL)* | *LSMean(KYSE180-p63fKO)* | *P-value* | *FDR* | *LSMean(Ctrl)* | *LSMean(p63fKO)* | *LSMean(KYSE450-CTRL)* | *LSMean(KYSE450-p63fKO)* |
|---|---|---|---|---|---|---|---|---|---|---|---|---|
| HLA-A | 4.24E-06 | 2.08E-04 | 0.76 | -1.31 | 476.45 | 624.72 | 8.07E-07 | 4.22E-05 | 0.70 | -1.42 | 394.84 | 561.61 |
| HLA-B | 6.55E-06 | 2.75E-04 | 0.61 | -1.64 | 193.74 | 317.17 | 3.08E-08 | 5.76E-06 | 0.32 | -3.15 | 76.15 | 239.82 |
| HLA-C | 1.00E-06 | 8.04E-05 | 0.69 | -1.45 | 268.92 | 390.69 | 4.10E-09 | 2.37E-06 | 0.41 | -2.41 | 117.47 | 283.20 |
| HLA-DMA | 5.80E-02 | 1.72E-01 | 0.31 | -3.28 | 0.37 | 1.22 | 2.55E-01 | 4.26E-01 | 0.55 | -1.83 | 0.82 | 1.49 |
| HLA-DRA | 2.90E-01 | 4.85E-01 | 0.00 | ##### | 0.00 | 0.05 | 1.57E-01 | 3.13E-01 | 0.12 | -8.03 | 0.05 | 0.43 |
| HLA-E | 3.52E-01 | 5.49E-01 | 0.97 | -1.04 | 167.14 | 173.16 | 3.74E-06 | 1.17E-04 | 0.60 | -1.66 | 137.98 | 228.47 |
| HLA-F | 1.29E-05 | 4.28E-04 | 0.60 | -1.66 | 35.88 | 59.40 | 4.24E-07 | 2.68E-05 | 0.42 | -2.38 | 19.78 | 47.05 |
| HLA-H | 3.47E-01 | 5.44E-01 | 0.77 | -1.31 | 5.75 | 7.51 | 7.61E-02 | 1.89E-01 | 0.59 | -1.68 | 2.88 | 4.85 |
| HLA-J | 3.02E-01 | 4.97E-01 | 0.64 | -1.57 | 0.99 | 1.55 | 7.03E-02 | 1.78E-01 | 0.35 | -2.82 | 0.31 | 0.88 |
| HLA-K | 8.12E-01 | 9.36E-01 | 0.74 | -1.35 | 1.73 | 2.33 | 1.00E+00 | 1.00E+00 | 1.00 | -1.00 | 0.00 | 0.00 |
| HLA-L | 6.88E-01 | 8.57E-01 | 0.89 | -1.12 | 5.58 | 6.25 | 1.16E-01 | 2.55E-01 | 0.56 | -1.79 | 1.74 | 3.12 |
| HLA-V | 2.29E-03 | 1.54E-02 | 1.34 | 1.34 | 31.57 | 23.64 | 1.04E-06 | 4.90E-05 | 0.35 | -2.85 | 13.23 | 37.71 |
| TAP1 | 3.26E-06 | 1.78E-04 | 0.58 | -1.74 | 40.94 | 71.05 | 2.04E-06 | 7.76E-05 | 0.55 | -1.81 | 28.99 | 52.51 |
| TAP2 | 1.32E-05 | 4.34E-04 | 0.65 | -1.53 | 85.76 | 131.00 | 2.95E-04 | 2.56E-03 | 0.78 | -1.29 | 43.01 | 55.42 |

Supplementary Table 15

The ISGs and genes regulated by ΔNp63 in ESCC cell lines and xenografts show negative correlations with TP63 expression in CCLE TP63+ lung SCC cell lines (CALU1, SW900, SKMES1, EPLC272H, KSN62, HARA, LUDLU1, LC1F, LC1SQSF, HCC95, and HCC2814). Bolded are representative genes shown in Fig. 7E.

| *Gene category* | *Gene* | *Correlation to TP63* | *p-value* | *False Discovery Rate* |
|---|---|---|---|---|
| ISGs | IFIT1 | -0.841587315 | 0.00116129 | 0.004132703 |
| | IFIT2 | -0.840400347 | 0.001198796 | 0.004132703 |
| | **IFIT3** | -0.90058874 | 0.000155913 | 0.004132703 |
| | ISG15 | -0.846641418 | 0.001011365 | 0.004132703 |
| | ISG20 | -0.819991629 | 0.001996986 | 0.005222886 |
| | OAS1 | -0.733758198 | 0.010159043 | 0.01817934 |
| | OAS2 | -0.86888871 | 0.000516664 | 0.004132703 |
| | OASL | -0.839880637 | 0.001215501 | 0.004132703 |
| | MX1 | -0.824682724 | 0.001786042 | 0.005060452 |
| | MX2 | -0.814968908 | 0.002242896 | 0.005447033 |
| | NLRC5 | -0.768211422 | 0.005749322 | 0.012217309 |
| | RSAD2 | -0.594732052 | 0.053620089 | 0.072923321 |
| | SAMHD1 | -0.781803214 | 0.004474439 | 0.010142062 |
| | USP18 | -0.000600087 | 0.998602873 | 0.998602873 |
| | XAF1 | -0.831228585 | 0.001520104 | 0.004698503 |
| dsRNA sensors | TLR3 | -0.462305706 | 0.152233196 | 0.184854595 |
| | IFIH1 | -0.701243143 | 0.016198431 | 0.026226031 |
| | DDX58 | -0.761362985 | 0.006484236 | 0.012248001 |
| IRF1 and targets | IRF1 | -0.548774012 | 0.080425695 | 0.105172063 |
| | APOL1 | -0.655011549 | 0.028711692 | 0.042443371 |
| | APOL6 | -0.762178028 | 0.006393348 | 0.012248001 |
| | PLAAT4 | -0.845705023 | 0.001037976 | 0.004132703 |
| | PSMB9 | -0.388812071 | 0.237281803 | 0.278192459 |
| | TRIM22 | -0.854965008 | 0.000796762 | 0.004132703 |
| | UBA7 | -0.888417401 | 0.000257471 | 0.004132703 |
| | UBE2L6 | -0.668700085 | 0.024471018 | 0.037818846 |
| MHCI molecules | HLA-A | -0.319812354 | 0.337690098 | 0.382715444 |
| | **HLA-B** | -0.636621839 | 0.035189375 | 0.049851615 |
| | HLA-C | -0.469835646 | 0.144793547 | 0.182332615 |
| | HLA-F | -0.071541735 | 0.834426642 | 0.859712298 |
| Known ΔNp63-induced genes as controls | KRT15 | 0.844923314 | 0.001060591 | 0.004132703 |
| | HAS3 | 0.71427833 | 0.013532234 | 0.023004798 |
| House keeping genes as controls | HSPA4 | -0.150873053 | 0.6579063 | 0.699025444 |
| | THOC1/p84 | 0.160138712 | 0.638105653 | 0.699025444 |

Supplementary Table 16
Primers used in the study.

| *Target* | *Forward oligo* | *Reverse oligo* | *Application* |
|---|---|---|---|
| *OAS1* | TGTGTGTCCAAGGTGGTAAAG | GGTGAGAGGACTGAGGAAGA | Quantitative PCR |
| *OAS2* | AGCTGAGAGCAATGGGAAAT | CTTCGTAGGGCTTCAGGTATTC | Quantitative PCR |
| *OASL* | GCCATGTACTCCAGAACTCATC | GGCCTGGGATAACTCATTGTAA | Quantitative PCR |
| *RSAD2* | TGGTGAGGTTCTGCAAAGTAG | GTCACAGGAGATAGCGAGAATG | Quantitative PCR |
| *SAMHD1* | GTGTTCAGATTGCTGGACTTTG | CCTTGTTCATGCGTCCATTTC | Quantitative PCR |
| *MX1* | CTGGTGCTGAAACTGAAGAAAC | TACCTCTGAAGCATCCGAAATC | Quantitative PCR |
| *ISG20* | CCTACACAAGAGCATCCAGAAC | CTCGGATTCTCTGGGAGATTTG | Quantitative PCR |
| *ERVL* | ATATCCTGCCTGGATGGGGT | GAGCTTCTTAGTCCTCCTGTGT | Quantitative PCR |
| *HERV-E* | GGTGTCACTACTCAATACAC | GCAGCCTAGGTCTCTGG | Quantitative PCR |
| *HERV-F* | CCTCCAGTCACAACAACTC | TATTGAAGAAGGCGGCTGG | Quantitative PCR |
| *HERV-K* | ATTGGCAACACCGTATTCTGCT | CAGTCAAAATATGGACGGATGGT | Quantitative PCR |
| *L1-5UTR* | AAGCAAGCCTGGGCAATG | ACGGAATCTCGCTGATTGCTA | Quantitative PCR |
| *L1-ORF1* | TGGCCCCCACTCTCTTCT | TCAAAGGAAAGCCCATCAGACTA | Quantitative PCR |
| *MDL1-5UTR-B* | CGAGATCAAACTGCAAGGCG | CCGGCCGCTTTGTTTACCTA | Quantitative PCR |
| *MDL1-ORF1* | ACCTGAAAGTGACGGGGAGA | CCTGCCTTGCTAGATTGGGG | Quantitative PCR |
| *MDL1-ORF2* | CAAACACCGCATATTCTCACTCA | CTTCCTGTGTCCATGTGATCTCA | Quantitative PCR |
| *LTR12* | GAAACTCCGGACACATCTGAA | GTTCTTGGTCTCGCTGACTT | Quantitative PCR |
| *LTR26E* | CCAGTTTGAAGACCCCCACA | GGGTGAAGTCATGGGACAGG | Quantitative PCR |
| *LTR82B* | CATTGGTGAAATGAGTGGTGTT C | GGGCTCGCCAGATTCTTATAC | Quantitative PCR |
| *MER50B* | CGCCTCTTTCCCATGATTCT | CGCCTCTTTCCCATGATTCT | Quantitative PCR |
| *MER54A* | CACCCAAGCAGGTTTCTCAT | CGTAGGACAGCTGGACACAA | Quantitative PCR |
| *TLR3* | CCCTGGTGGTCCCATTTATTT | CTCAACTGGGATCTCGTCAAAG | Quantitative PCR |
| *IFIH1* | CACCATCTGCTTGGGAGAA | CCTGAAGCACGAGATGAGATAG | Quantitative PCR |
| *C1QBP* | TTTGATGGTGAGGAGGAACC | GGGAGTTGATGTCAGTTCAGG | Quantitative PCR |
| *DDX58* | CCATGCTGTTCTTGGGATAGT | GATGAGAGAGAGAGTGTGTGTAAA G | Quantitative PCR |
| *IRF1* | GTGTGGATCTTGCCACATTTC | CCGAGCAAGGCACTGTATAA | Quantitative PCR |
| *APOL1* | GAACAGGTGGAGAGGGTTAATG | GACTACATCCAGCACAAGAAAGA | Quantitative PCR |
| *APOL6* | TGCTGTCAGTTCATTGGTAGAG | CGCTTTGCTAGATCCCTAGATG | Quantitative PCR |
| *PLAAT4* | CAGGCGTTCTCTAGATCCTTTC | GGGCAGATGGCTGTTTATTG | Quantitative PCR |
| *PSMB9* | CATGGGATAGAACTGGAGGAAC | GCAGACAAGTCCTCTCGATATT | Quantitative PCR |
| *TRIM22* | CTCGACCTGCTTATCCGTATTT | CTCAGCACAAGGGCTACTATG | Quantitative PCR |
| *UBA7* | CCACCTACACAAAGACCCTATC | CTTGAATACAGCTCCTCATCCA | Quantitative PCR |
| *UBE2L6* | CCTCAATGTGCTGGTGAATAGA | GGTGAACTCTTCGGCATTCT | Quantitative PCR |
| *STING1* | GCCGAACTCTCTCAATGGTATC | CTCCTCCTCCTCTCCATTCTT | Quantitative PCR |
| *RPS13* | GGTTGAAGTTGACATCTGACGA | CTTGTGCAACACCATGTGAAT | Quantitative PCR |
| *HSPA4* | AGTGTGTCCAGTGCATCTTTAG | TGCATCTTCTCTTCCTCCTTTG | Quantitative PCR |
| Unmethylated L1 loci | TGTGTGTGAGTTGAAGTAGGGT | ACCCAATTTTCCAAATACAACCATC A | LINE1 methylation |
| Methylated L1 loci | CGCGAGTCGAAGTAGGGC | ACCCGATTTTCCAAATACGACCG | LINE1 methylation |

Supplementary Table 17
Antibodies used in the study.

| *Target* | *Manufacturer* | *Catalog No.* | *Application* |
|---|---|---|---|
| STAT1 | Cell Signaling Technology (Danvers, MA) | 9172 | Western blotting |
| p-STAT1 | ABclonal (Woburn, MA) | AP0135 | Western blotting |
| p63α | Cell Signaling Technology | 13109 | Western blotting |
| Histone H3 | Cell Signaling Technology | 4499 | Western blotting |
| LINE1ORF1 | Merck Millipore (Burlington, MA) | MABC1152 | Western blotting |
| STING1 | Cell Signaling Technology | 13647 | Western blotting |
| p-STING1 | Cell Signaling Technology | 50907 | Western blotting |
| p84 | GeneTex (Irvine, CA) | GTX70220 | Western blotting |
| Cas9 | Cell Signaling Technology | 14697 | Western blotting |
| H3K4me1 | Cell Signaling Technology | 5326 | Western blotting |
| H3K4me2 | Cell Signaling Technology | 9725 | Western blotting |
| H3K4me3 | Cell Signaling Technology | 9751 | Western blotting |
| H3K9me1 | Cell Signaling Technology | 14186 | Western blotting |
| H3K9me2 | Cell Signaling Technology | 4658 | Western blotting |
| H3K9me3 | Cell Signaling Technology | 13969 | Western blotting |
| H3K27me3 | Cell Signaling Technology | 9733 | Western blotting |
| H2A | Cell Signaling Technology | 12349 | Western blotting |
| MAVS | Cell Signaling Technology | 3993 | Western blotting |
| IRF1 | Cell Signaling Technology | 8478 | Western blotting |
| pan-Cytokeratin | Thermo Fisher Scientific | MA5-13156 | Immunohistochemical staining |
| Ki67 | Thermo Fisher Scientific | MA5-14520 | Immunohistochemical staining |
| p63 | Abcam (Cambridge, UK) | ab735 | Immunohistochemical staining |
| HLA-ABC | Thermo Fisher Scientific | 11-9983-41 | Flow cytometry |
| EPCAM | BioLegend | 324225 | Flow cytometry |
| Ly6g | BioLegend | 127628 | Flow cytometry |
| Ly6c | BioLegend | 128033 | Flow cytometry |
| Cd11b | BioLegend | 101228 | Flow cytometry |
| Cd45 | BioLegend | 103147 | Flow cytometry |
| mouse I-A/I-E | BioLegend | 107611 | Flow cytometry |
| F4/80 | BioLegend | 123114 | Flow cytometry |
| Cd88 | Thermo Fisher Scientific | 51-0882-80 | Flow cytometry |
| H-2K$^d$ | BioLegend | 116628 | Flow cytometry |
